# Frameshifts may carry oncogenic potential beyond loss of function and categorize genes’ role in tumor development

**DOI:** 10.1101/2022.07.10.499483

**Authors:** Stefan Kirov

## Abstract

In this work I present evidence that frameshift mutations represent substantial oncogenic potential across multiple tumor types and may change our understanding of the function of some genes with well established tumor suppressor. I analyzed data deposited in Cbio portal and show that frameshifts, even when they result in the removal of a substantial part of a protein have the potential to create recurring large domains with unknown function. Based on this analysis I propose a novel categorization of genes according to their association with cancer that is more reflective of a complex nature that goes beyond the simple division to tumor suppressors and oncogenes.

## Introduction

Mutations have been recognized as a driving force in cancer for a long time^1,2^. As modern molecular biology techniques provide a critical mass of genetics and transcriptomics information, the field has gained new level of understanding of the different types of mutations and their impact on tumor development^3^.

Frameshifts have been explored in the context of cancer for a very long time^4^ and to a large extent have been considered mostly as another mechanism to disrupt protein activity in tumor suppressor genes^5,6^. This is not surprising given how well established tumor suppressor genes such as APC, etc. are frequently disturbed by frameshifts in contrast to oncogenes like KRAS that are almost never mutated in such a way^7^.

At the same time, ample research in various species (viruses, prokaryotes, eukaryotes) show that frameshifts may provide a driver for a new phenotype^8,9^. In MSI-H/dMMR cancers there are studies showing increased generation of neoantigens providing evidence that these alternative sequences are indeed translated and potentially functional^10,11^. These findings can fuel next generation cancer vaccines and targeted development of small molecules and ADC^12,13^.

The development of Next Generation Sequencing technologies(NGS)^14^ and the consequential precipitous fall in genome sequencing costs^15^ provides us with access to critical levels of genomics data^16,17^. Based on the vast number of cancer genomes we present evidence that frameshifts are much more likely to have oncogenic potential beyond loss of function.

## Materials and Methods

### Data and code availability

All data was obtained from CBio portal git repository^18^. Scripts used to analyze the data were deposited in Github^19^.

We selected the longest protein form from Ensembl release 106 to determine the representative protein length for a gene. As Cbio portal data is not standardized this means that in some cases this length will be incorrect but we expect these discrepancies to be relatively rare and with minor impact.

The data as analyzed with perl and R scripts.

### Gene set enrichment

Gene set enrichment analysis was done using Webgestalt^20^ using gene symbols as input, protein coding genes as background set and selecting non redundant categories.

## Results

Out of all patients for which Cbio portal stores mutation data, 41232 patients carried at least 1 frameshift mutation. Of those 5456 or 13% harbored at least 1 large frameshift mutation, which created an alternative C-end of the protein of more than 100 amino acids. In 15841 or 38% of the patients at least 1 frameshift occurs closely to the end of the protein, removing less than 20% of the original sequence. The union of these categories amounts to events in 17841 or 43% of all patients with a frameshift mutation.

I summarized the data with respect to the length of the alternative C-end fragment generated (Figure 1). There is an expected bias towards short alternative fragments (<10 aa) with mean length of 29 aa, however there is a tail with a maximal value of 947 aa which is close to the average protein length based on ensembl release 106 of 1278 aa. This distribution supports the hypothesis that most frameshifts are truncating in nature and abolish completely or partially the function of the genes. However this explanation is unlikely for the frameshifts with length of the alternative end that often exceeds many well annotated proteins.

**Figure 1.**
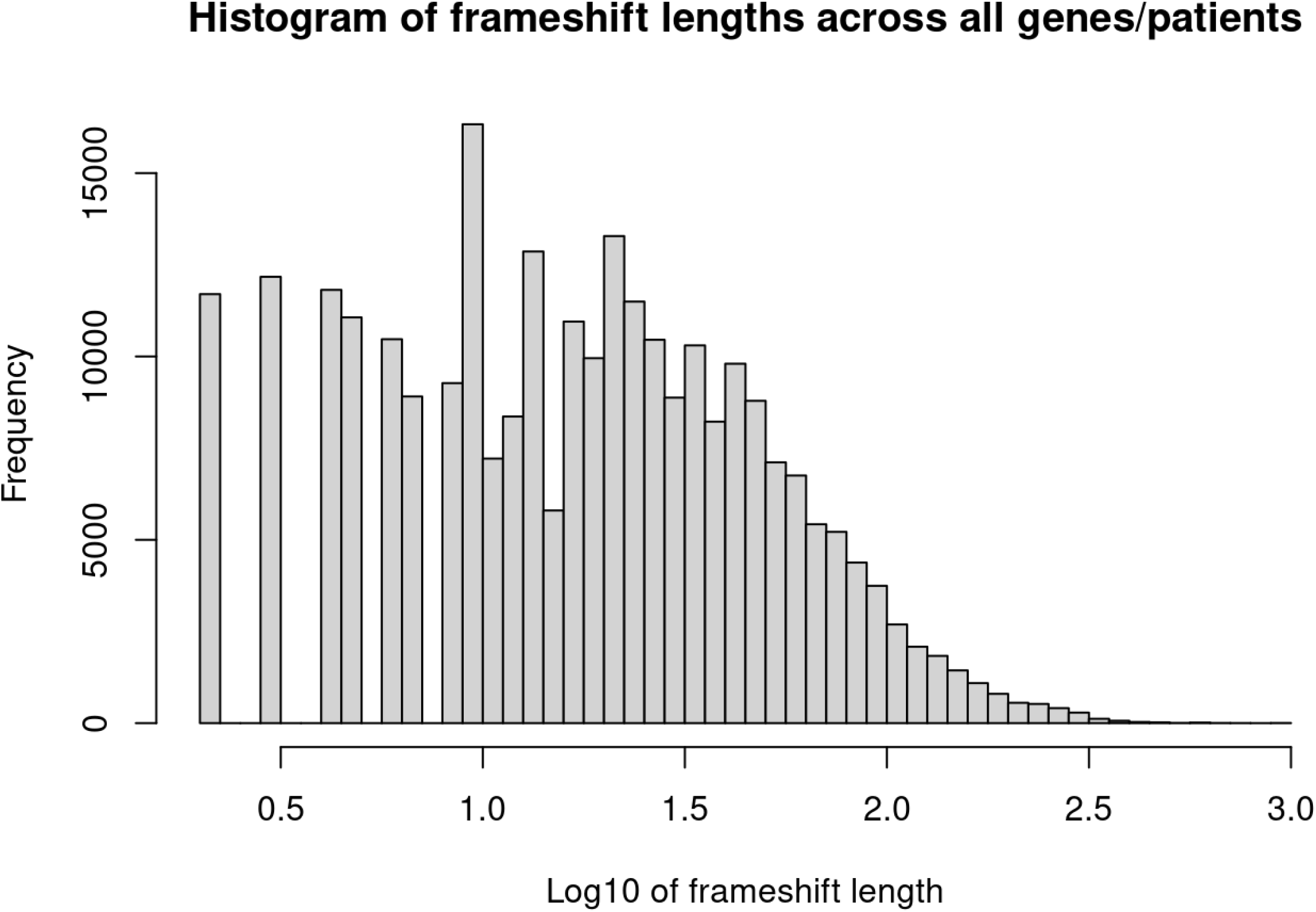
Frequency of the length of the terminal C-end created by a frameshift in amino acid coordinates

Another important factor to consider is the distance of a frameshift from the normal stop codon (Figure 2). Quite surprisingly there is no enrichment for frameshifts to begin at the start of a protein if all frameshifts are truly loss of function. Instead we observe a uniform distribution that is probably representative of more complex mix of consequences. This distribution is quite different in some groups of genes, for example in RARA and BRD4 frameshifts are substantially enriched at the end of the protein (Supplementary information) which could be indicative of gain of function events. In other genes such as IDH1, PLEKHA6, NF1 and RB1 the framshifts start predominantly very early in the protein sequence as expected in a loss of function scenario. We have a third class of genes where there is a local enrichment of frameshift start that is in the middle of the protein, for example APC. Fourth category of genes exhibit bimodal enrichment, both at the start and end of the protein with an extreme example of STK11. Finally, a number of genes have a dispersed pattern, surprisingly TP53 and VHL among them. Summarized proposal to categorize genes and their association with cancer is given in Table 1.

**Figure 2.**
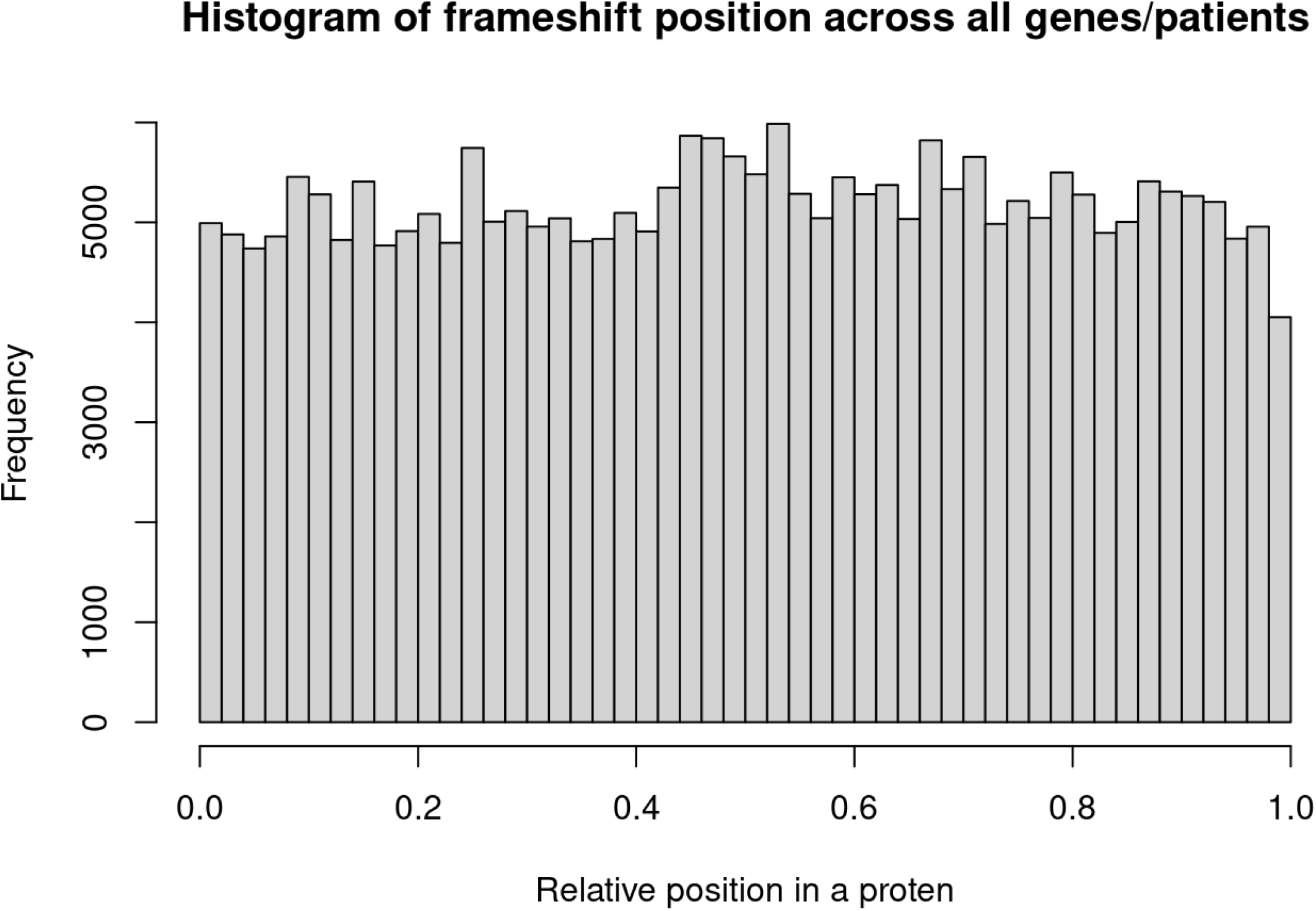
Normalized position of the frameshift start across all genes

**Table 1.** Putative category based on specific features

| Category | Description | Examples |  |
| --- | --- | --- | --- |
| Loss of function | Short frameshifts at the start of the gene | IDH1, PLEKHA6, NF1 and RB1 | LOF |
| Gain of function | Frameshifts enriched at the end of the gene | BRD4, RARA | GOF |
| Change of function | Longer frameshifts enriched in the middle of a gene |  | COF |
| Loss or gain of function | Enrichment at the start and end of the gene | STK11 | L/GOF |
| Complex frameshifts | Uniform distribution may need additional features to deconvolute the effect | TP53, VHL | undetermined |

Another important consideration is the reuse of the same frameshift. In any protein position there are 2 possible novel ORFs to use. In a number of genes there is a strong bias to reuse specific ORF. As an example, in SOX9 (Figure 3A), RUNX1(Figure 3B), ARID1A (Figure 3C) and TP53 (Figure 3D) same end is re-used which indicates that the same ORF is being translated as a consequence from the frameshifts. This observation can be explained by a potential change of function, or a need to evade the alternative ORF. In the case of ARID1A the first explanation is more plausible as the length of the alternative C-ends is considerable (>100aa), where with TP53 frameshifts are short <100aa and while still may have an alternative function this is less likely.

**Figure 3.**
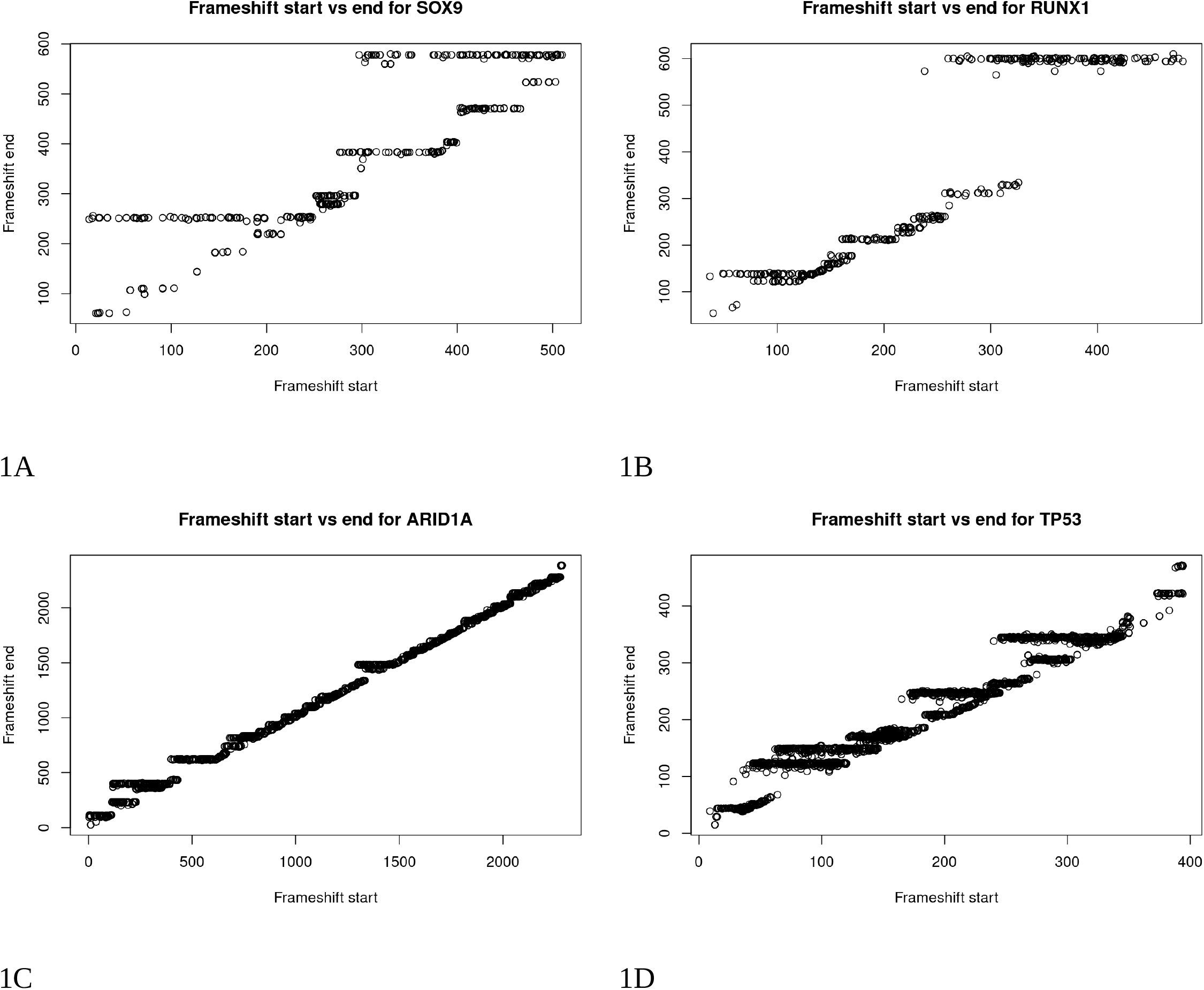
Frameshift position with respect to the calculated novel C-end coordinates

Exploring specific ORFs, most are in well established tumor suppressor genes, sucha s TP53, GATA3 and APC (Supplementary information), but some are consistent with gain of function, for example frameshifts at the end of NPM1 and RNF43. NPM1 frameshifts are restricted to AML and MDS, whereas RNF43 are originating from GI cancers and to a smaller degree from prostate cancer. NPM1 recurrent C-end is only 12 aa, whereas RNF43 is more variable and much larger with mean length of 42 aa.

The ratio between short and long frameshifts observed for any given gene can be used to predict canonical tumor suppressors with the caveat that this overall statistics may be masking the more complex interaction between frameshift length and position distribution (Table 2, Supplementary information). Similarly, we can identify genes where long frameshifts are favored (Table 3, Supplementary information). We can assume these are mostly change of gain of function genes with the same stipulation as with tumor suppressors. These genes are substantially enriched for several functional categories: chromatin remodeling, gene silencing, etc (Table 4, Figure 4). Reactome pathways containing Notch gene family members are strongly enriched (Supplementary information). Additional pathways with significant enrichment include RUNX1 transcriptional program and epigenetic modulation. I did not observe significant chromosome location enrichment that could be an indication of a technical confounding factor.

**Table 2.**
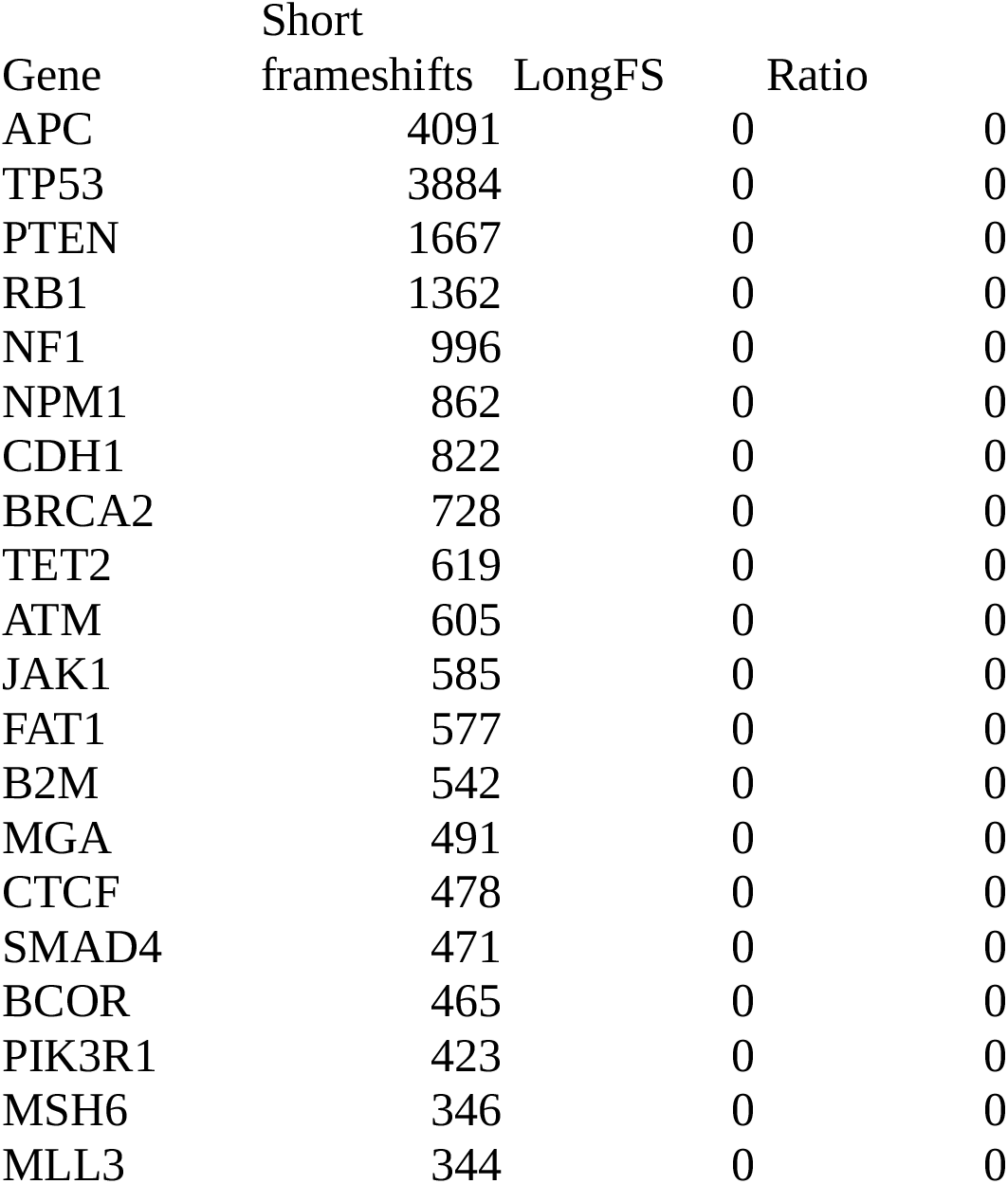
Top 20 tumor suppressor genes as defined by the ratio of long and short frameshifts, sorted by all frameshifts

**Table 3.** Top 20 change/gain of function genes as defined by the ratio of long and short frameshifts, sorted by all frameshifts

| Gene | AllFS | LongFS | Ratio |
| --- | --- | --- | --- |
| AXIN1 | 66 | 84 | 1.27 |
| ZFP36L2 | 71 | 80 | 1.13 |
| BRD4 | 83 | 79 | 0.95 |
| RUNX1 | 354 | 239 | 0.68 |
| NOTCH3 | 147 | 76 | 0.52 |
| SOX9 | 401 | 170 | 0.42 |
| CEBPA | 60 | 25 | 0.42 |
| JAK3 | 53 | 22 | 0.42 |
| NKX2-1 | 64 | 26 | 0.41 |
| TBX3 | 334 | 131 | 0.39 |
| FLG | 61 | 19 | 0.31 |
| NOTCH1 | 454 | 139 | 0.31 |
| EPHA2 | 74 | 20 | 0.27 |
| INPPL1 | 285 | 75 | 0.26 |
| ELF3 | 234 | 59 | 0.25 |
| SMARCA4 | 196 | 46 | 0.23 |
| GPS2 | 87 | 20 | 0.23 |
| RBM10 | 231 | 49 | 0.21 |
| DOT1L | 85 | 18 | 0.21 |
| SPEG | 68 | 14 | 0.21 |

**Table 4.**
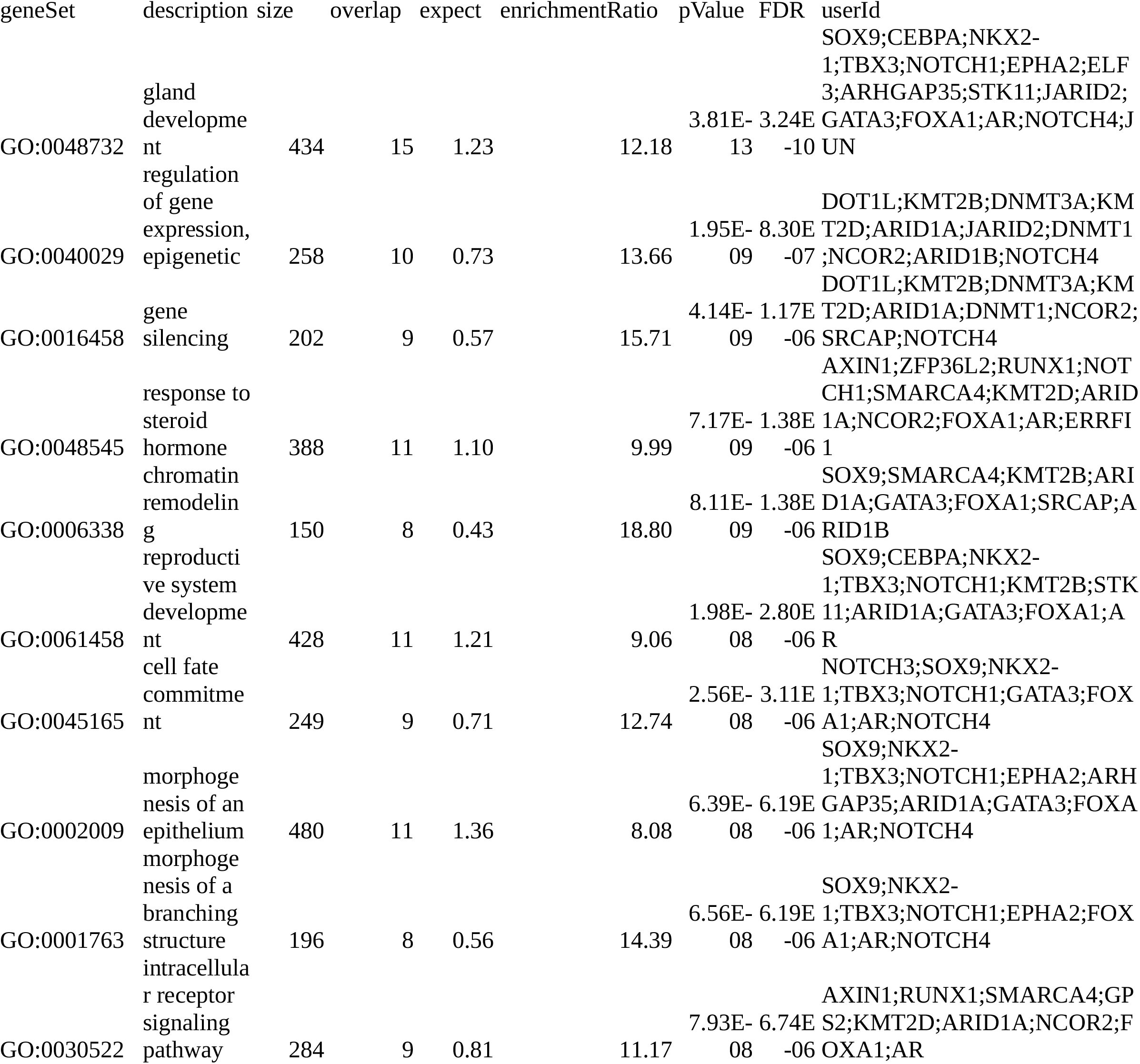
Gene set enrichment analysis of genes with >0.1 ratio of long vs all mutations and at least 50 frameshifts observed in the cohort.

**Figure 4.**
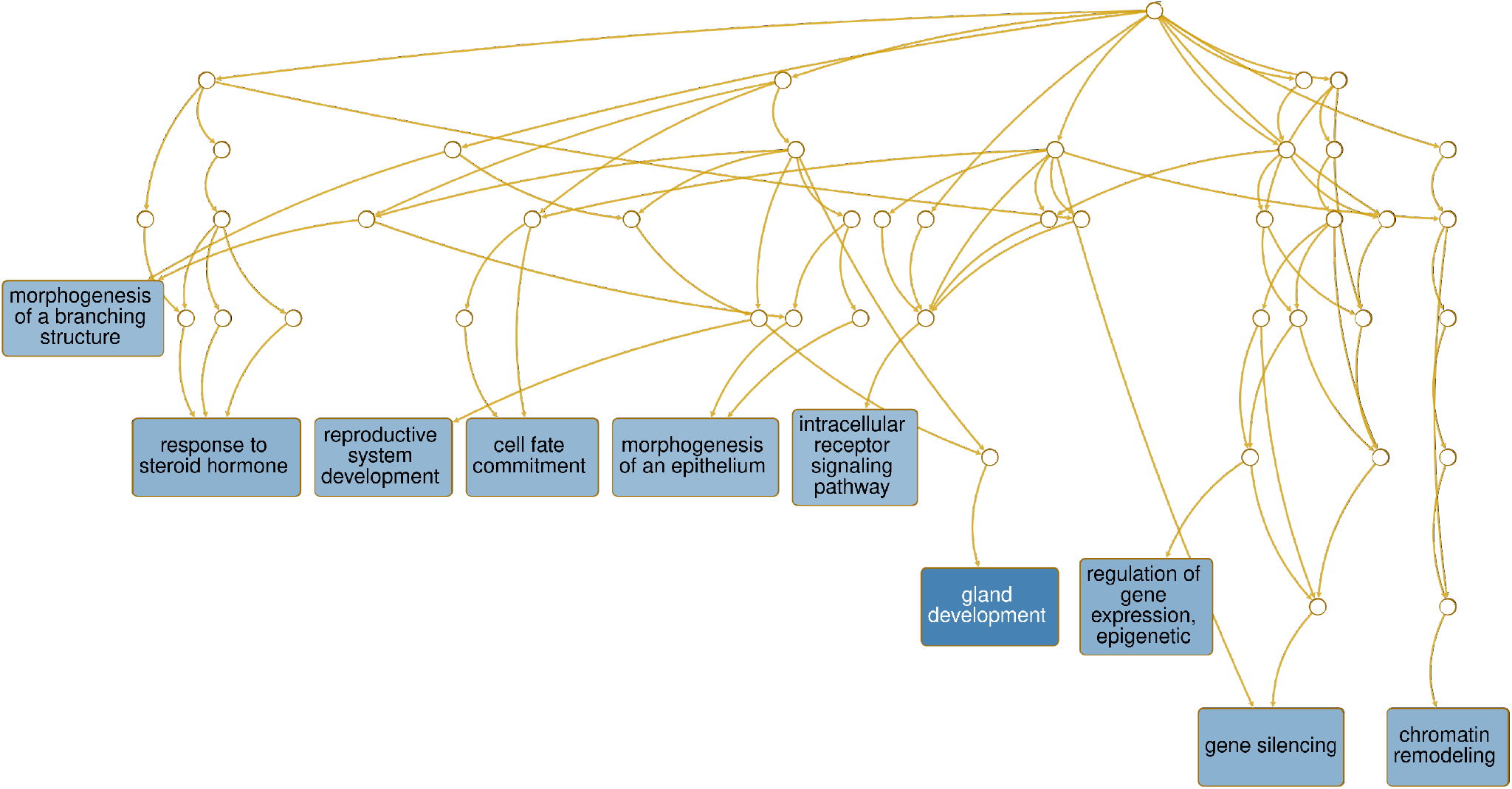
Direct acyclic graph of the enrichment results for the change/gain of function dataset

## Discussion

The role of frameshifts in cancer is a challenging question and have been explored mostly in the context of MSI-high^11^. At the same time frameshifts are thought to play considerable role in ccRCC^10^, however this tumor type has very low frequency of MSI-high tumors^21^. Based on the analysis I present here, it seems plausible that frameshifts play important role in cancer, in a much larger population of patients.

The fact that there is a significant number of shared peptide fragments originating from common shared ORFs is appealing in the context of drug development beyond the scope of the proposed immuno-oncology focus already proposed by Roudko and co-authors^11^. Indeed, if NPM1 and RNF43 are gain of function recurring alternative and cancer specific C-ends, these can be targeted by variety of modalities. NPM1 specifically is an appealing target in AML and MDS where its importance is well recognized^22^, however there is still no active development targeting the cancer specific fragment. That is not the case with RNF43 where phase I trial was initiated and eventually terminated due to lack of financial support^23,24^. Another trial focused on peptide vaccine derived from the native protein and was completed in 2015 but results were not published^25^. None of these drug development programs focused on the cancer specific C-terminal fragment which is arguably a better therapeutic target.

One catgory of recurrent frameshifts could be of particular interest-mutations that give rise to substantial novel protein sequence in both genes with well described association with cancer, such as STK11 and RARA, but also novel or not well established genes, for example DOCK3 and XYLT2. STK11 is particularly interesting examples as it is not only well known tumor suppressor, but has also been associated with resistance to nivolumab treatment^26^. It appears from this analysis that it can have unexpected oncogenic properties. Quite importantly, these mutations apear to be identical to STK11 germ line mutation that is a risk factor for Peutz-Jeghers syndrome^27^, which is associated with 20 fold increase in cancer risk. In fact, the involvement of STK11 in this syndrome was used to define the gene as a tumor suppressor gene. However, given the recurrent frameshift towards the end of the protein raises the possibility that simple loss of function similar to the frameshifts close to the N-end is not a sufficient explanation.

RUNX1 presents another interesting case where half of the protein is substituted by a novel sequence that is aproximately twice the size of the lost fragment. This frameshift occurs mostly in AML and MDS patients, however it is observed in solid tumors as well. This could be an enticing drug development as the fragment size is very large and may be easier to develop specific small molecule against it. Similarly, an already appealing target in cancer, RARA, has been extensively studied because of fusions in leukemia^28^. The data I present here shows this could be further extended to frameshifts in a number of solid tumors.

With the substantial increase in sequenced cancer genomes it is becoming increasingly clear that the current categorization of genes as tumor suppressors or oncogenes is not sufficient to reflect the true complexity of oncogenesis^29^. There are many examples of genes which defy this simple categorization and are known or suspected to act as both tumor supressors or oncogenes^30–32^. Even perhaps the most widely known tumor suppresor gene TP53 is now thought to have oncogenic properties in certain cases^33,34^. Another example of how this binary definition is not sufficiently adequate is the R132 mutation in IDH1^35^. While the mutation leads to an oncogenic transformation through a 2-hydroxyglutarate production it is somewhat misleading to categorize this even as a gain of function compared for example to the hyperactivation of EGFR and other RTKs^36^.

## Acknowledgments

“The results <published or shown> here are in whole or part based upon data generated by the TCGA Research Network: https://www.cancer.gov/tcga.”

