## Supplementary Information for "Frameshifts may carry oncogenic potential beyond loss of function and categorize genes’ role in tumor development"

### FrameshiftsInCancer

Stefan Kirov

2022-07-07

### R Load/prep data

### General stats

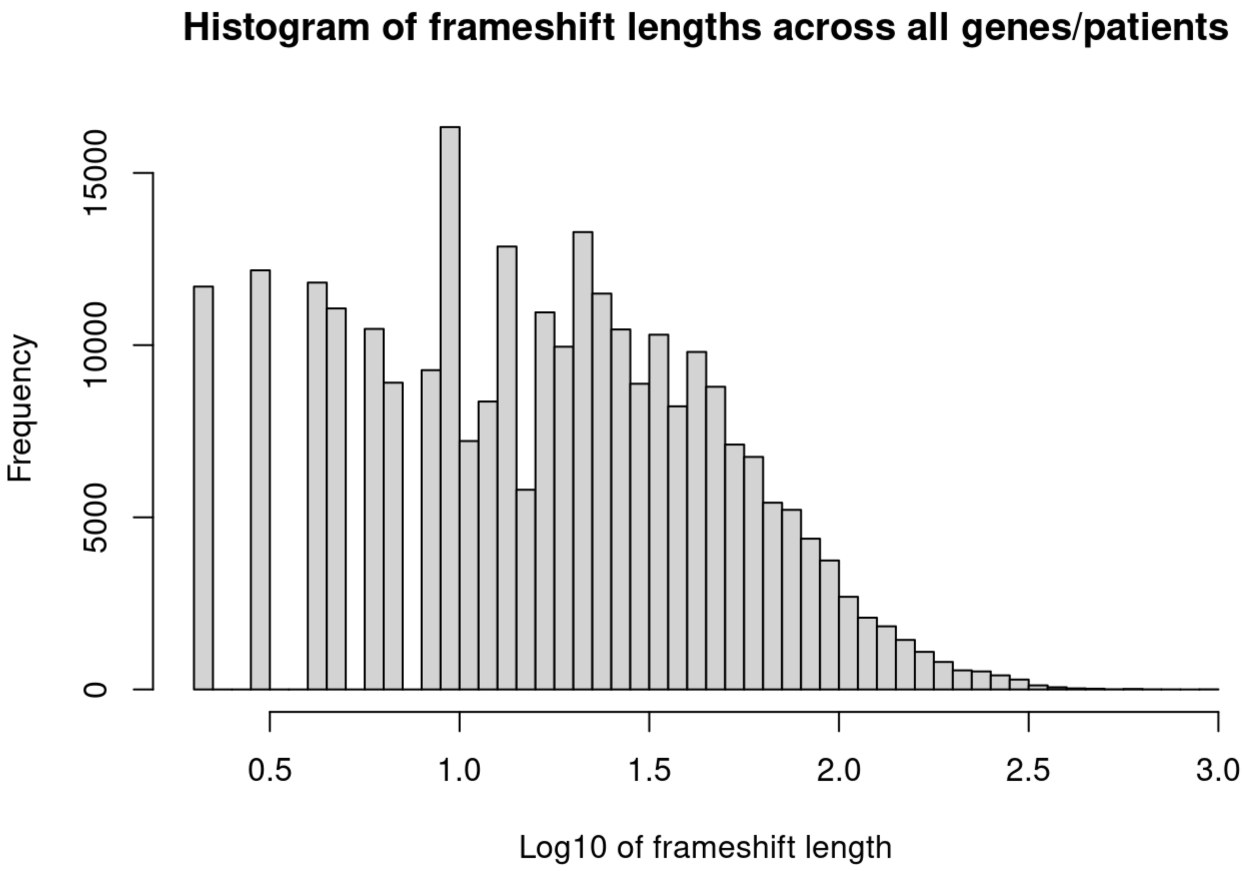

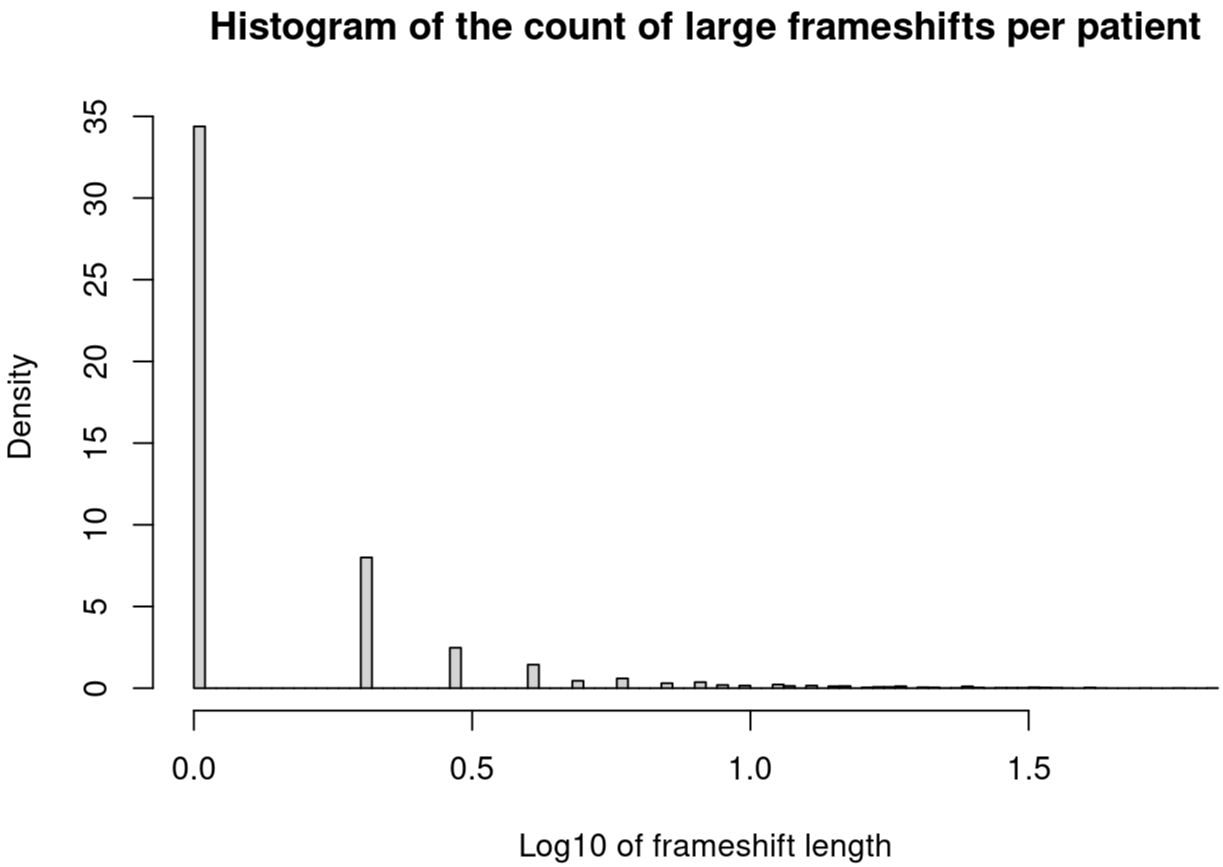
Histogram of the count of large frameshifts per patient
Density
0
5
10
15
20
25
30
35
0.0
0.5
1.0
1.5
Log10 of frameshift length

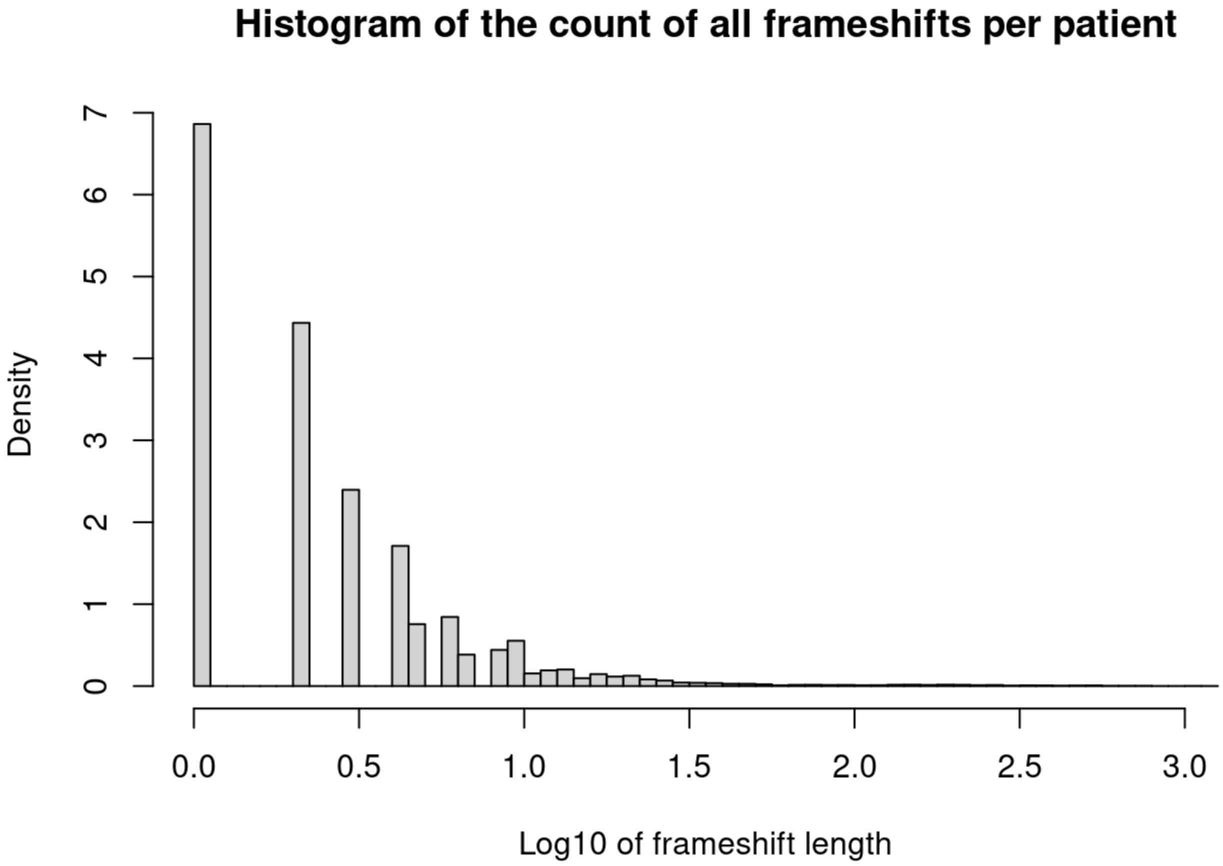
Histogram of the count of all frameshifts per patient
Density
0
1
2
3
4
5
6
7
0.0
0.5
1.0
1.5
2.0
2.5
3.0
Log10 of frameshift length

Note that the `echo = FALSE` parameter was added to the code chunk to prevent printing of the R code that generated the plot.

### Advanced analysis

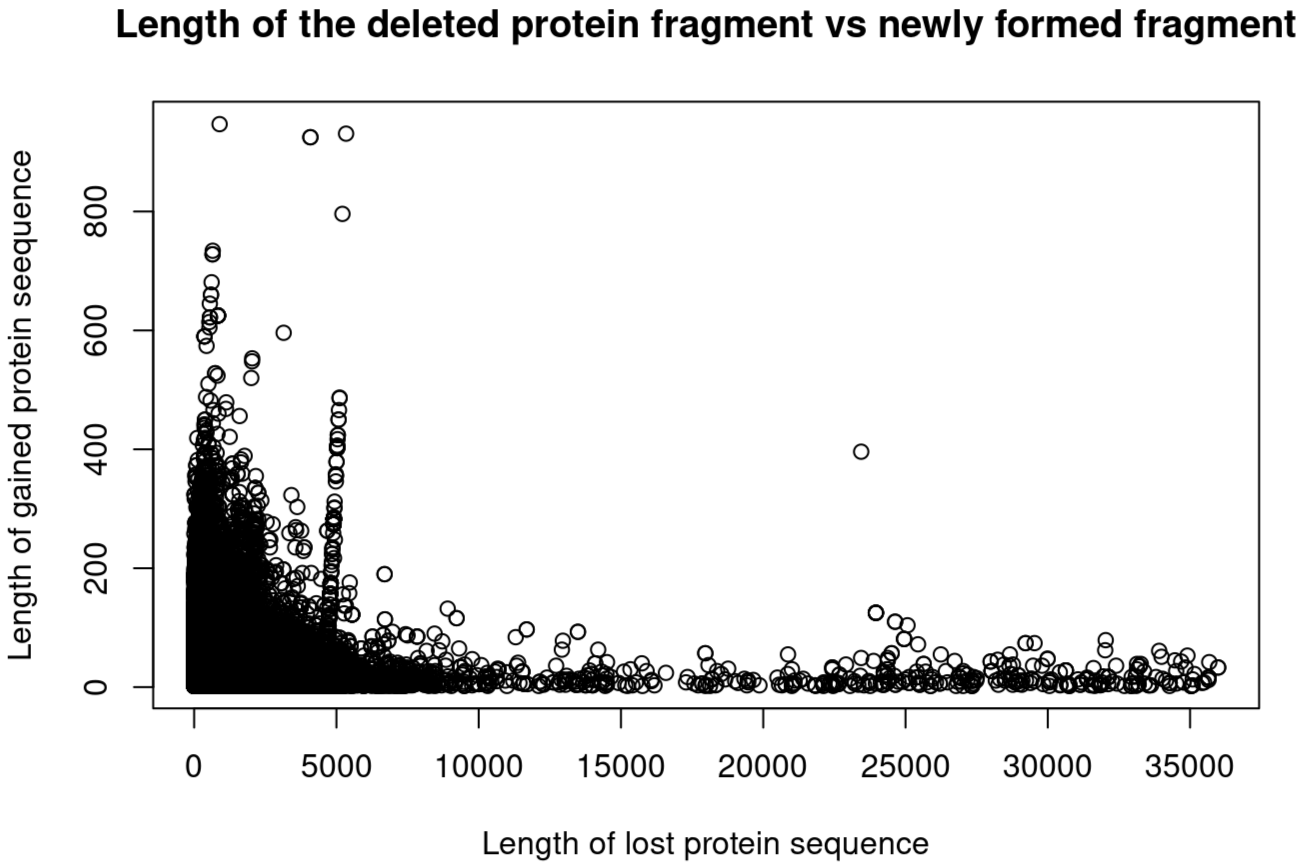

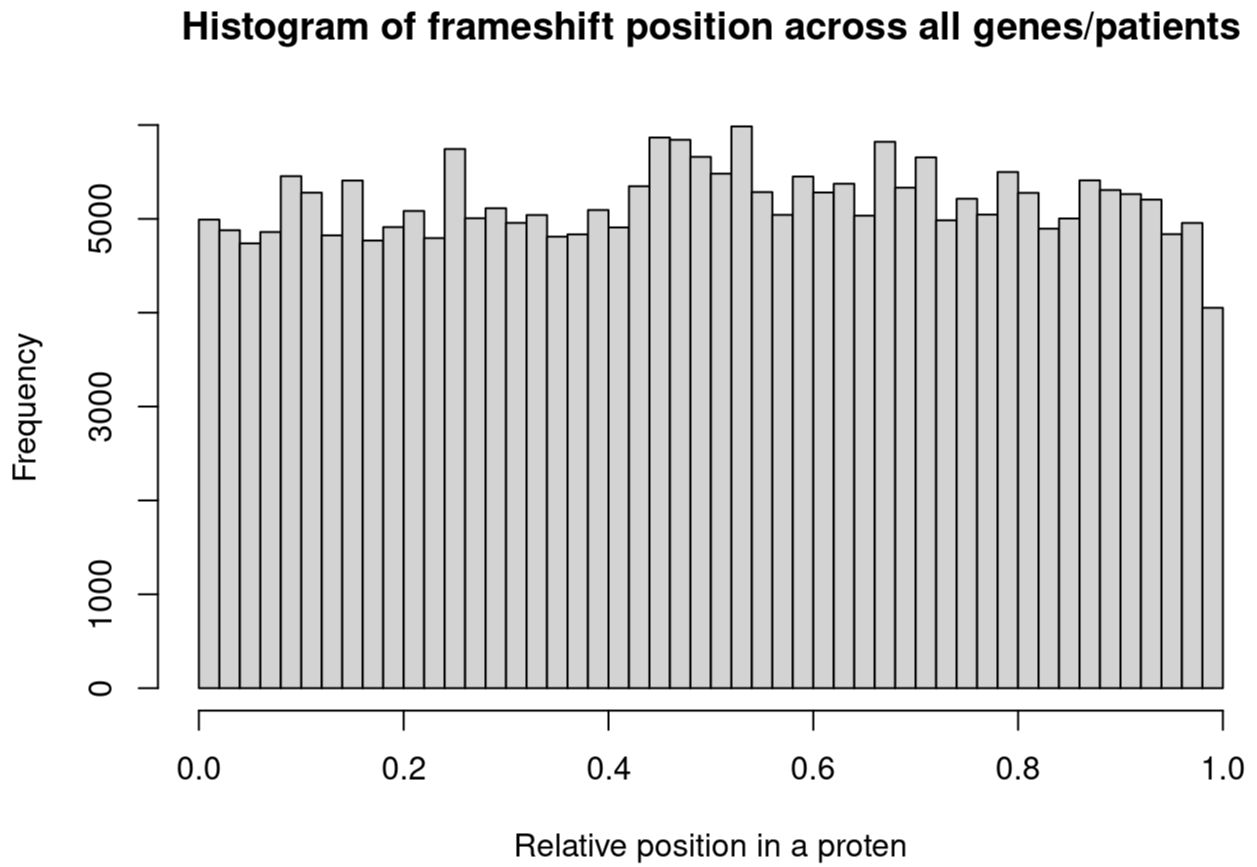


```
## [1] 15841
```

```
## [1] 17841
```

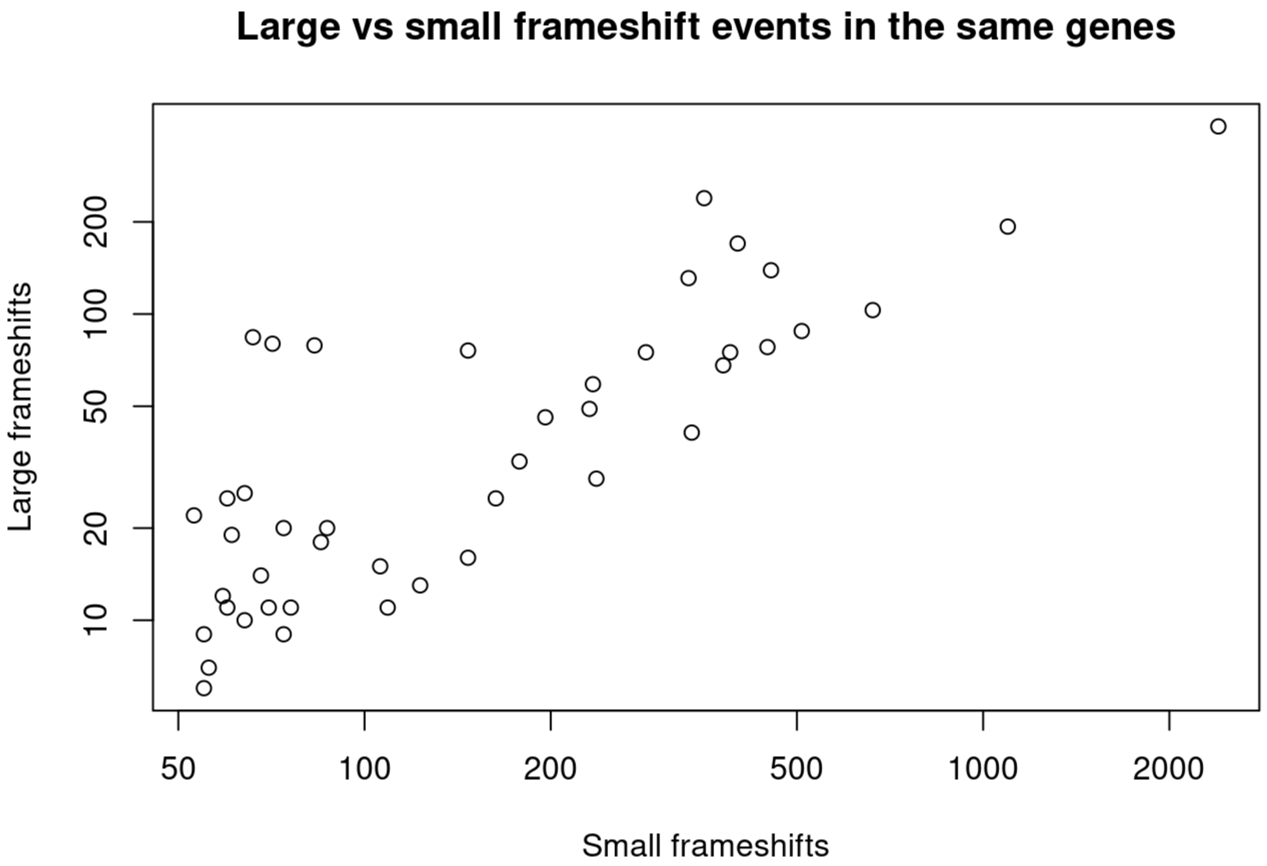
Large vs small frameshift events in the same genes
Large frameshifts
Small frameshifts
10
20
50
100
200
50
100
200
500
1000
2000

```
## [1] 0.8748147
```

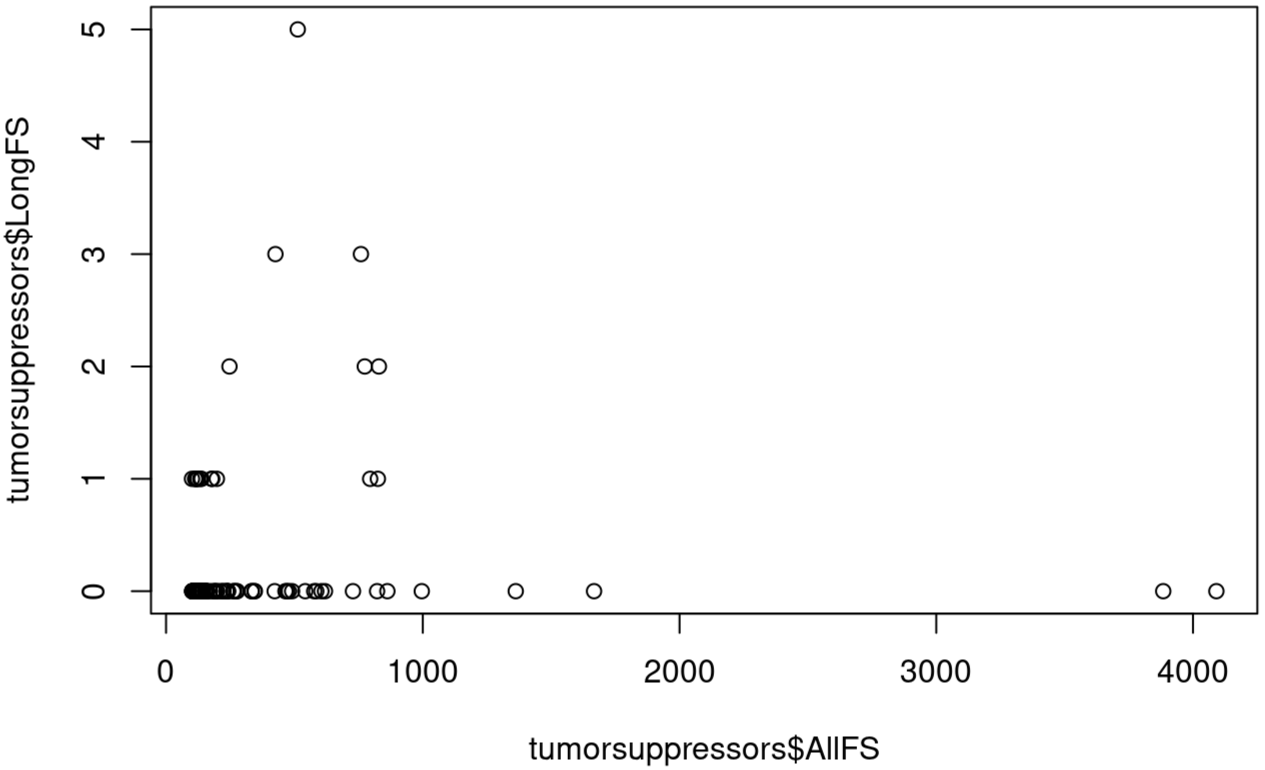


```
## [1] 0.04443743
```

### Gene stats

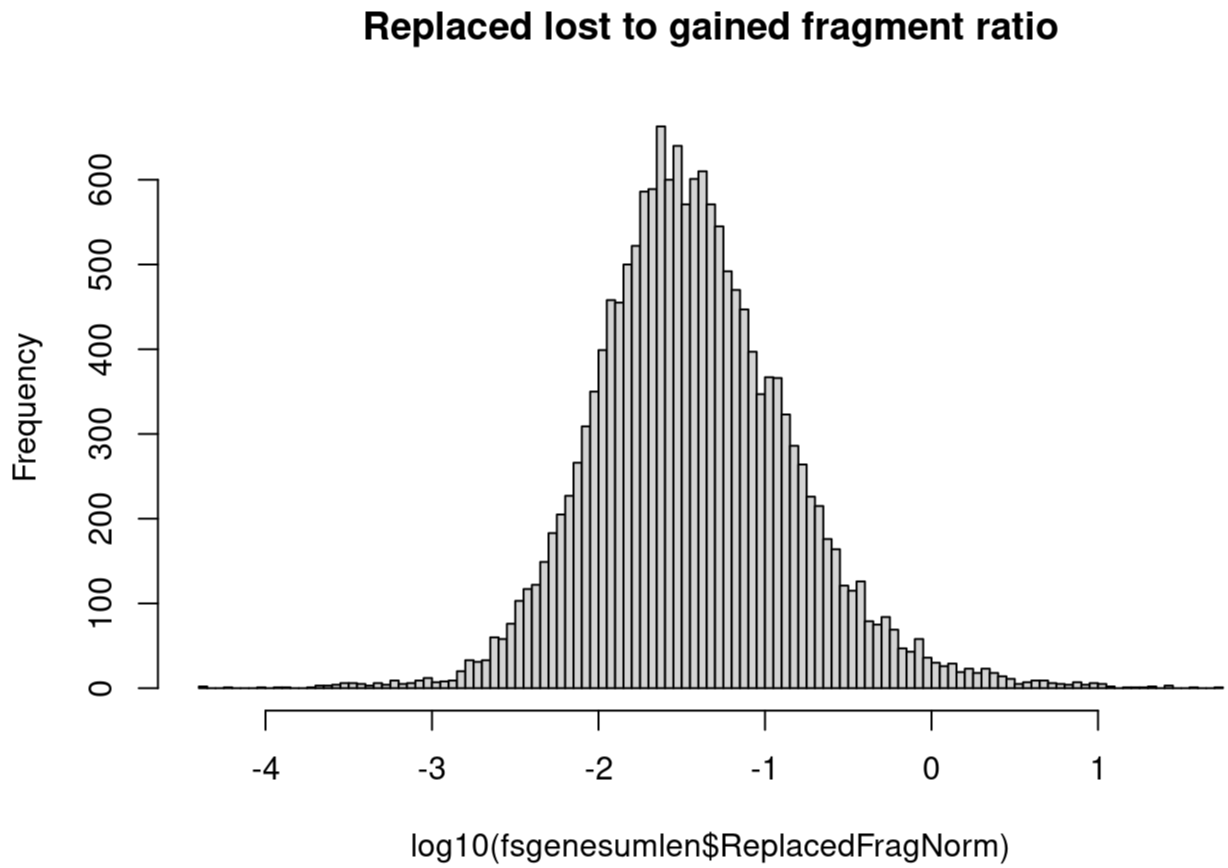


### Examples of TS and others

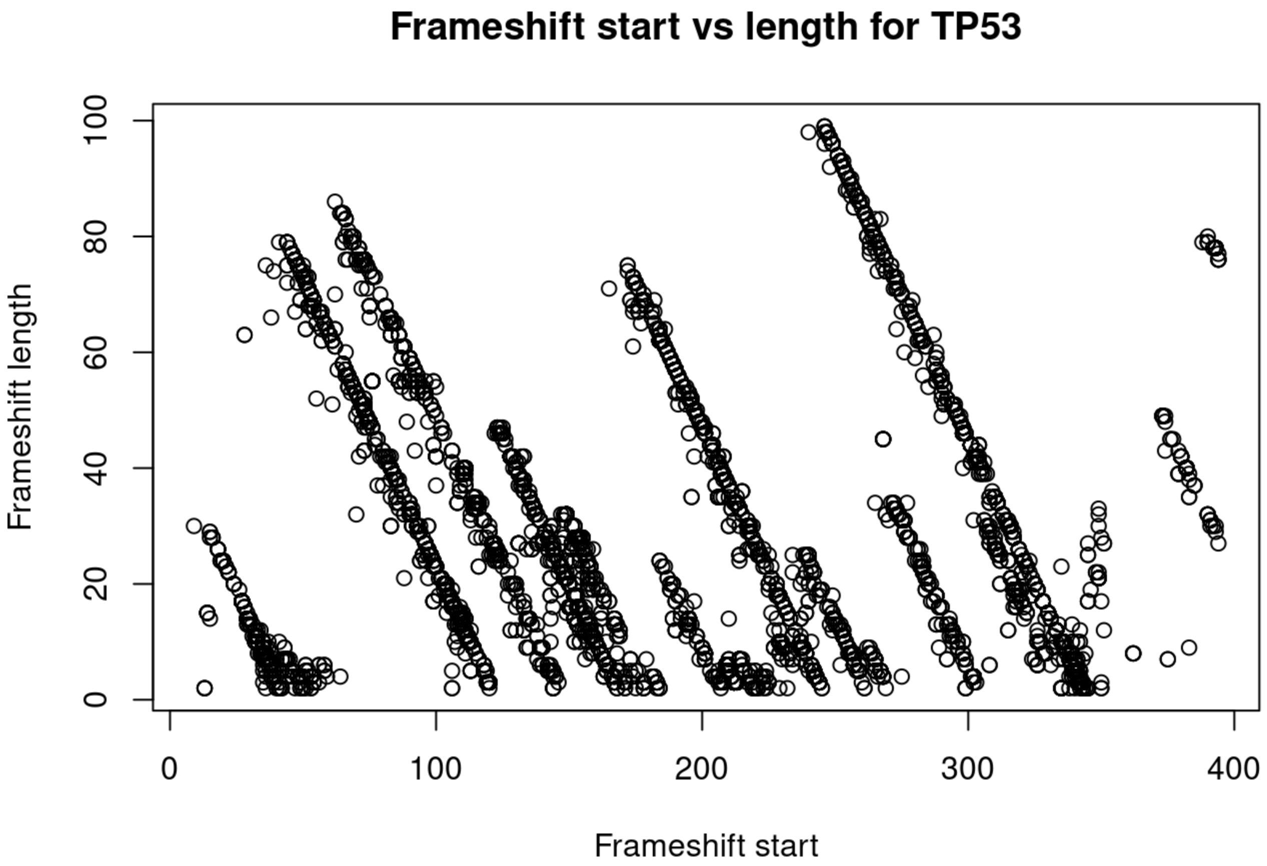
Frameshift start vs length for TP53
Frameshift length
Frameshift start
0
20
40
60
80
100
0
100
200
300
400

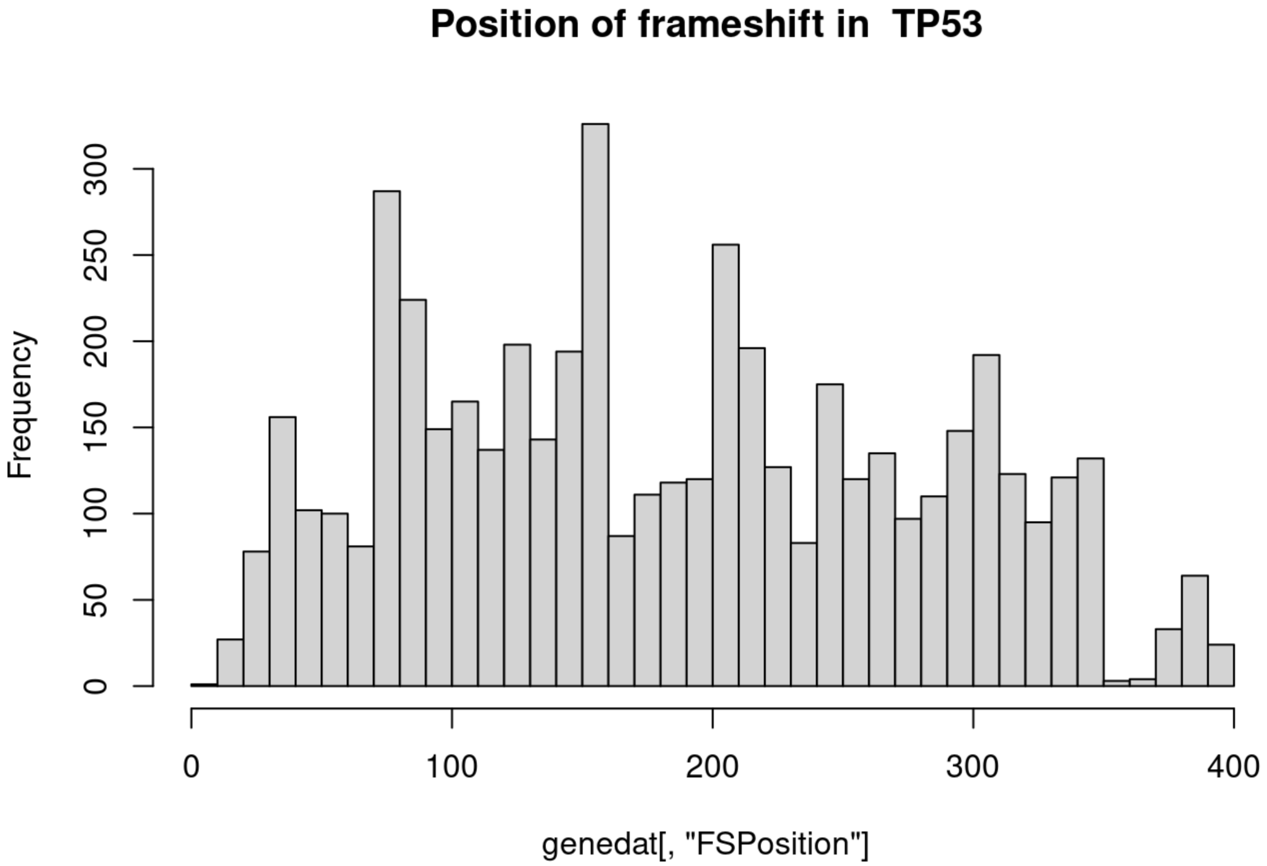
Position of frameshift in TP53
Frequency
genedat[, "FSPosition"]
0
50
100
150
200
250
300
0
100
200
300
400

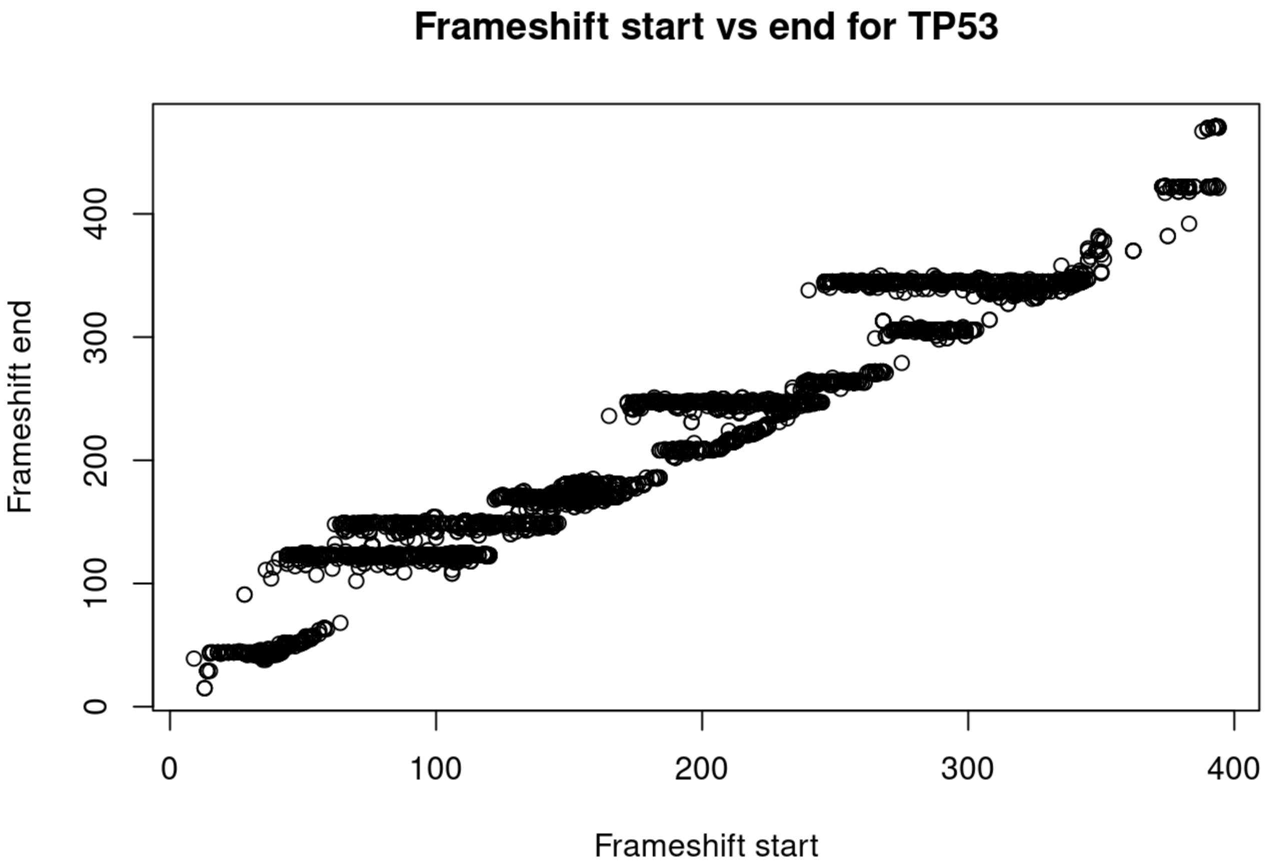
Frameshift start vs end for TP53
Frameshift end
0
100
200
300
400
Frameshift start
0
100
200
300
400

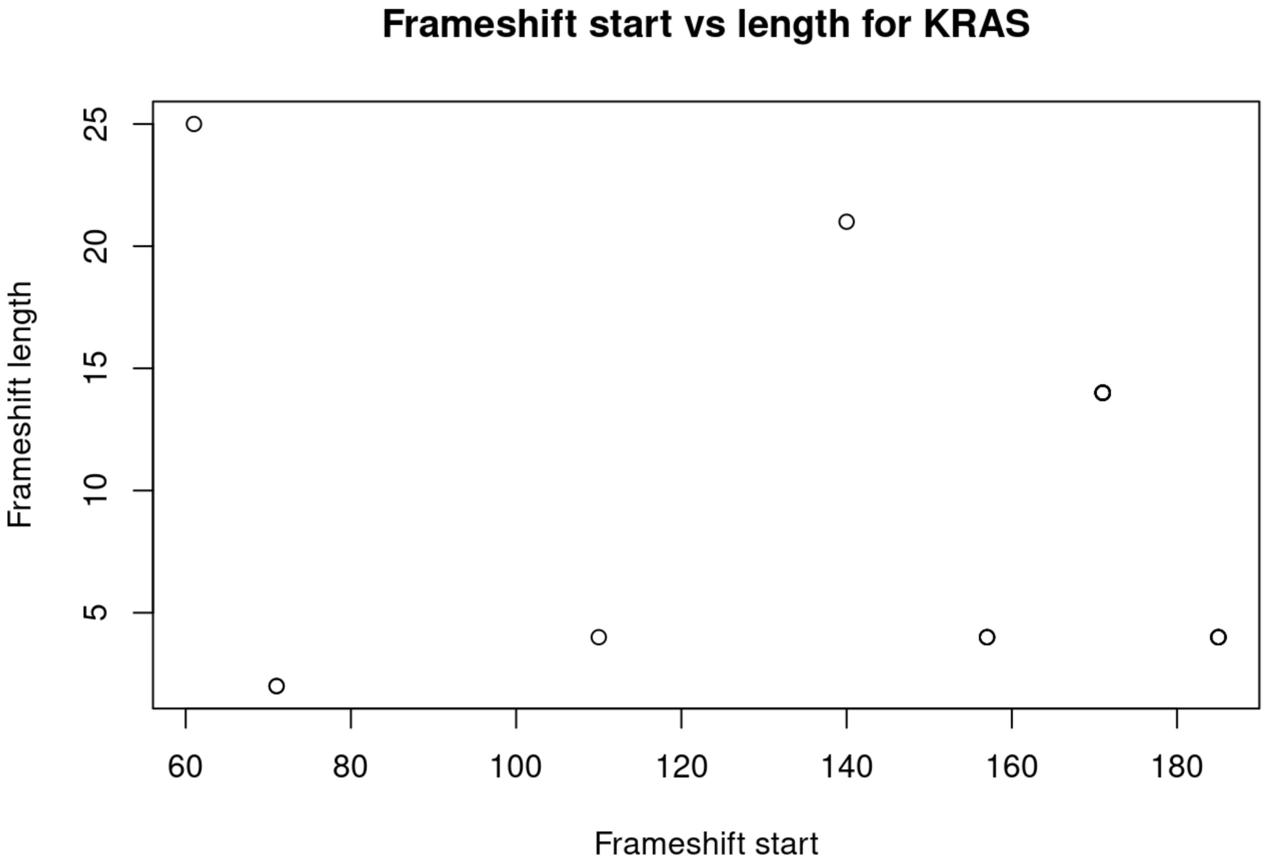
Frameshift start vs length for KRAS
Frameshift length
5
10
15
20
25
Frameshift start
60
80
100
120
140
160
180

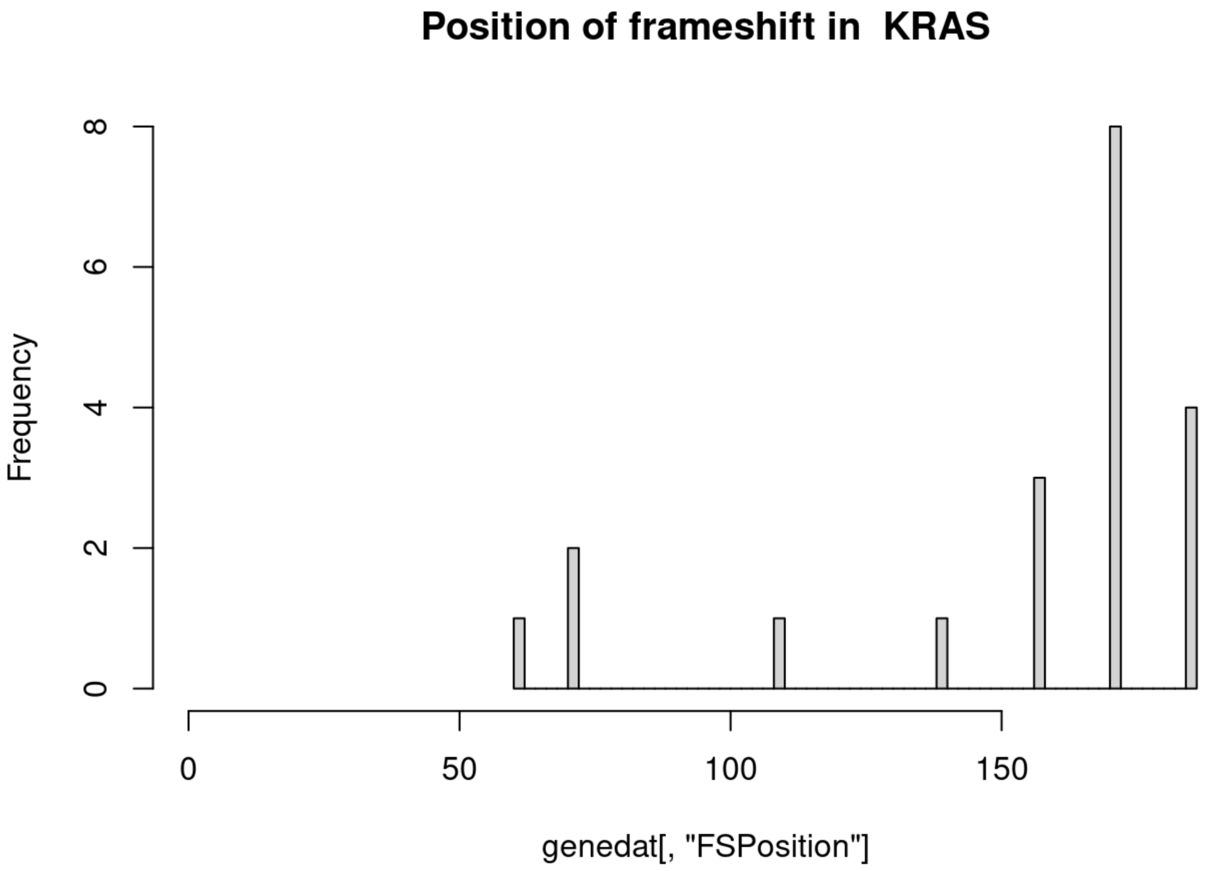
Position of frameshift in KRAS
Frequency
0
2
4
6
8
0
50
100
150
genedat[, "FSPosition"]

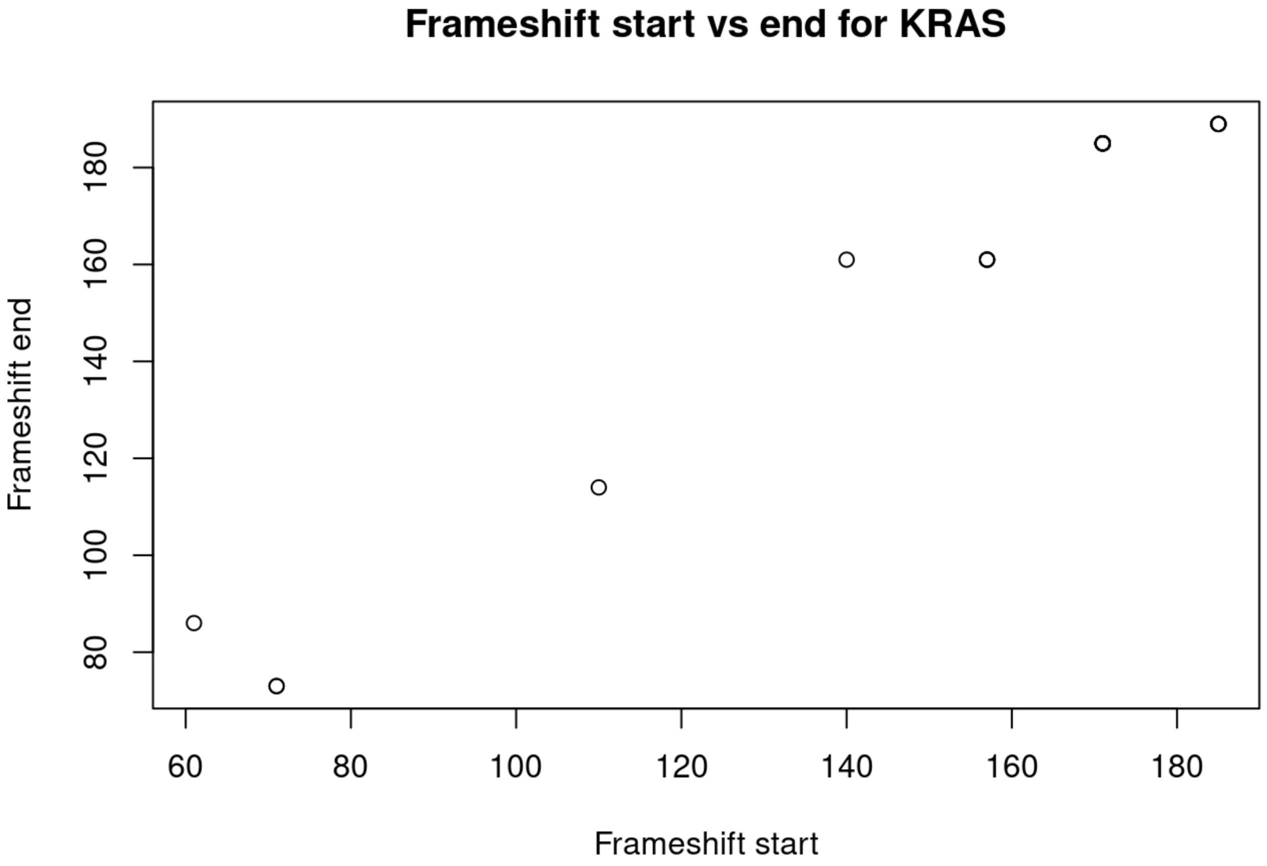
Frameshift start vs end for KRAS
Frameshift end
80
100
120
140
160
180
60
80
100
120
140
160
180
Frameshift start

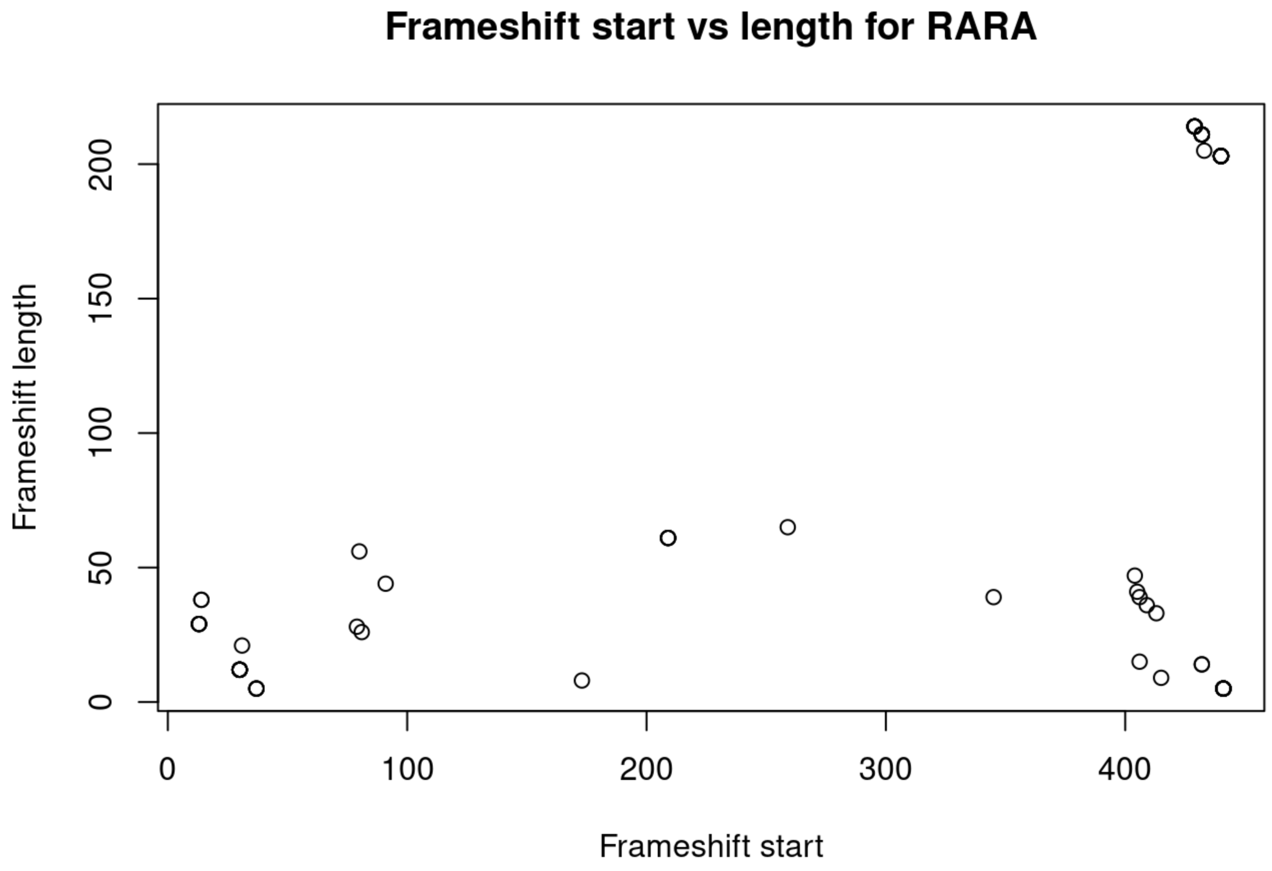
Frameshift start vs length for RARA
Frameshift length
Frameshift start
0
50
100
150
200
0
100
200
300
400

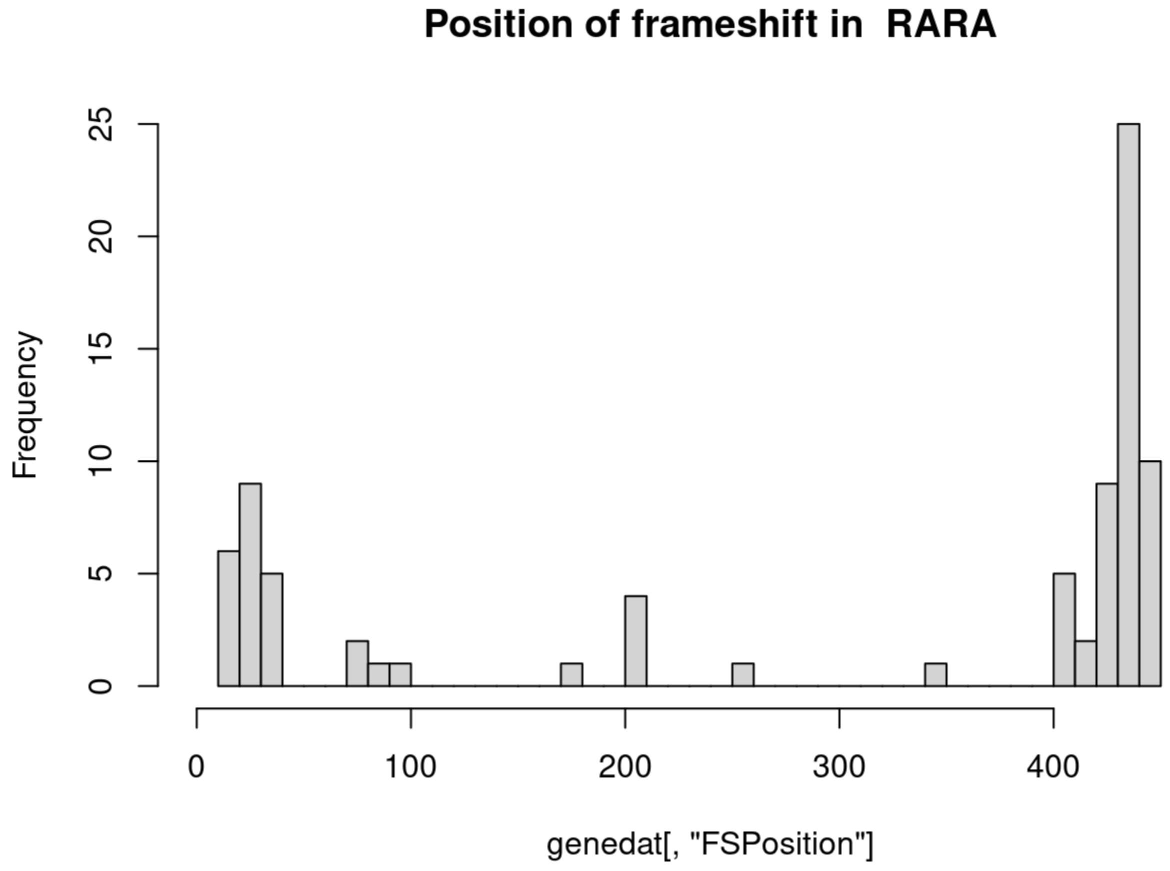
Position of frameshift in RARA
Frequency
genedat[, "FSPosition"]
0
5
10
15
20
25
0
100
200
300
400

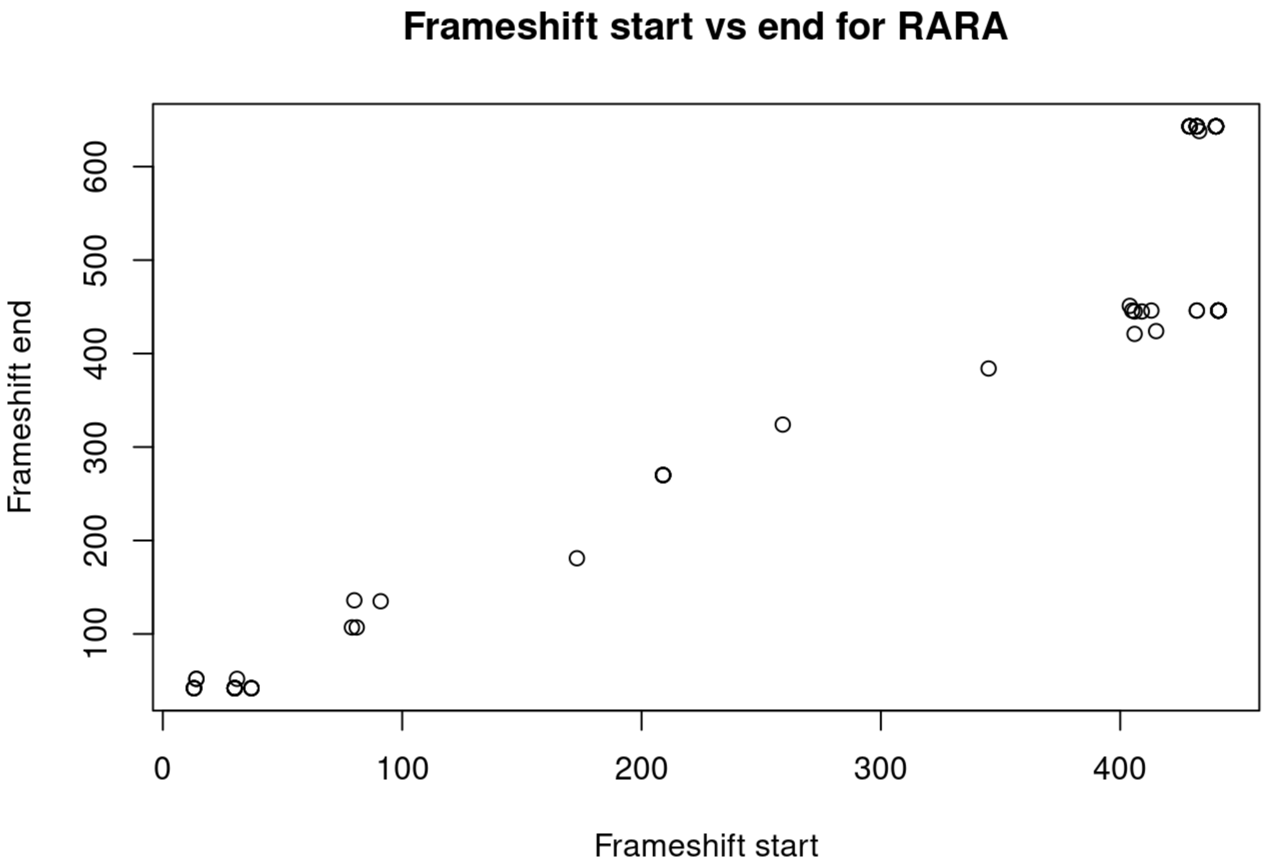
Frameshift start vs end for RARA
Frameshift end
Frameshift start
0
100
200
300
400
500
600

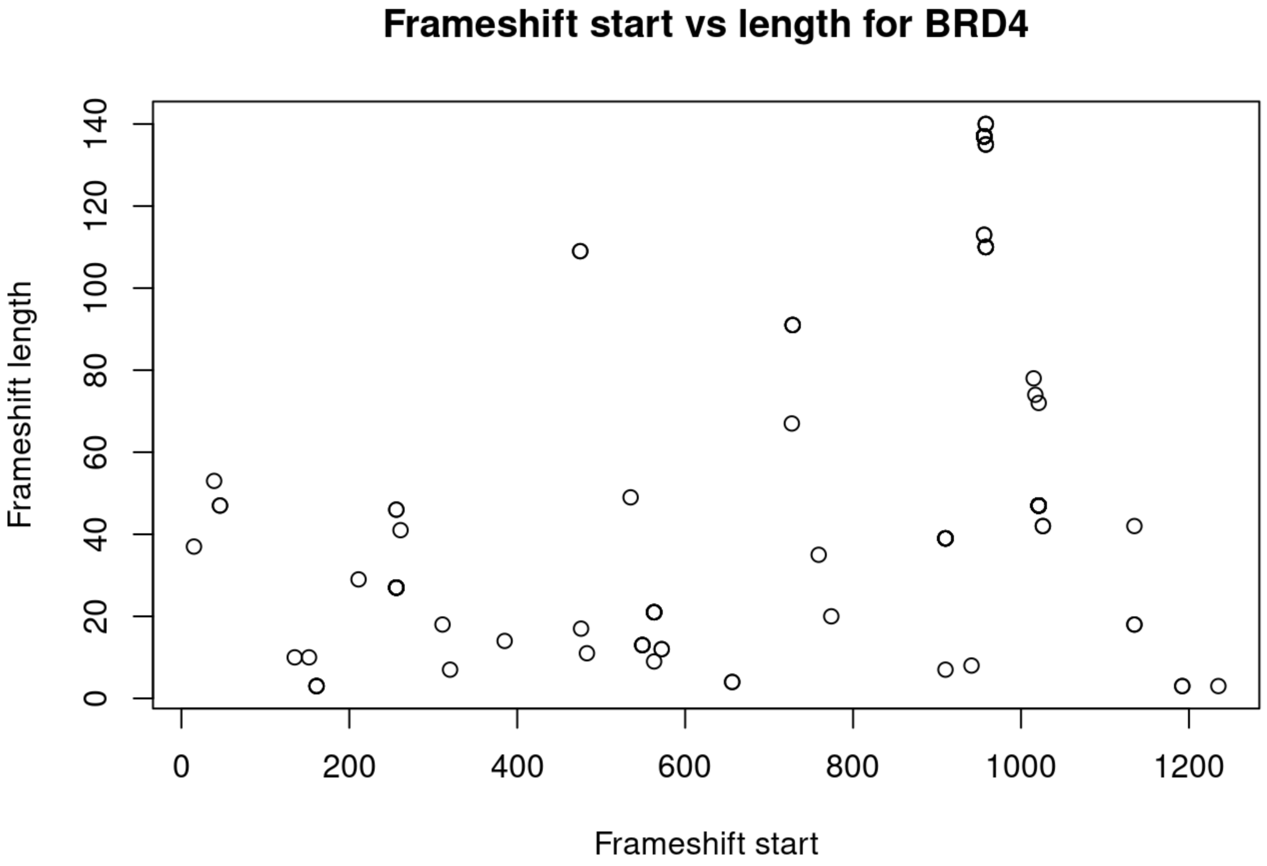
Frameshift start vs length for BRD4
Frameshift length
Frameshift start
0
20
40
60
80
100
120
140
200
400
600
800
1000
1200

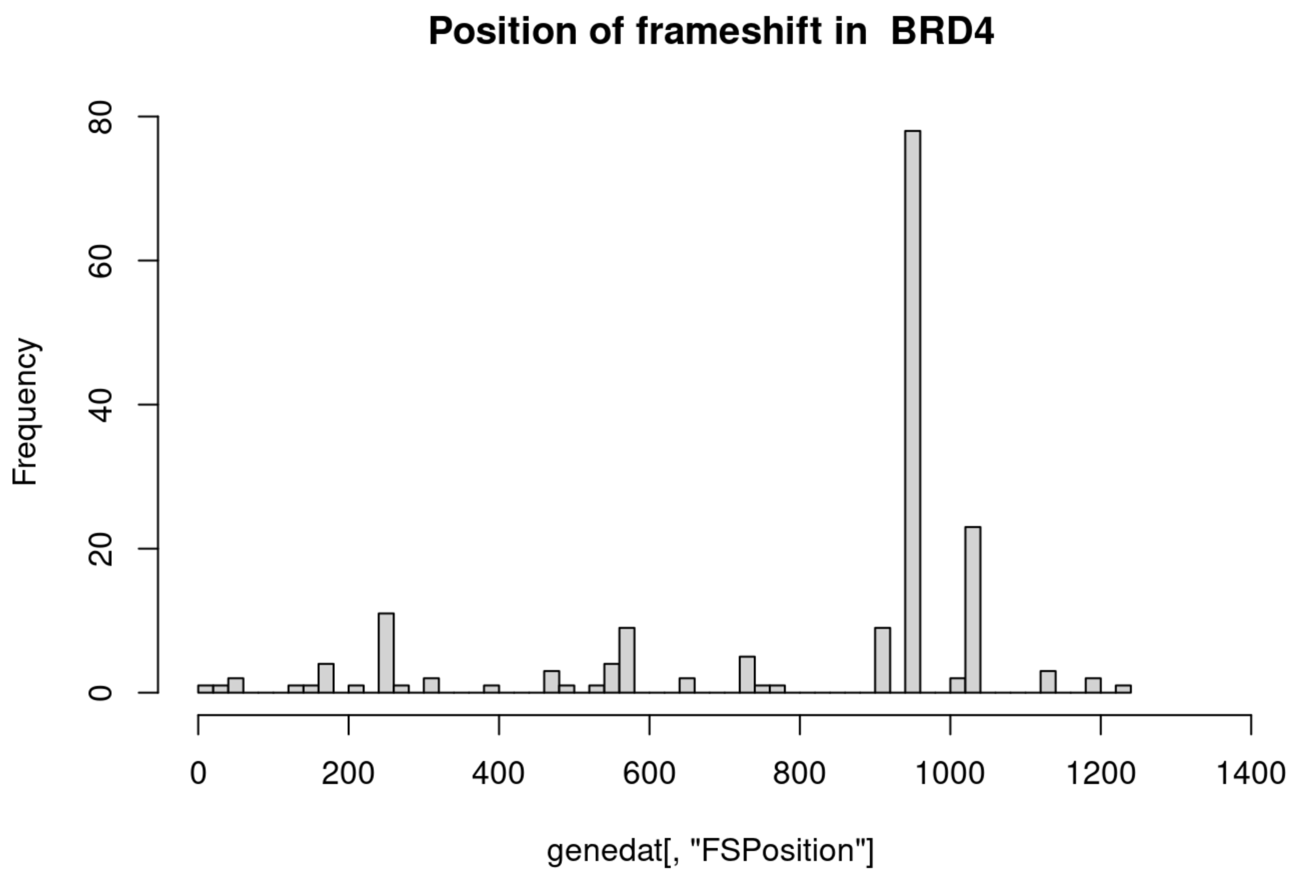
Position of frameshift in BRD4
Frequency
0
20
40
60
80
0
200
400
600
800
1000
1200
1400
genedat[, "FSPosition"]

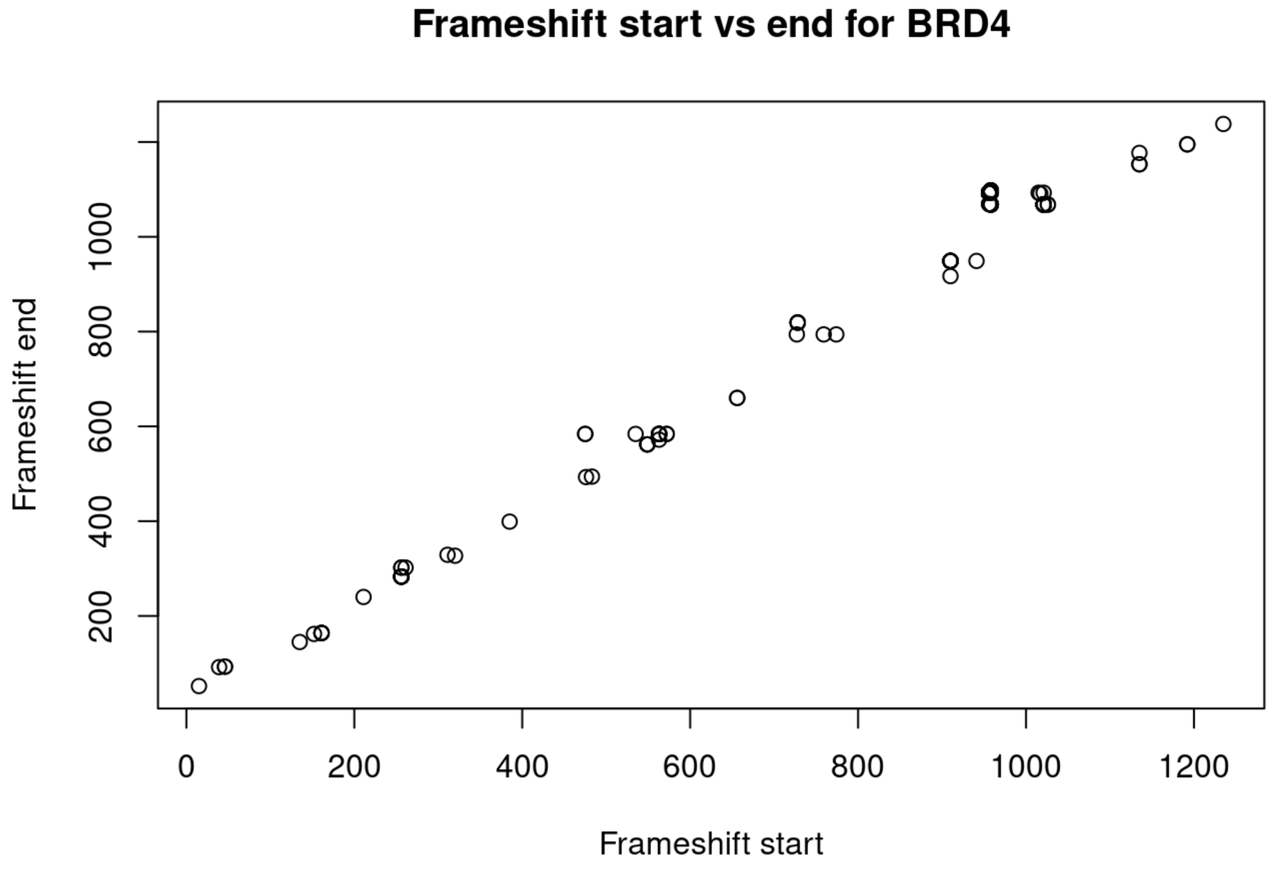
Frameshift start vs end for BRD4
Frameshift end
200
400
600
800
1000
0
200
400
600
800
1000
1200
Frameshift start

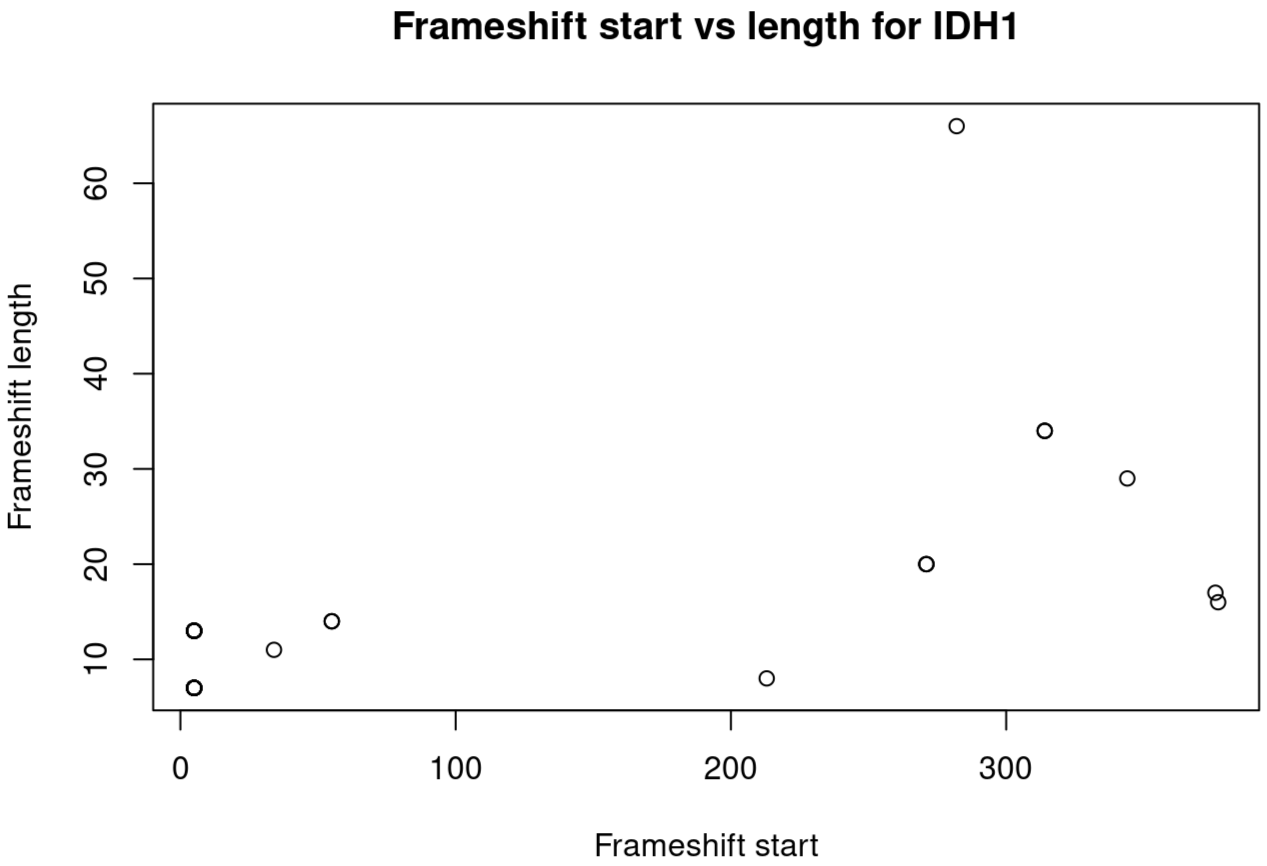
Frameshift start vs length for IDH1
Frameshift length
Frameshift start
10
20
30
40
50
60
0
100
200
300

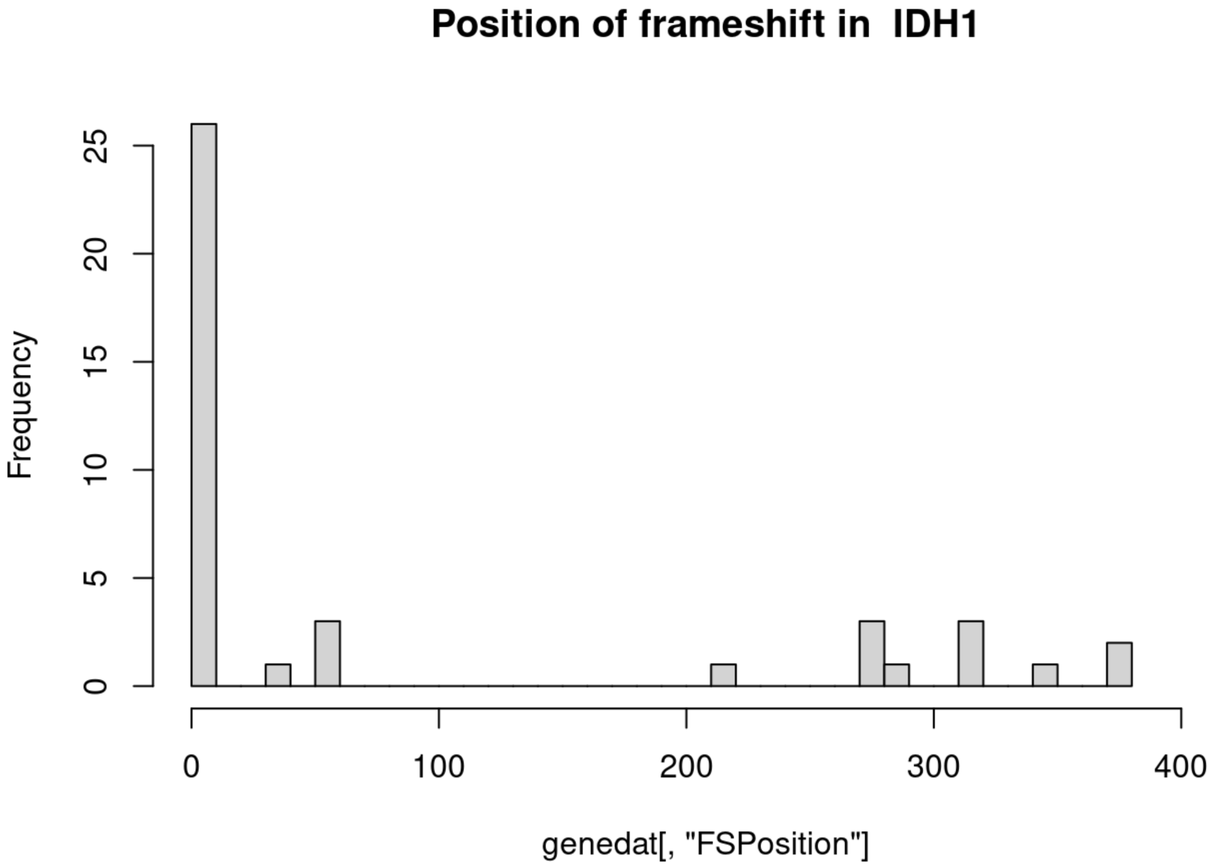
Position of frameshift in IDH1
Frequency
genedat[, "FSPosition"]
0
5
10
15
20
25
0
100
200
300
400

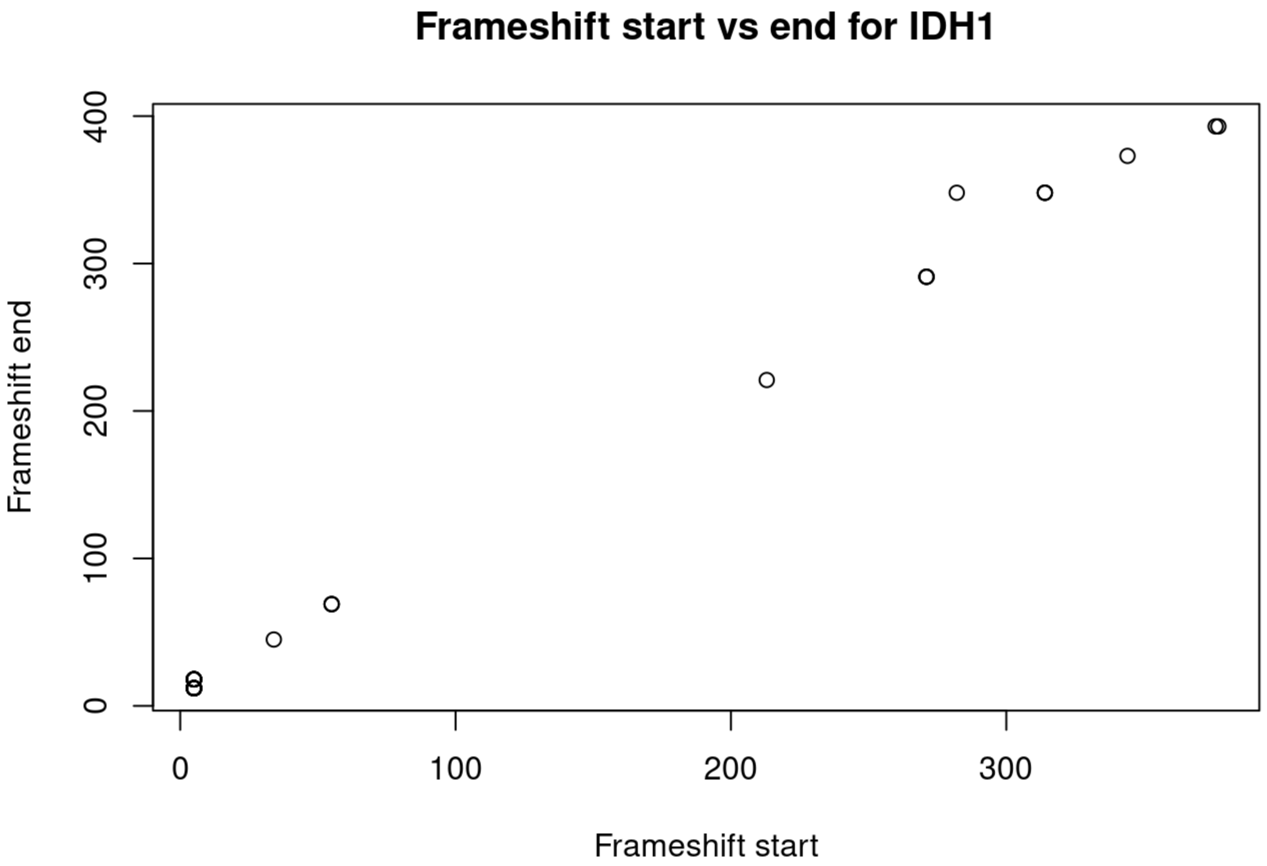
Frameshift start vs end for IDH1
Frameshift end
0
100
200
300
400
0
100
200
300
Frameshift start

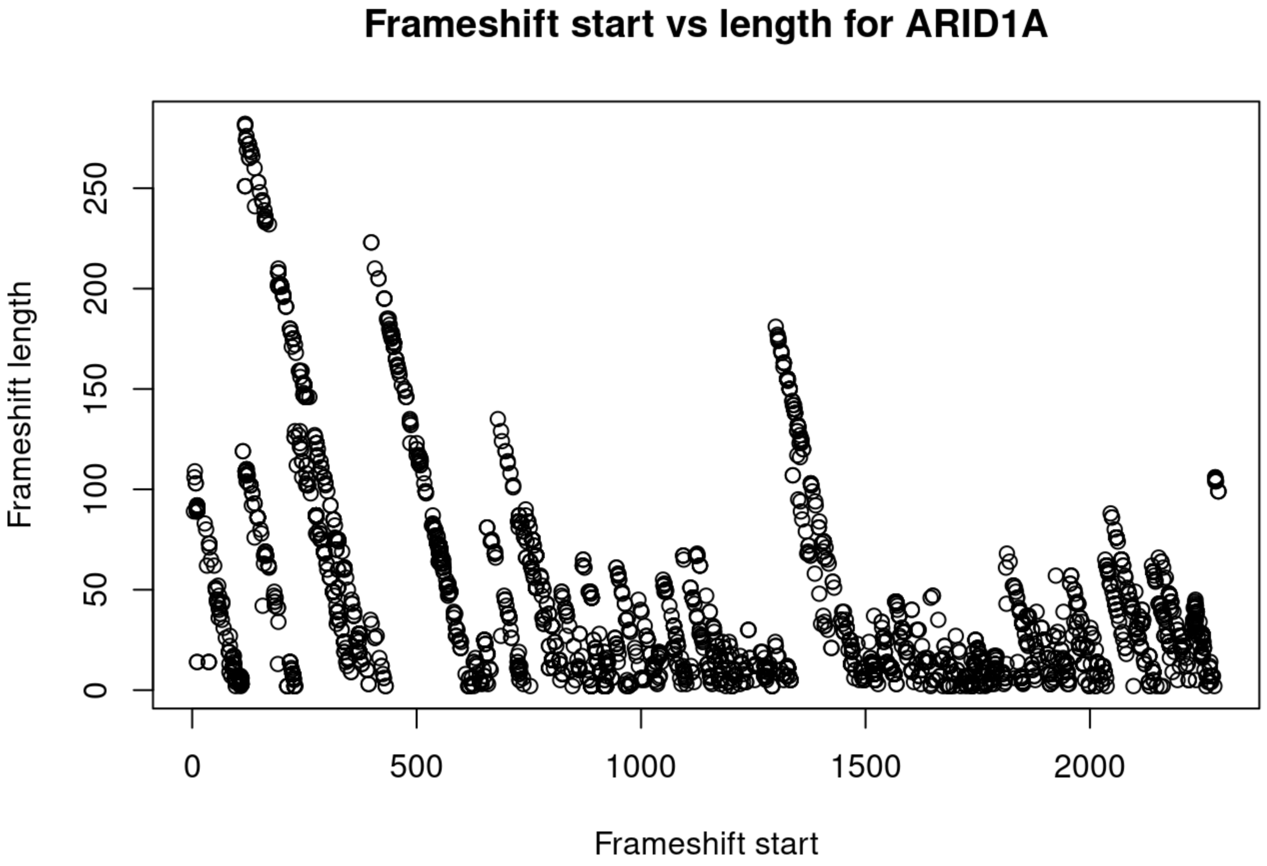
Frameshift start vs length for ARID1A
Frameshift length
0
50
100
150
200
250
0
500
1000
1500
2000
Frameshift start

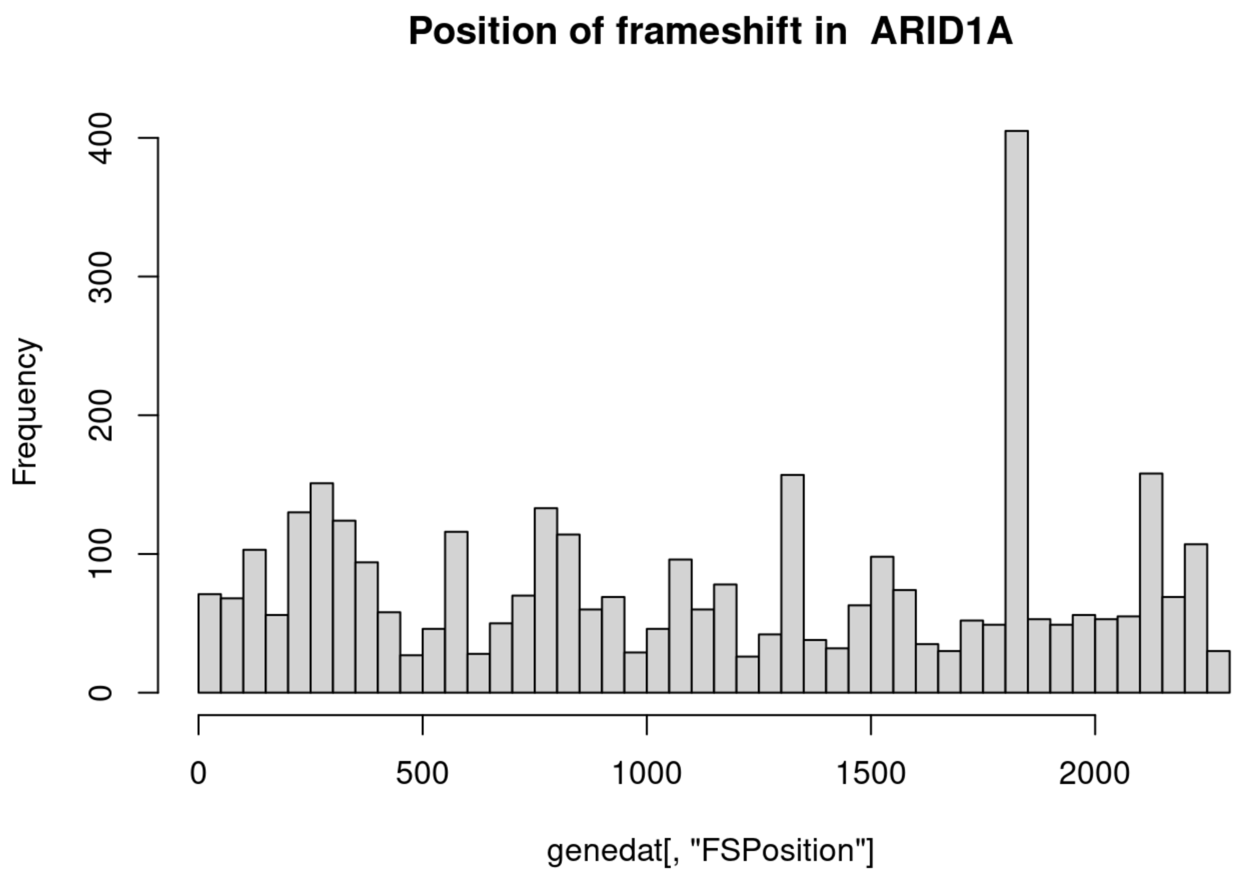
Position of frameshift in ARID1A
Frequency
0
100
200
300
400
0
500
1000
1500
2000
genedat[, "FSPosition"]

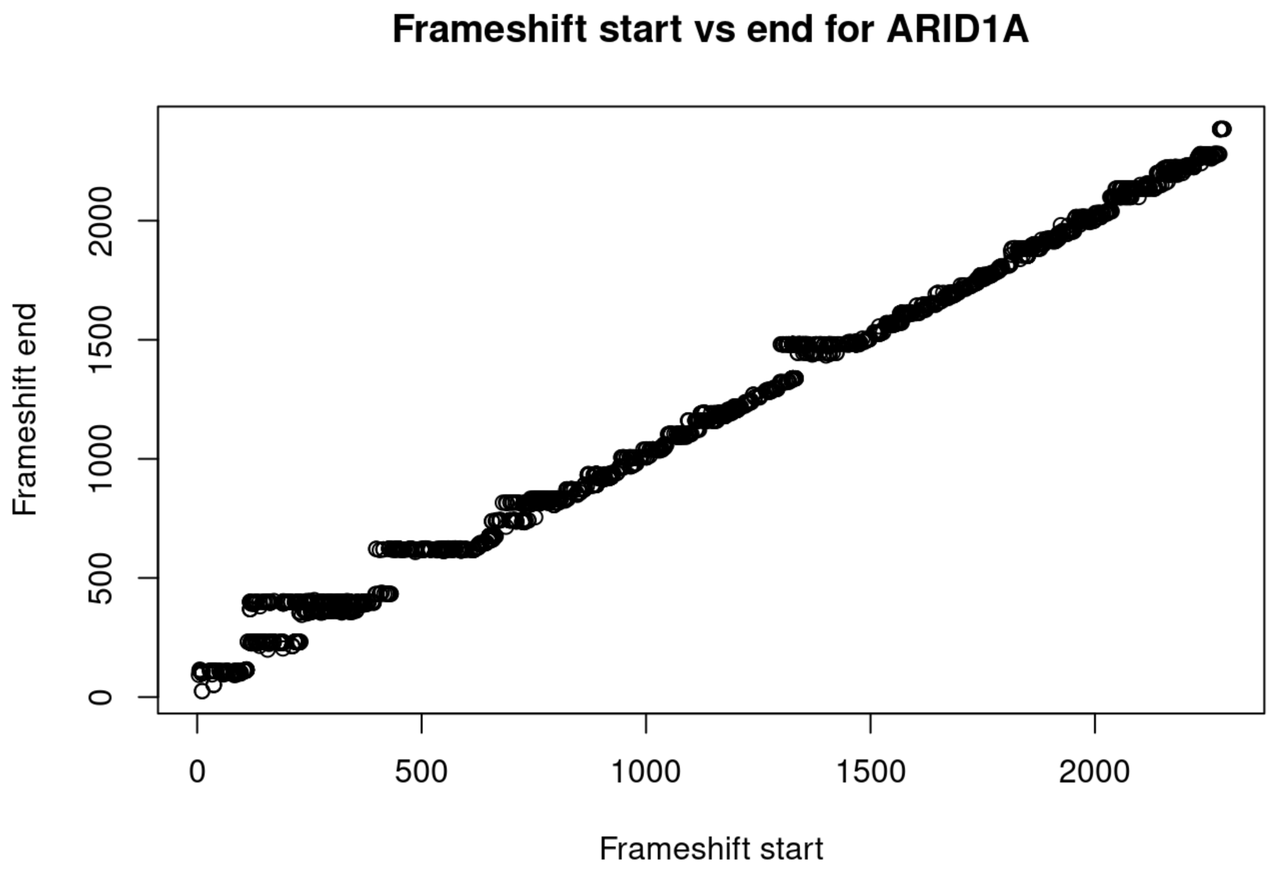
Frameshift start vs end for ARID1A
Frameshift end
0
500
1000
1500
2000
0
500
1000
1500
2000
Frameshift start

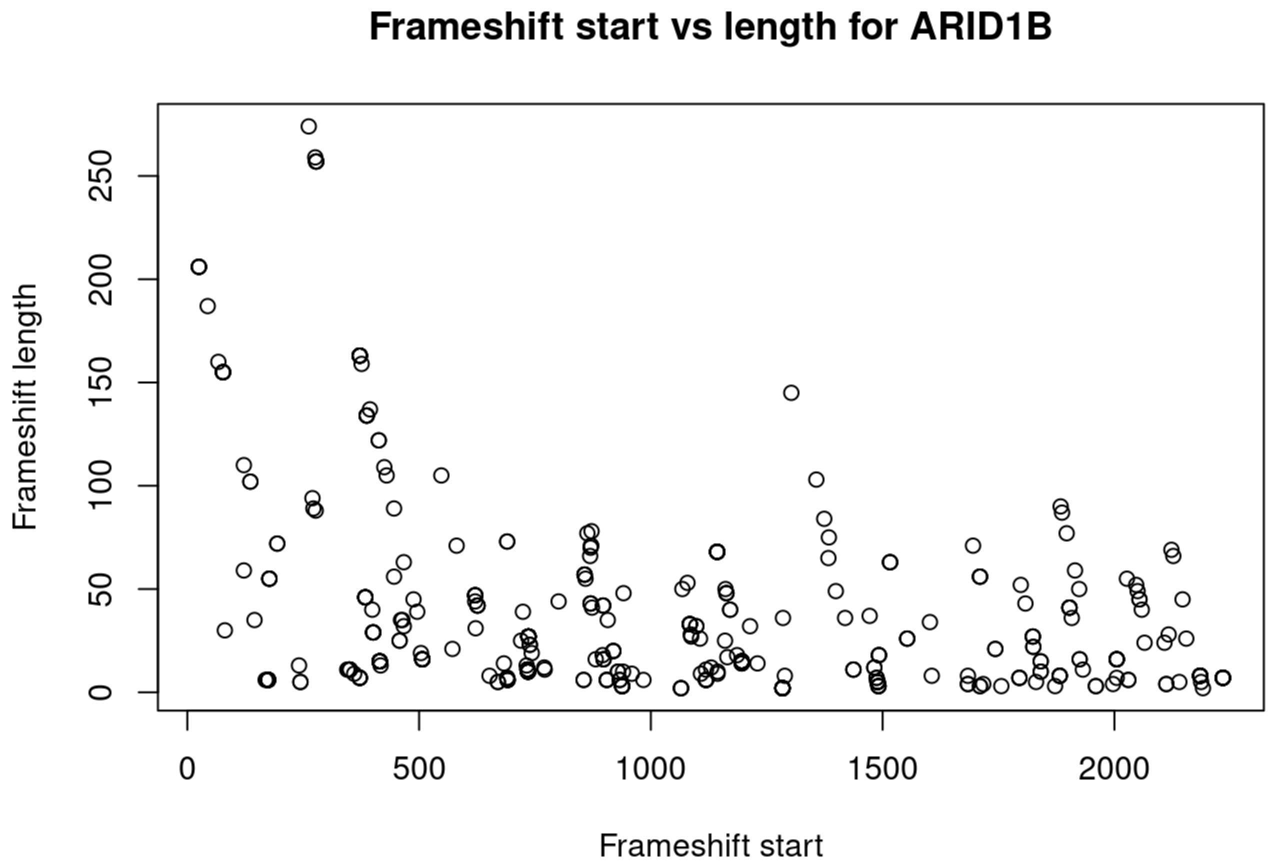
Frameshift start vs length for ARID1B
Frameshift length
Frameshift start
0
50
100
150
200
250
0
500
1000
1500
2000

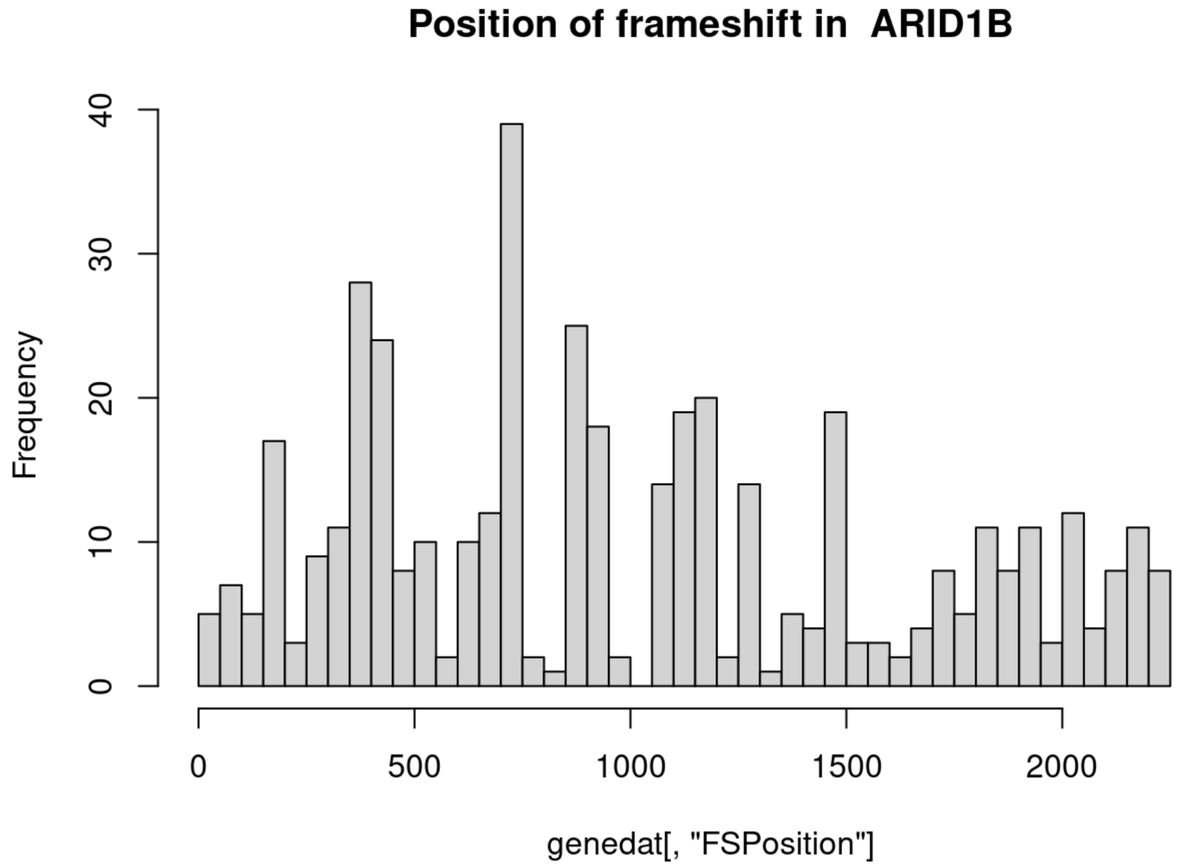
Position of frameshift in ARID1B
Frequency
genedat[, "FSPosition"]
0
10
20
30
40
0
500
1000
1500
2000

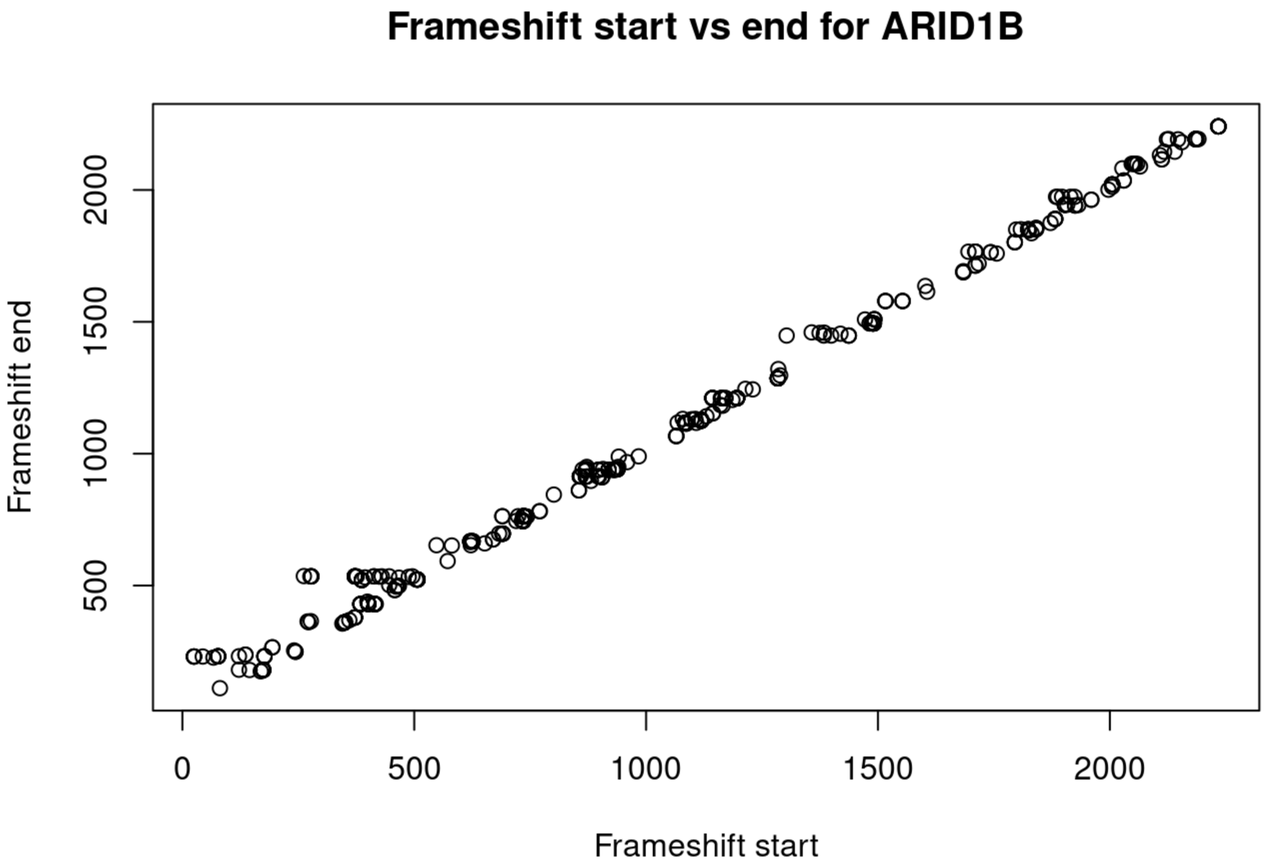
Frameshift start vs end for ARID1B
Frameshift end
Frameshift start
500
1000
1500
2000
0
500
1000
1500
2000

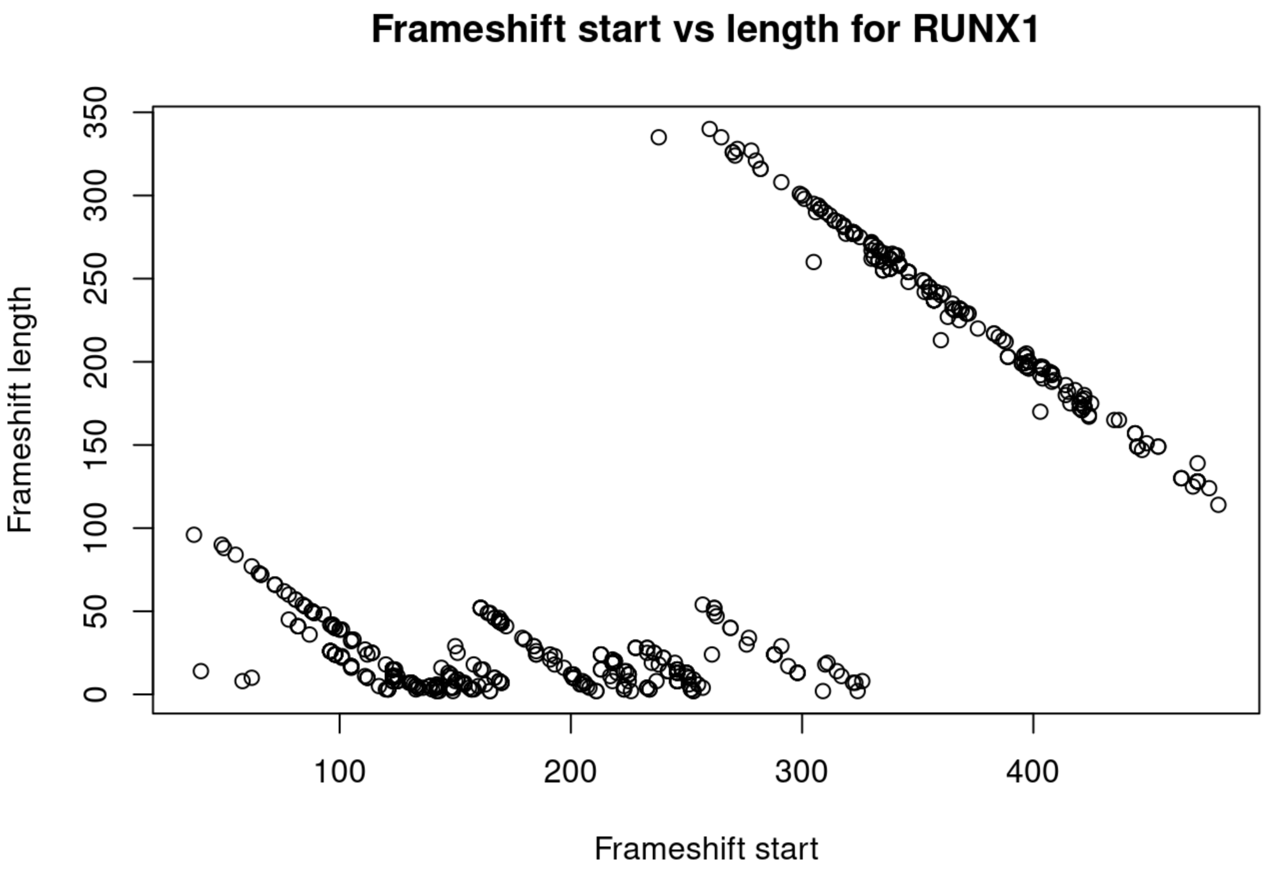
Frameshift start vs length for RUNX1
Frameshift length
Frameshift start
0
50
100
150
200
250
300
350
100
200
300
400

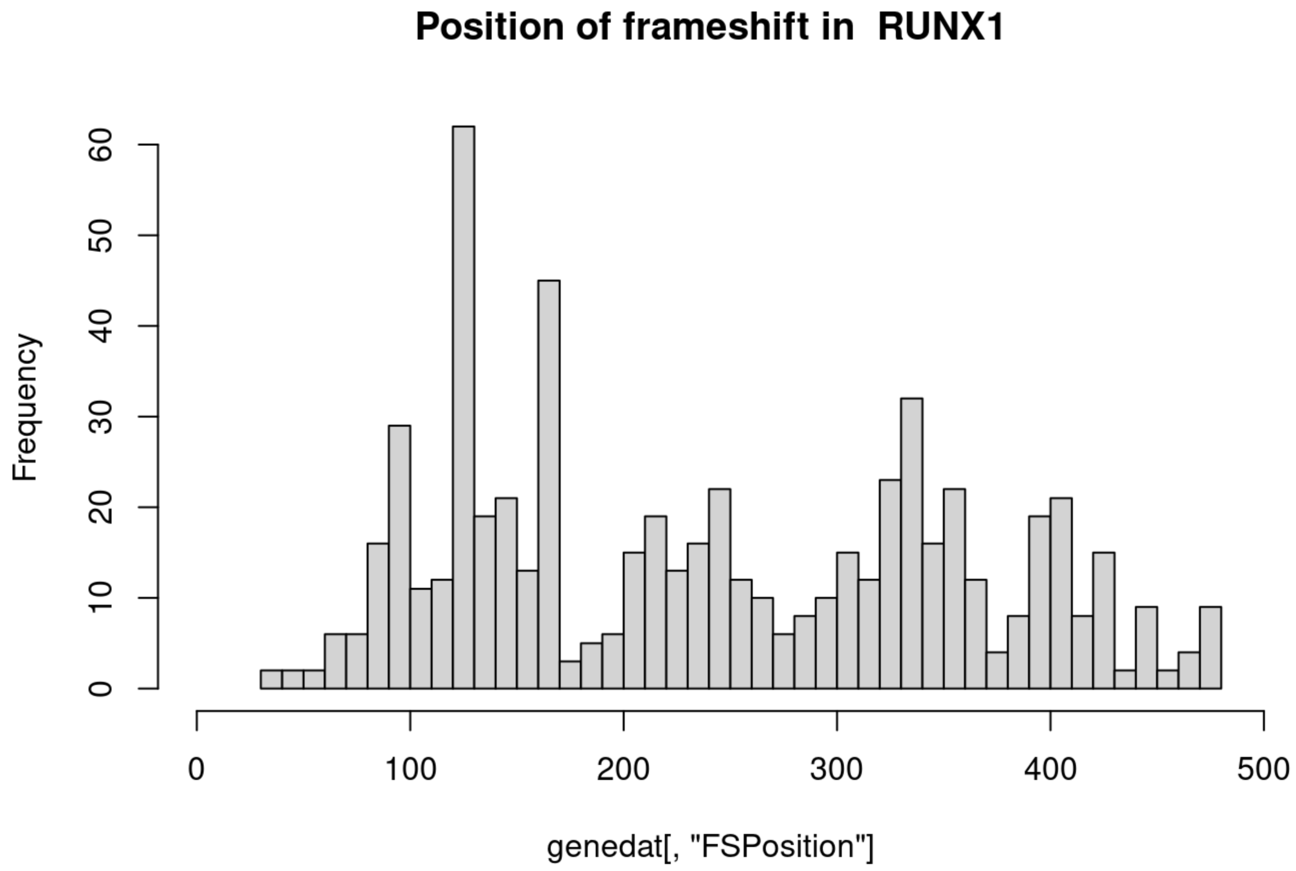
Position of frameshift in RUNX1
Frequency
0
10
20
30
40
50
60
0
100
200
300
400
500
genedat[, "FSPosition"]

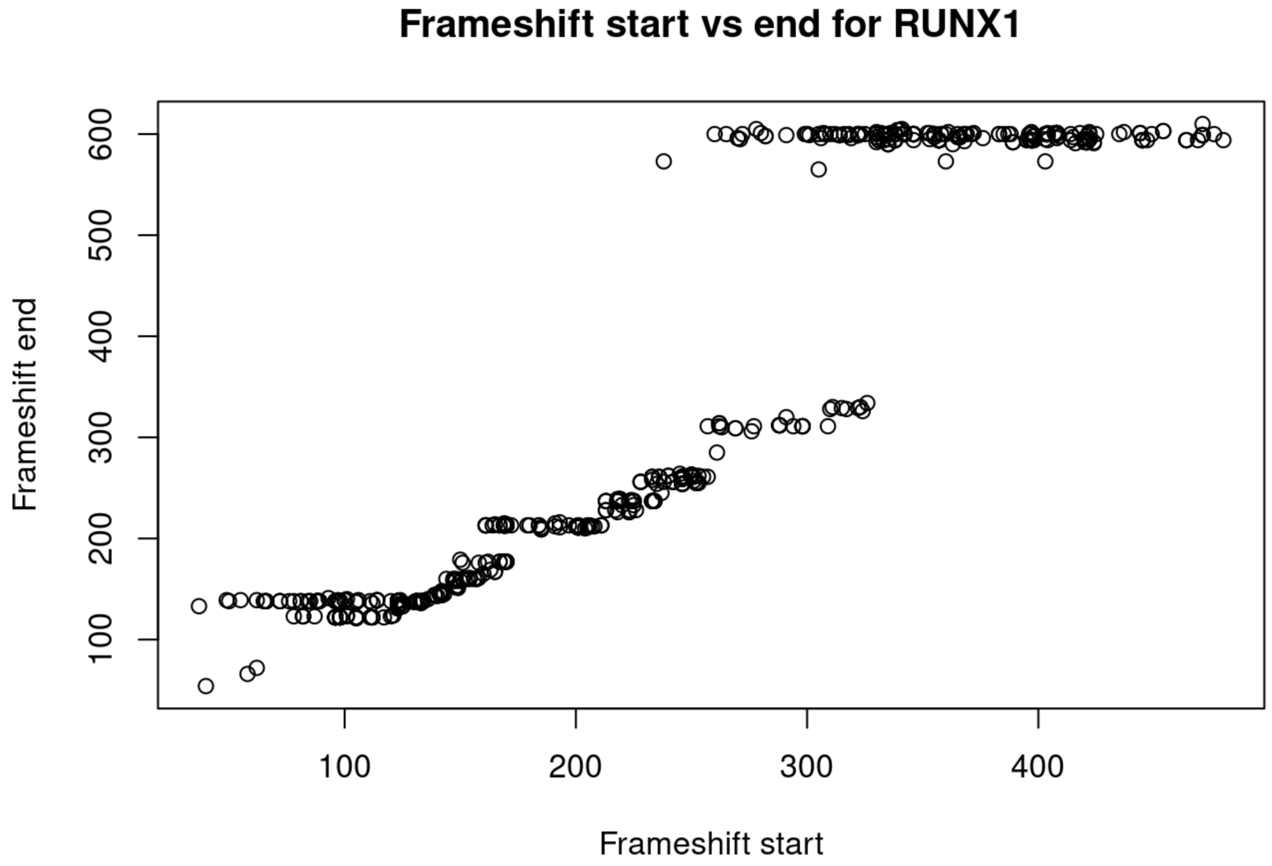
Frameshift start vs end for RUNX1
Frameshift end
100
200
300
400
500
600
100
200
300
400
Frameshift start

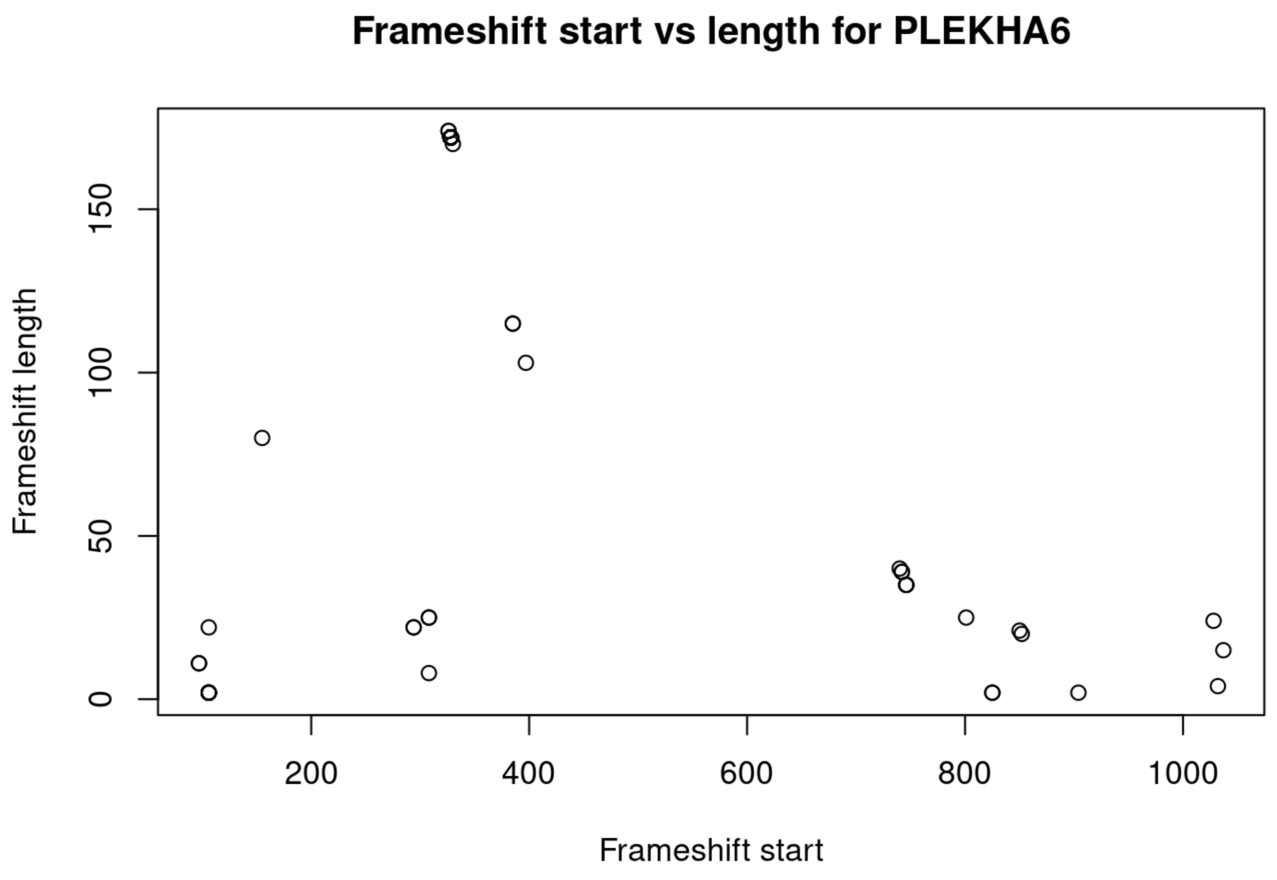
Frameshift start vs length for PLEKHA6
Frameshift length
Frameshift start
0
50
100
150
200
400
600
800
1000

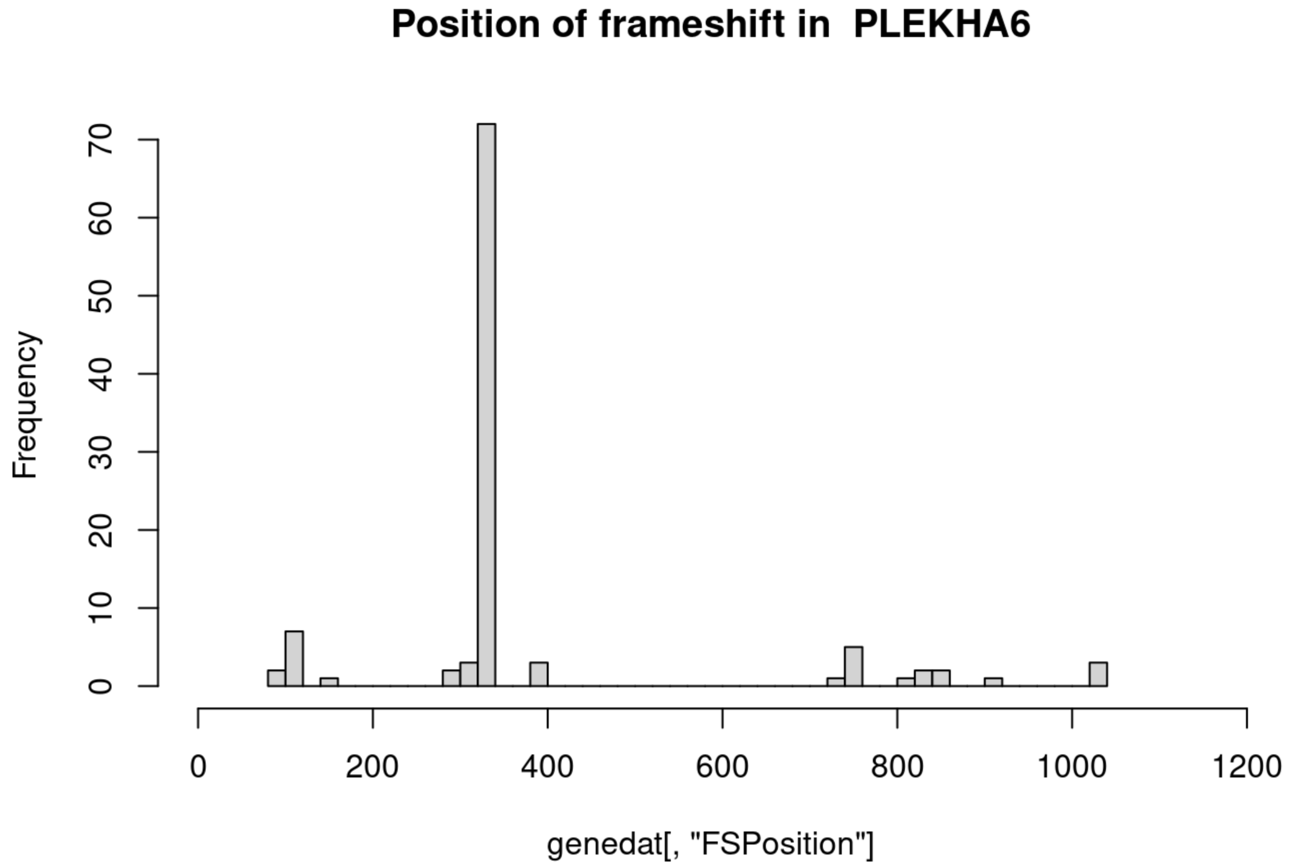
Position of frameshift in PLEKHA6
Frequency
genedat[, "FSPosition"]
0
10
20
30
40
50
60
70
0
200
400
600
800
1000
1200

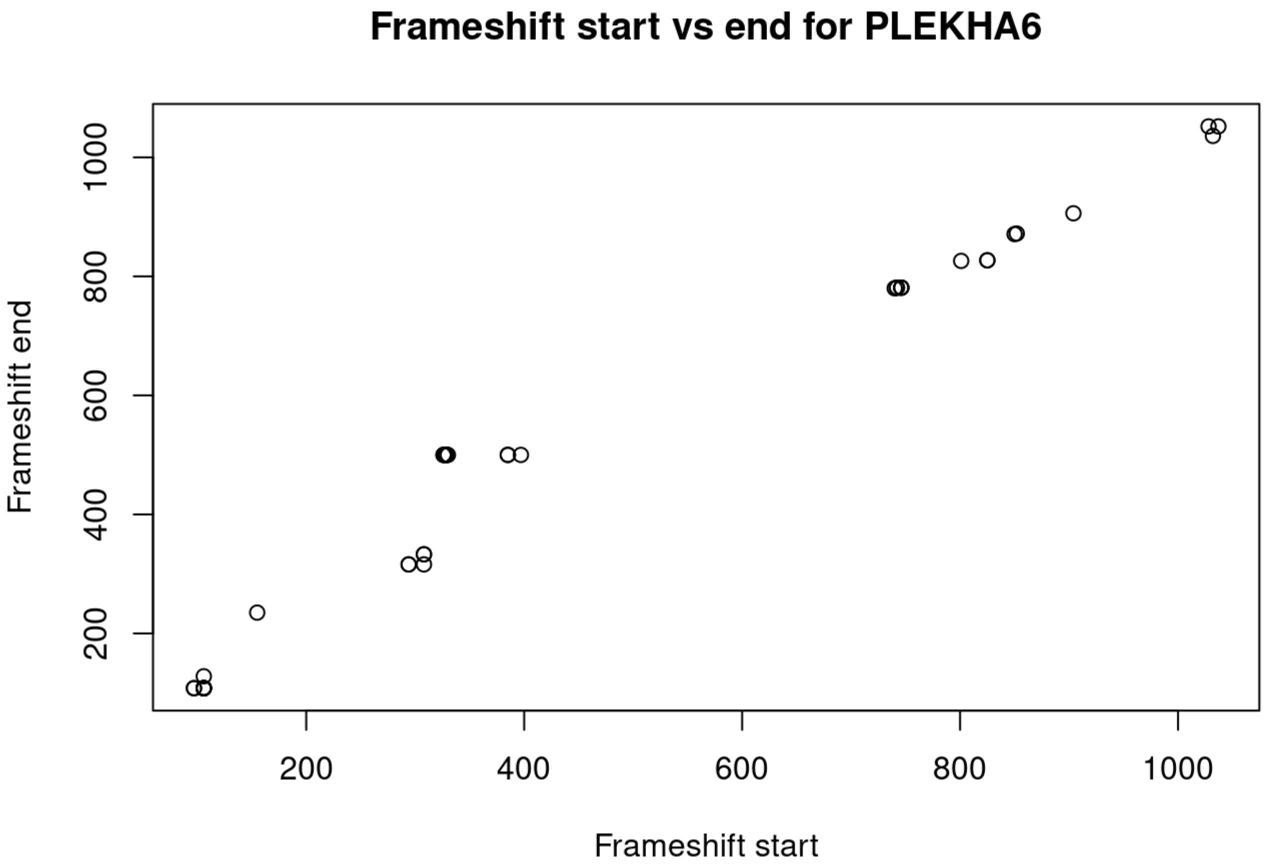
Frameshift start vs end for PLEKHA6
Frameshift end
Frameshift start
200
400
600
800
1000

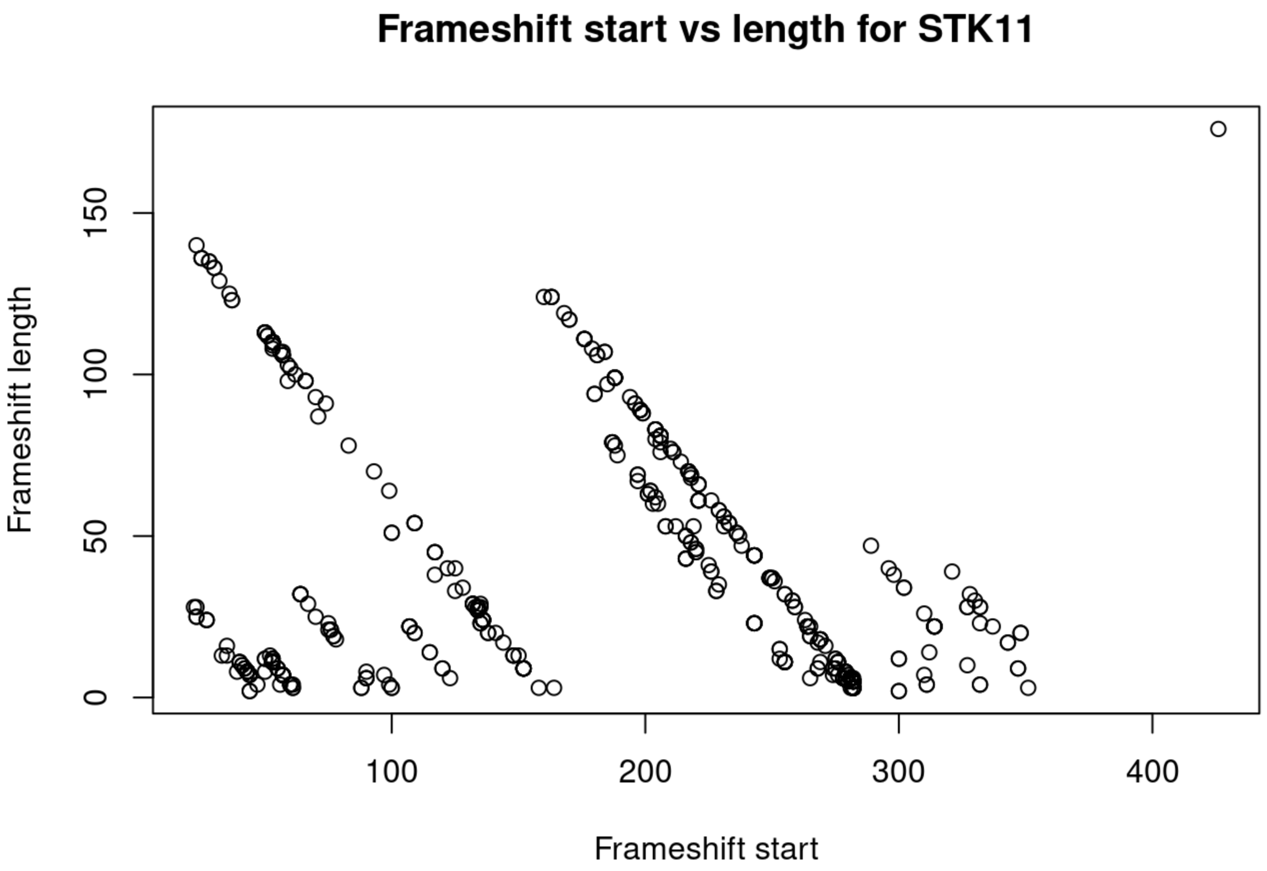
Frameshift start vs length for STK11
Frameshift length
Frameshift start
0
50
100
150
200
300
400

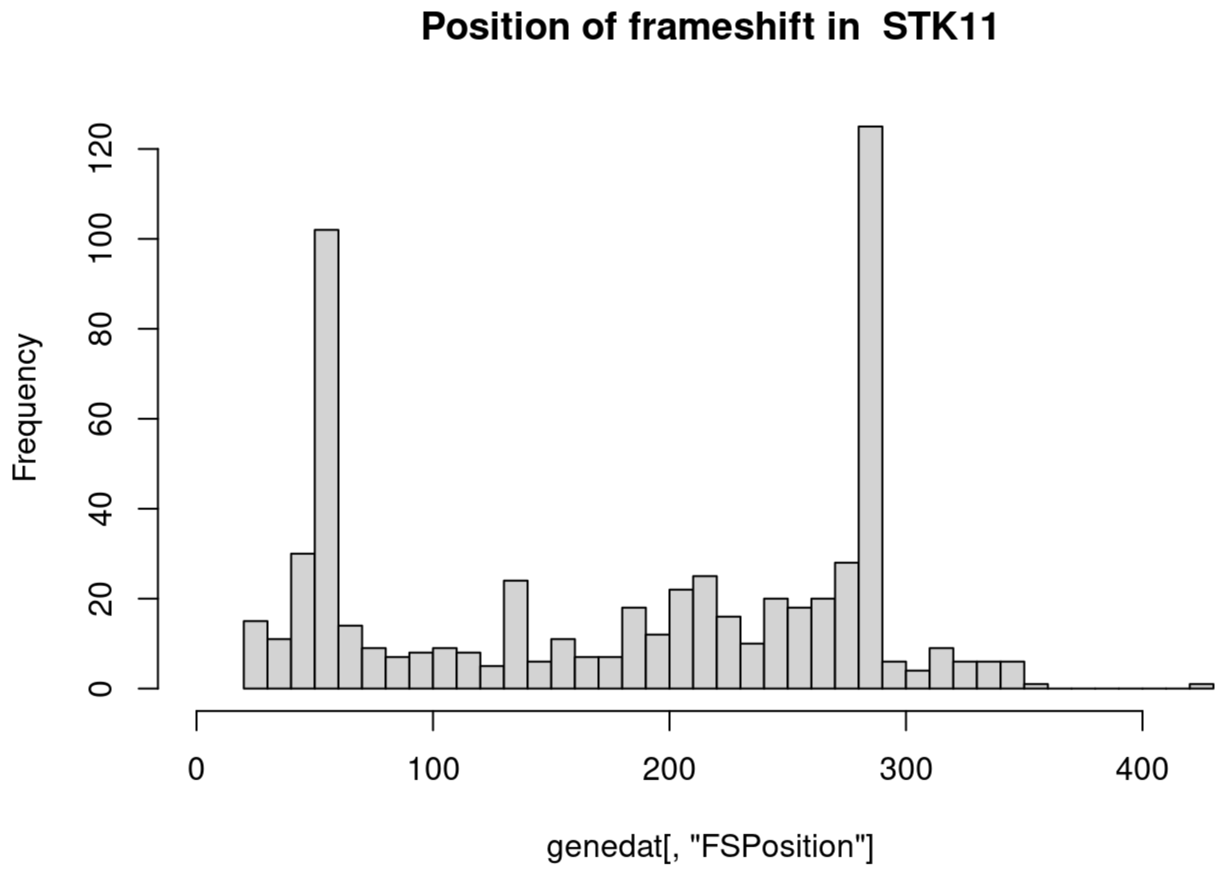
Position of frameshift in STK11
Frequency
0
20
40
60
80
100
120
0
100
200
300
400
genedat[, "FSPosition"]

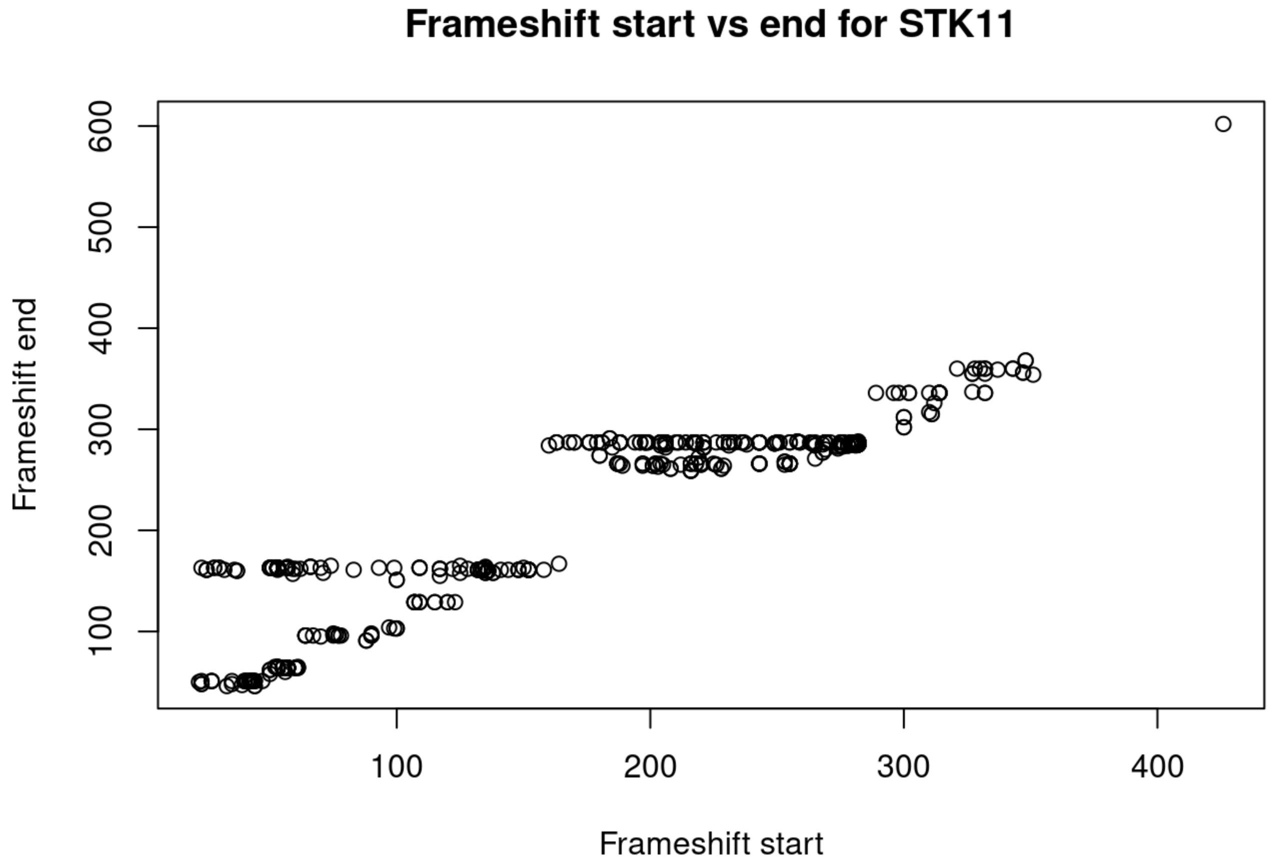
Frameshift start vs end for STK11
Frameshift end
100
200
300
400
500
600
100
200
300
400
Frameshift start

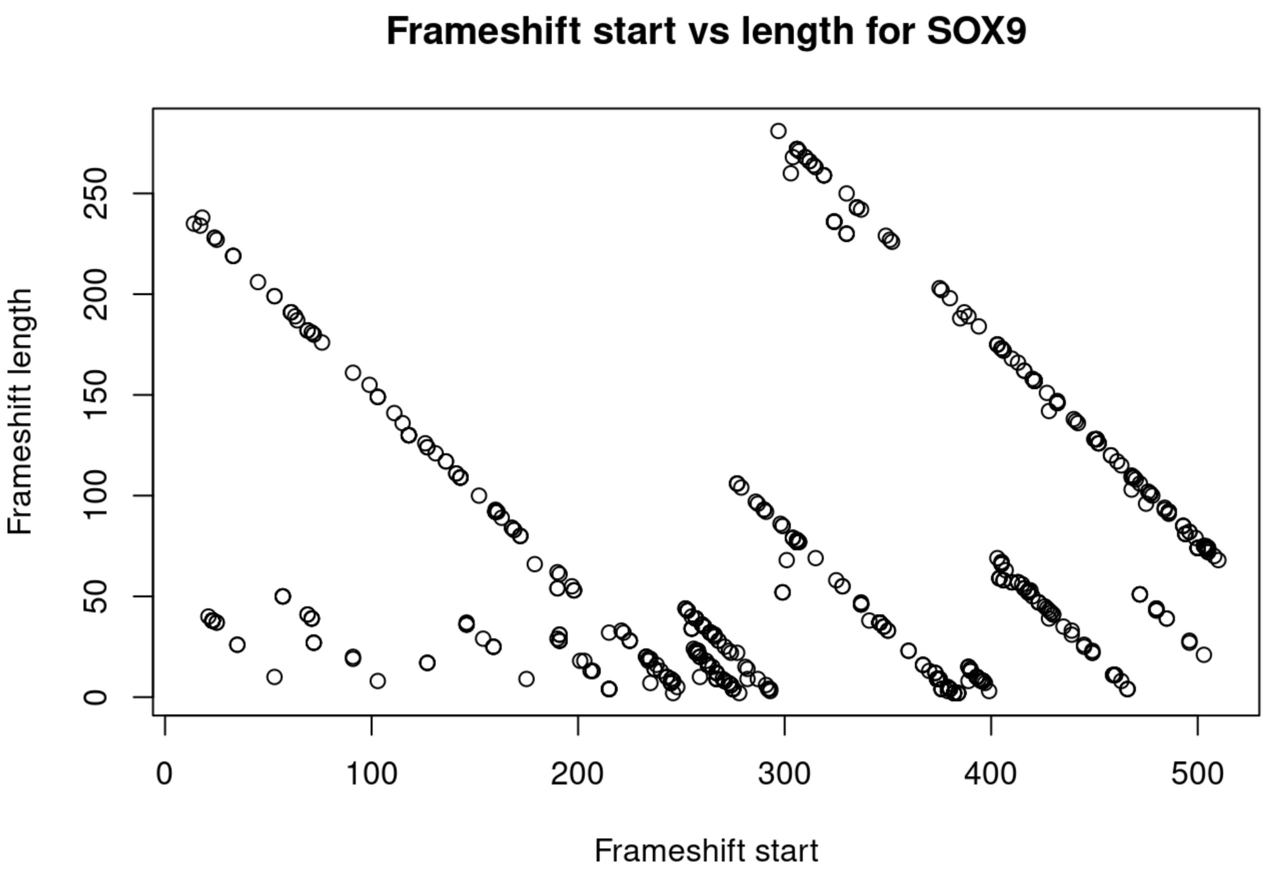
Frameshift start vs length for SOX9
Frameshift length
Frameshift start
0
50
100
150
200
250
0
100
200
300
400
500

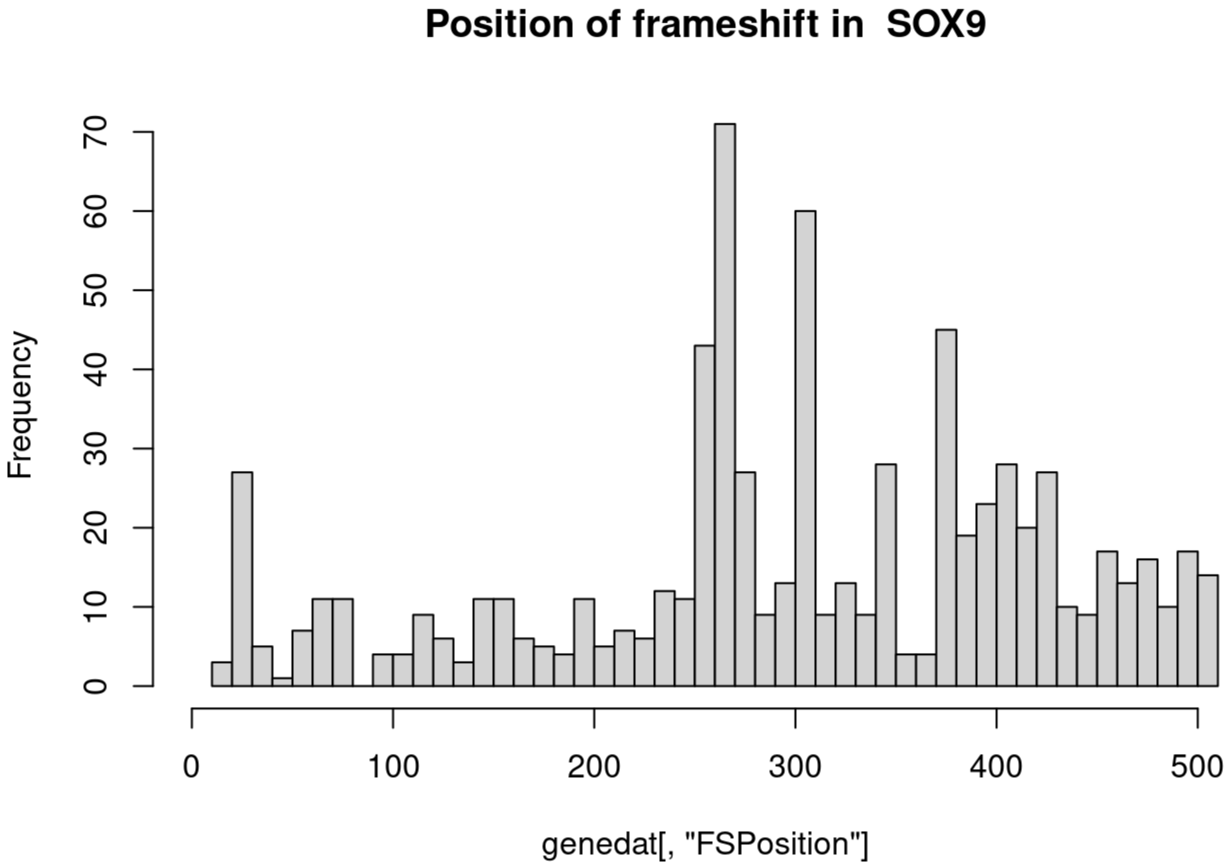
Position of frameshift in SOX9
Frequency
genedat[, "FSPosition"]
0
10
20
30
40
50
60
70
0
100
200
300
400
500

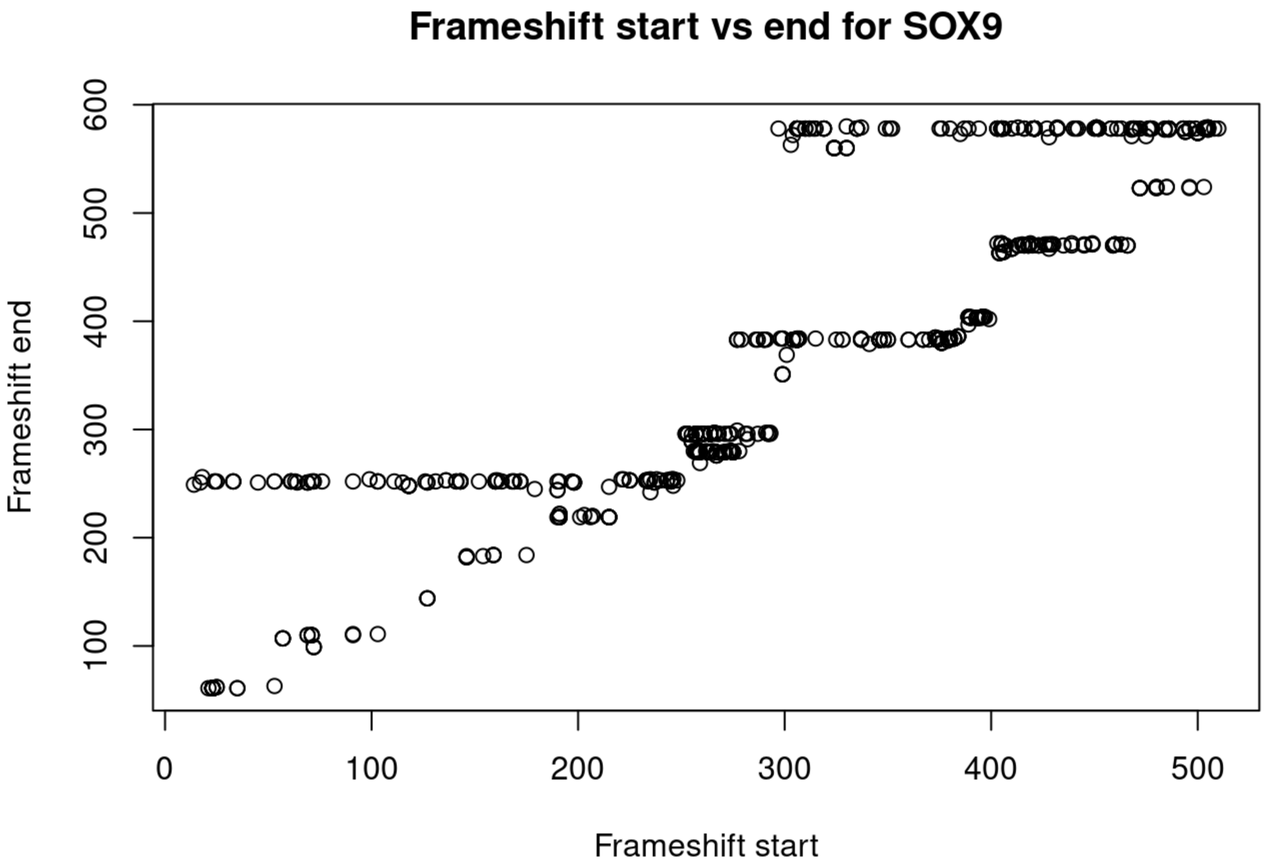
Frameshift start vs end for SOX9
Frameshift end
Frameshift start
600
500
400
300
200
100
0
100
200
300
400
500

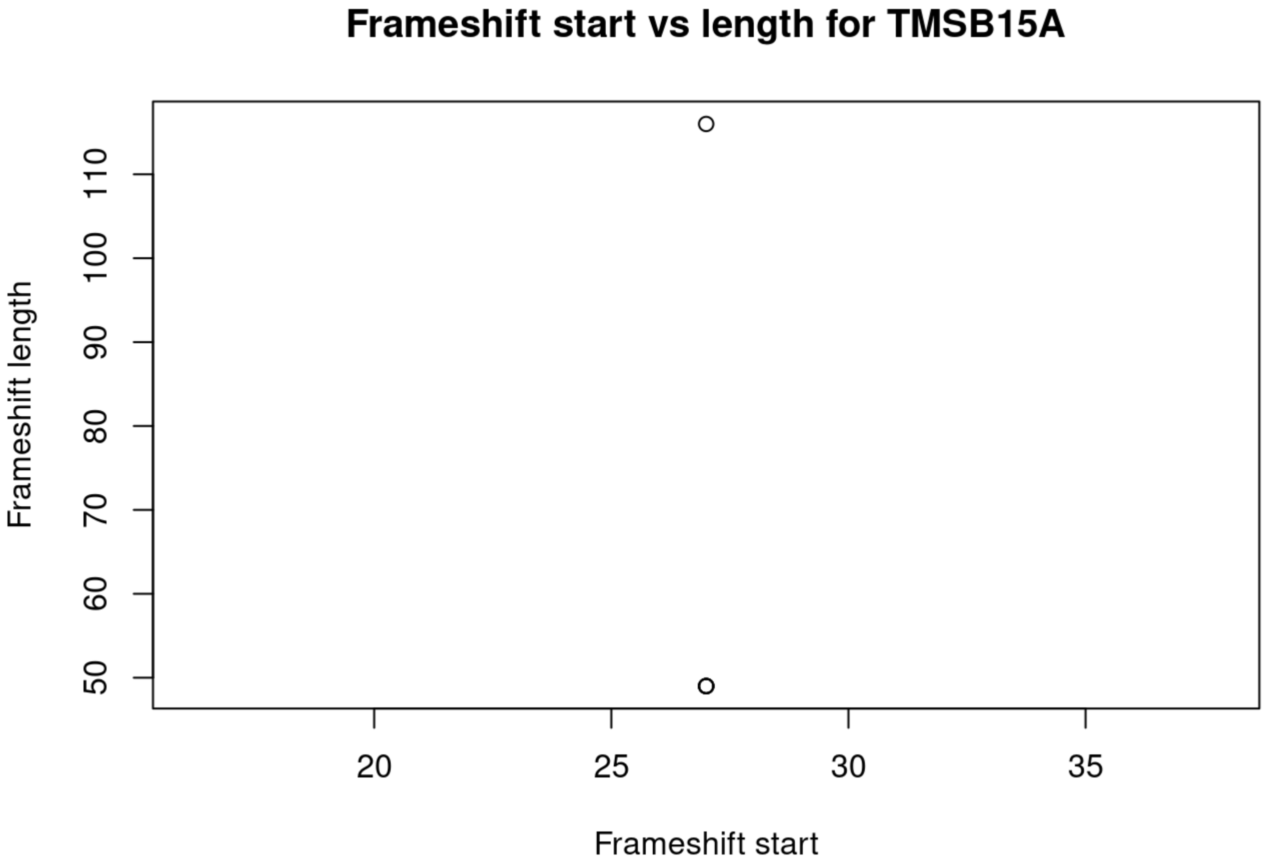
Frameshift start vs length for TMSB15A
Frameshift length
Frameshift start
110
100
90
80
70
60
50
20
25
30
35

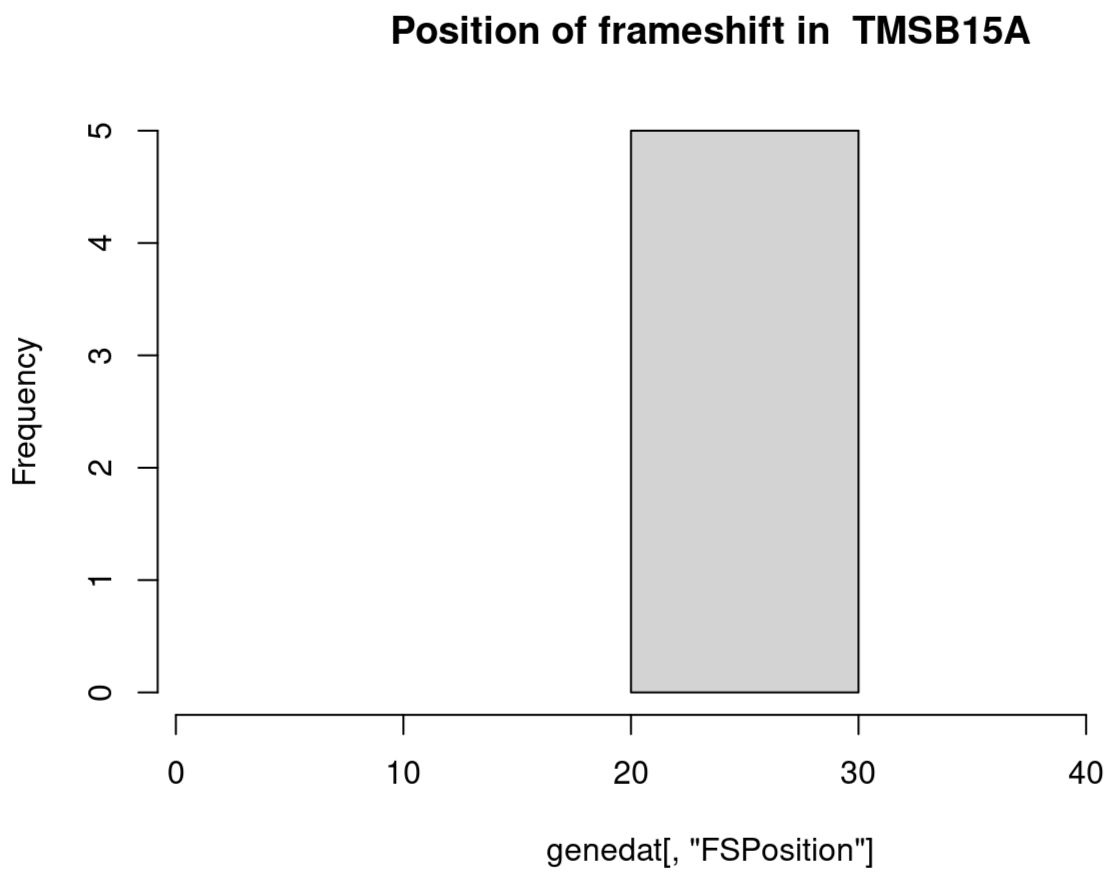
Position of frameshift in TMSB15A
Frequency
0
1
2
3
4
5
0
10
20
30
40
genedat[, "FSPosition"]

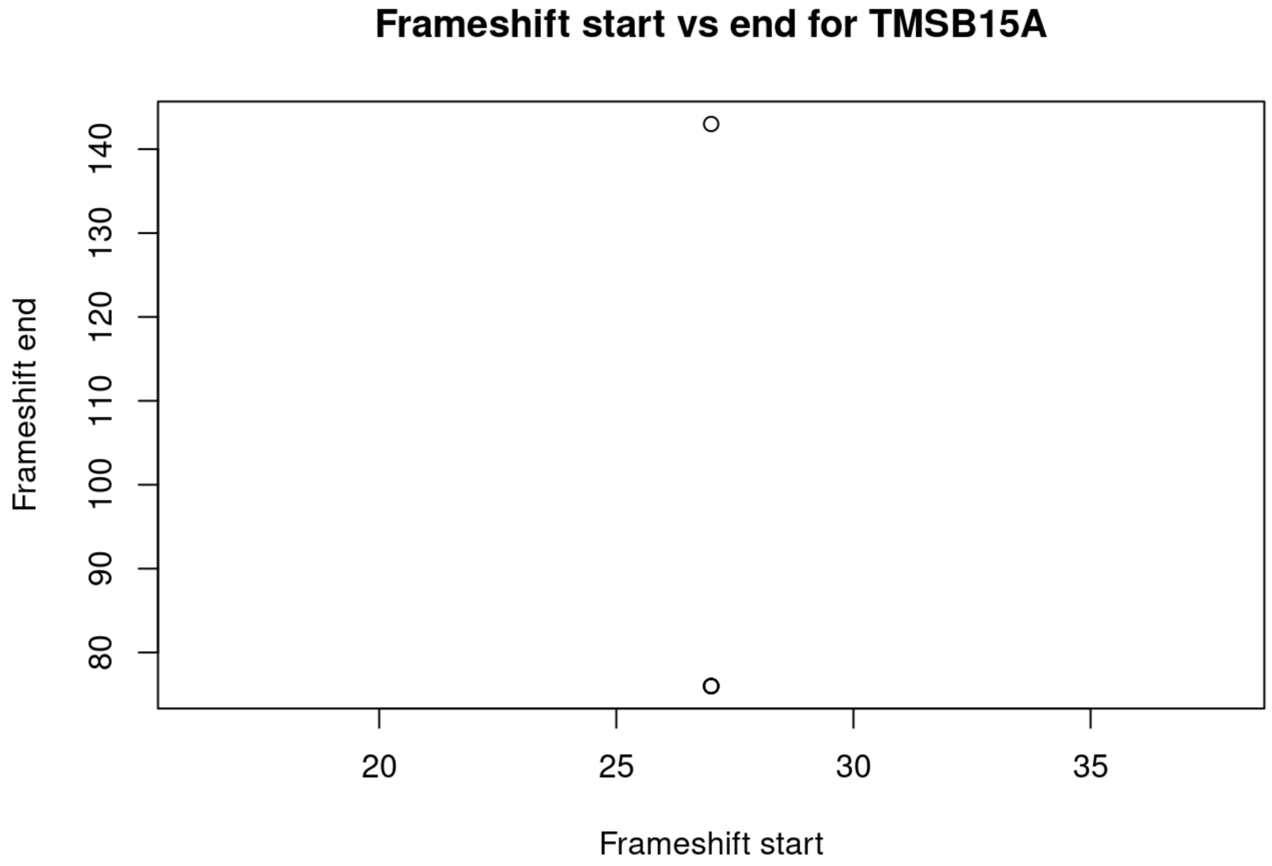
Frameshift start vs end for TMSB15A
Frameshift end
80
90
100
110
120
130
140
20
25
30
35
Frameshift start

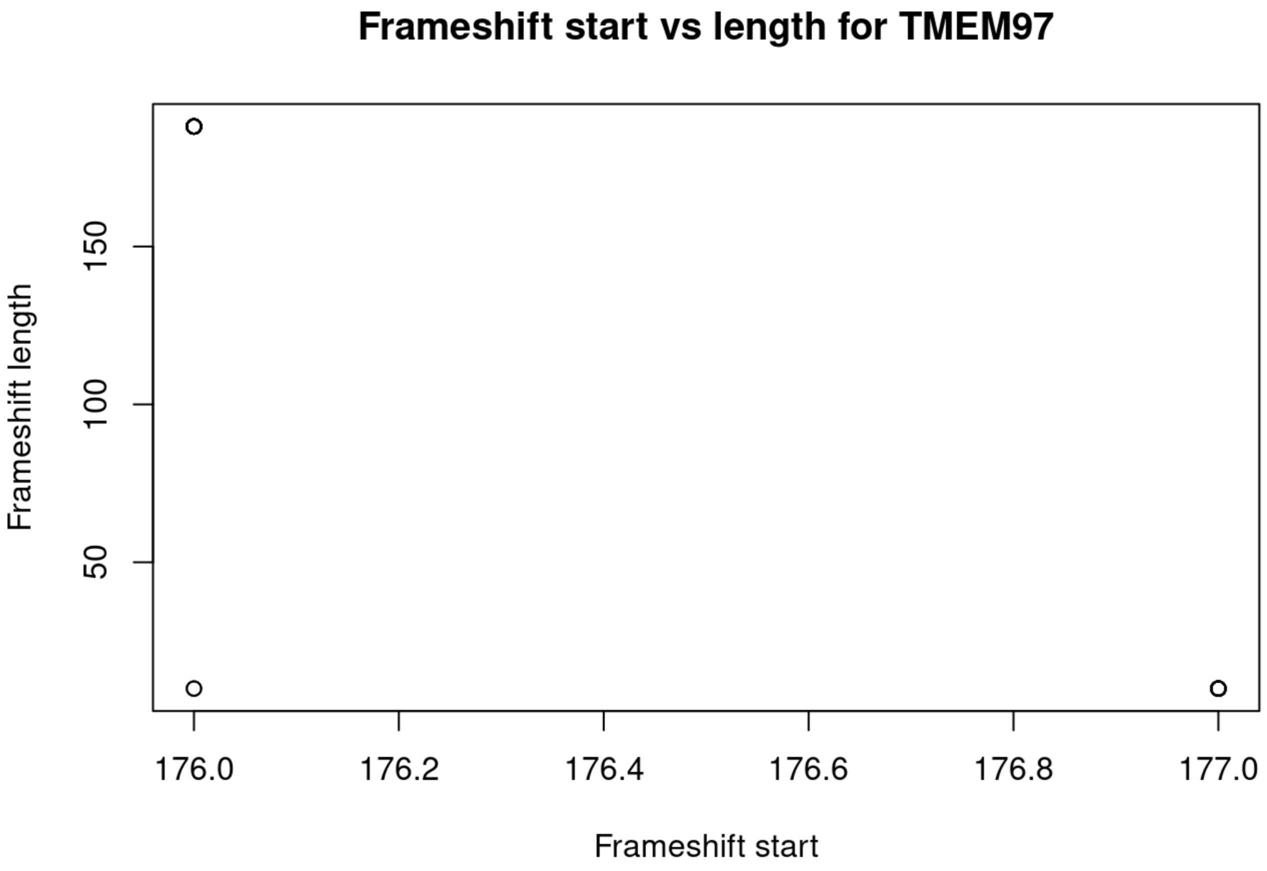
Frameshift start vs length for TMEM97
Frameshift length
50
100
150
176.0
176.2
176.4
176.6
176.8
177.0
Frameshift start

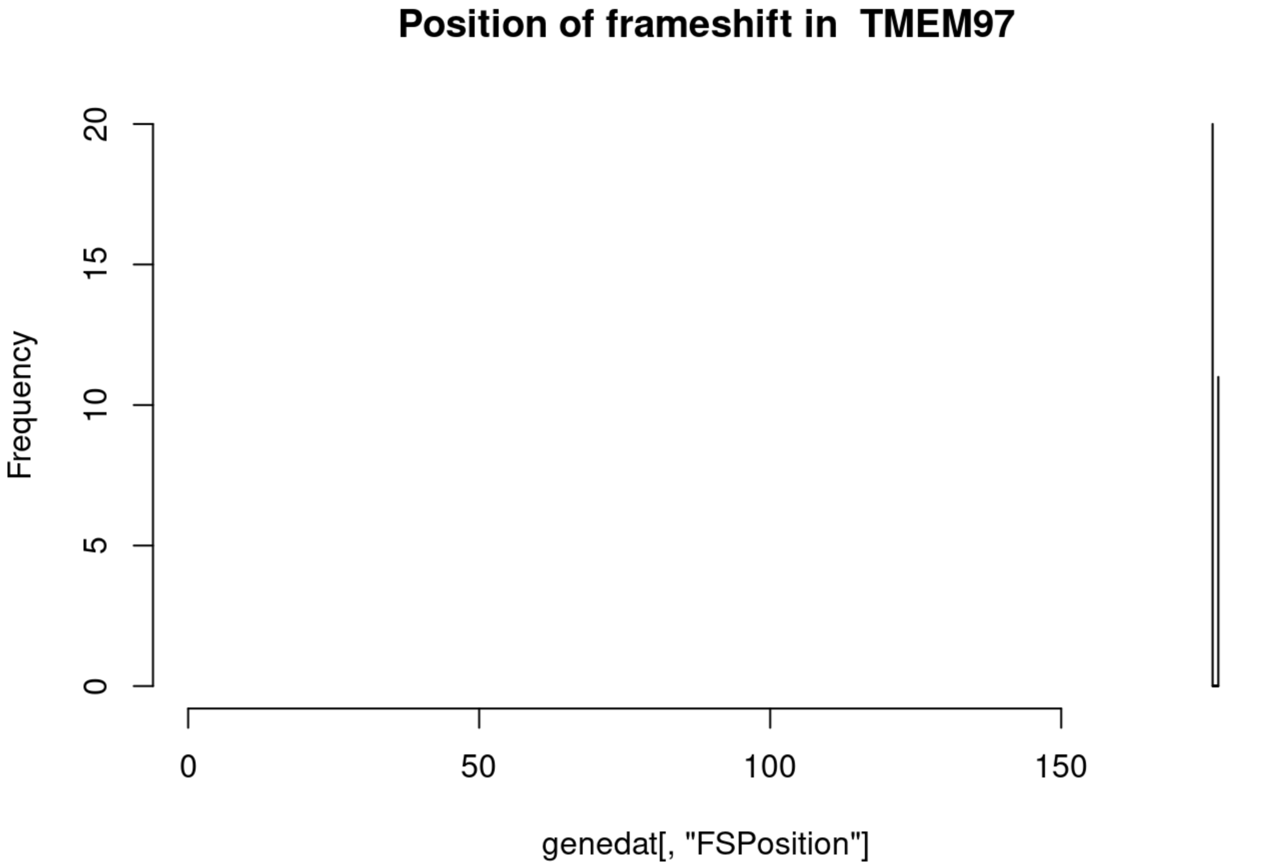
Position of frameshift in TMEM97
Frequency
0
5
10
15
20
0
50
100
150
genedat[, "FSPosition"]

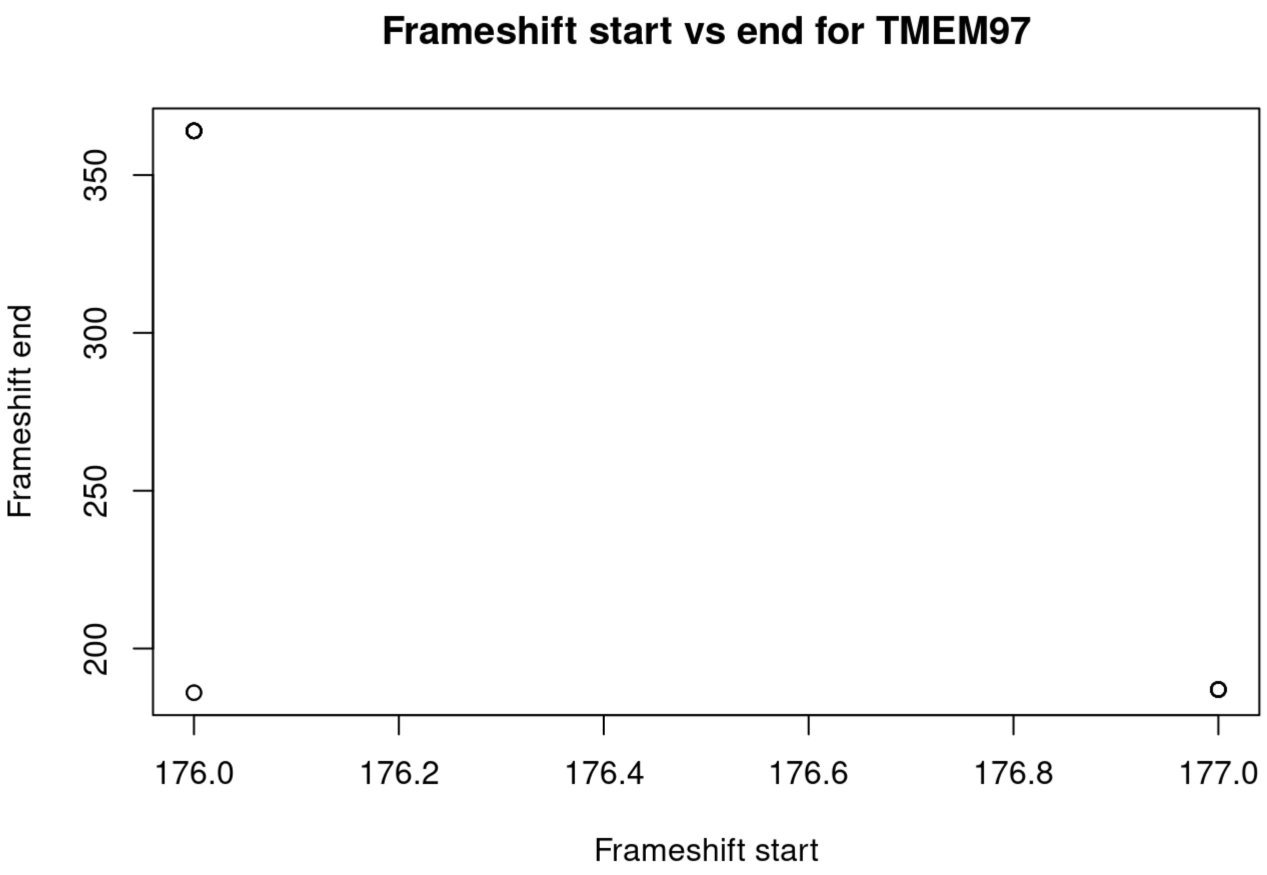
Frameshift start vs end for TMEM97
Frameshift end
350
300
250
200
176.0
176.2
176.4
176.6
176.8
177.0
Frameshift start

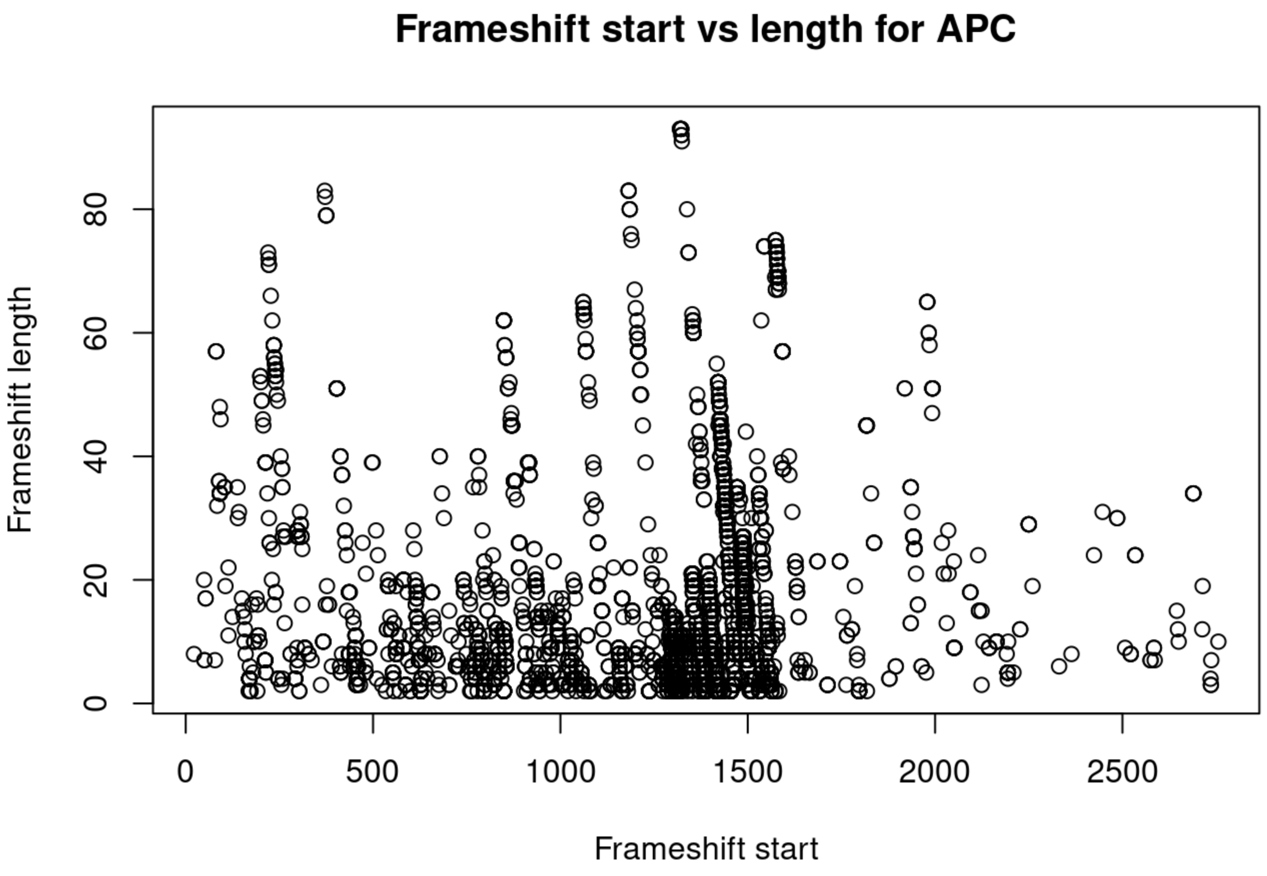
Frameshift start vs length for APC
Frameshift length
80
60
40
20
0
0
500
1000
1500
2000
2500
Frameshift start

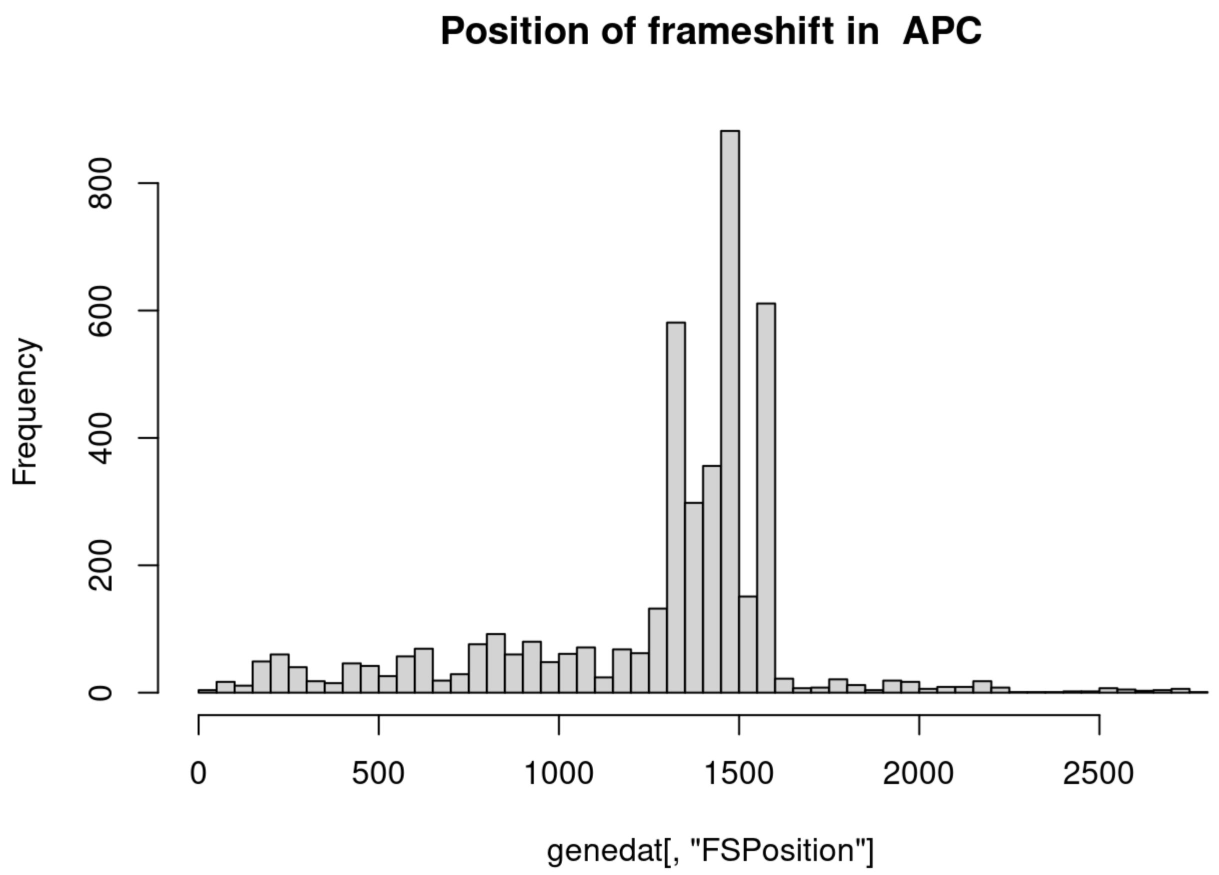

Position of frameshift in APC
Frequency
0
200
400
600
800
0
500
1000
1500
2000
2500
genedat[, "FSPosition"]


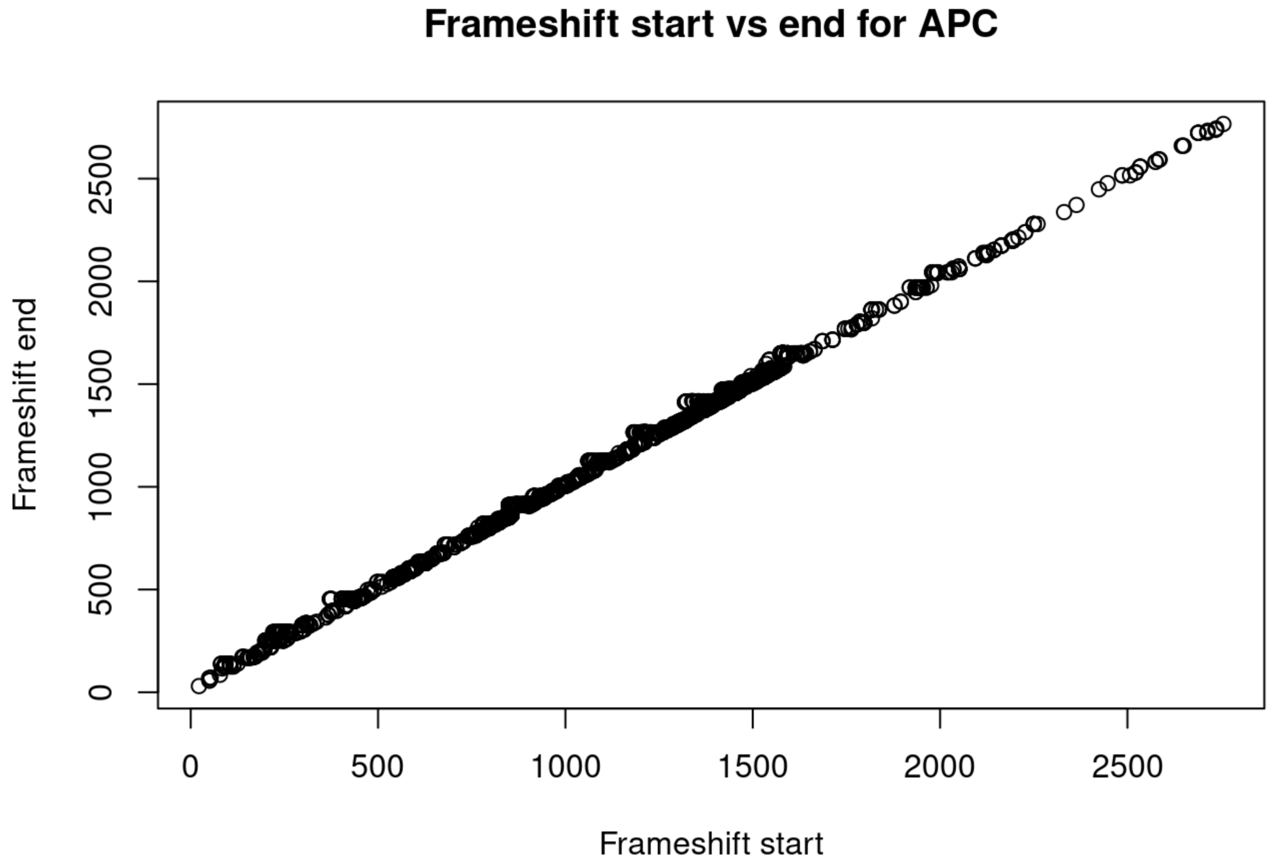

Frameshift start vs end for APC
Frameshift end
0
500
1000
1500
2000
2500
Frameshift start

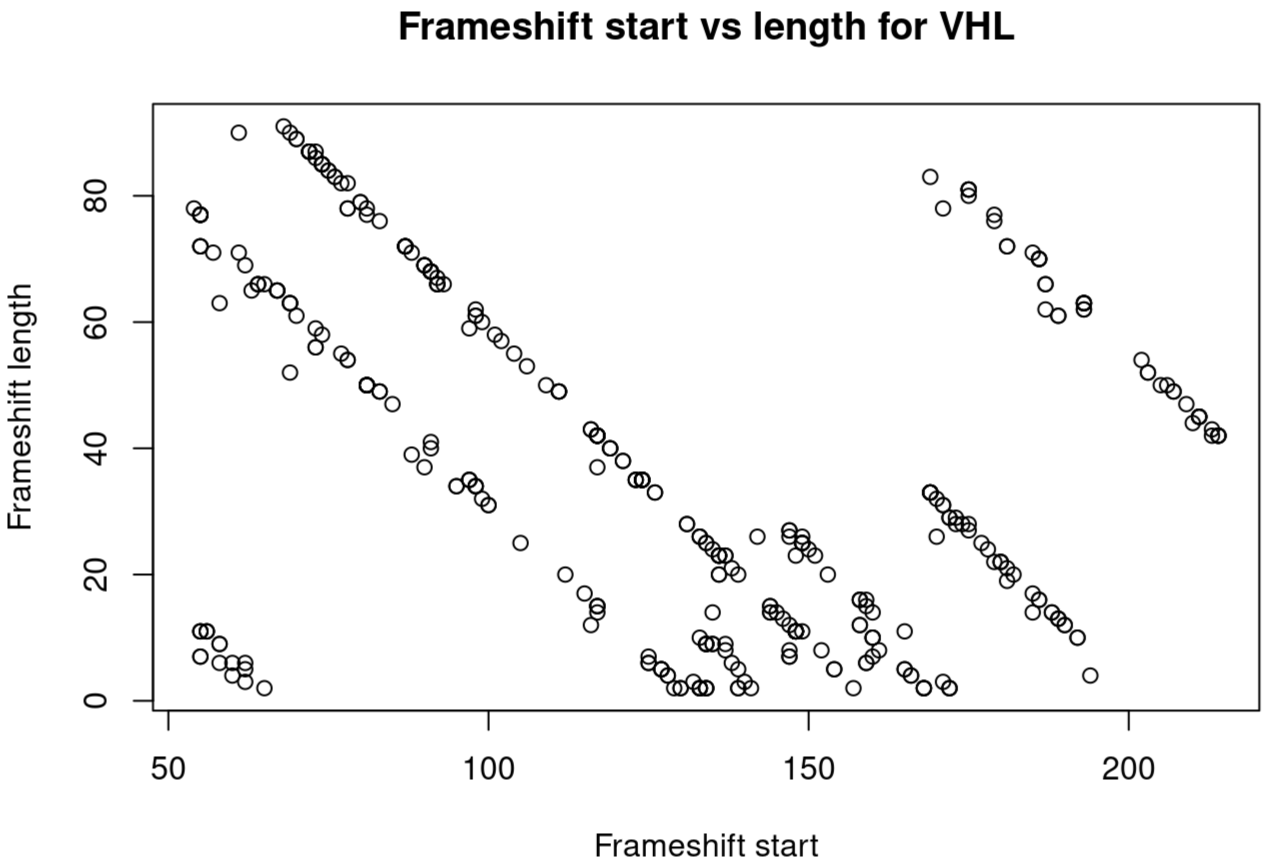
Frameshift start vs length for VHL
Frameshift length
Frameshift start
0
20
40
60
80
50
100
150
200

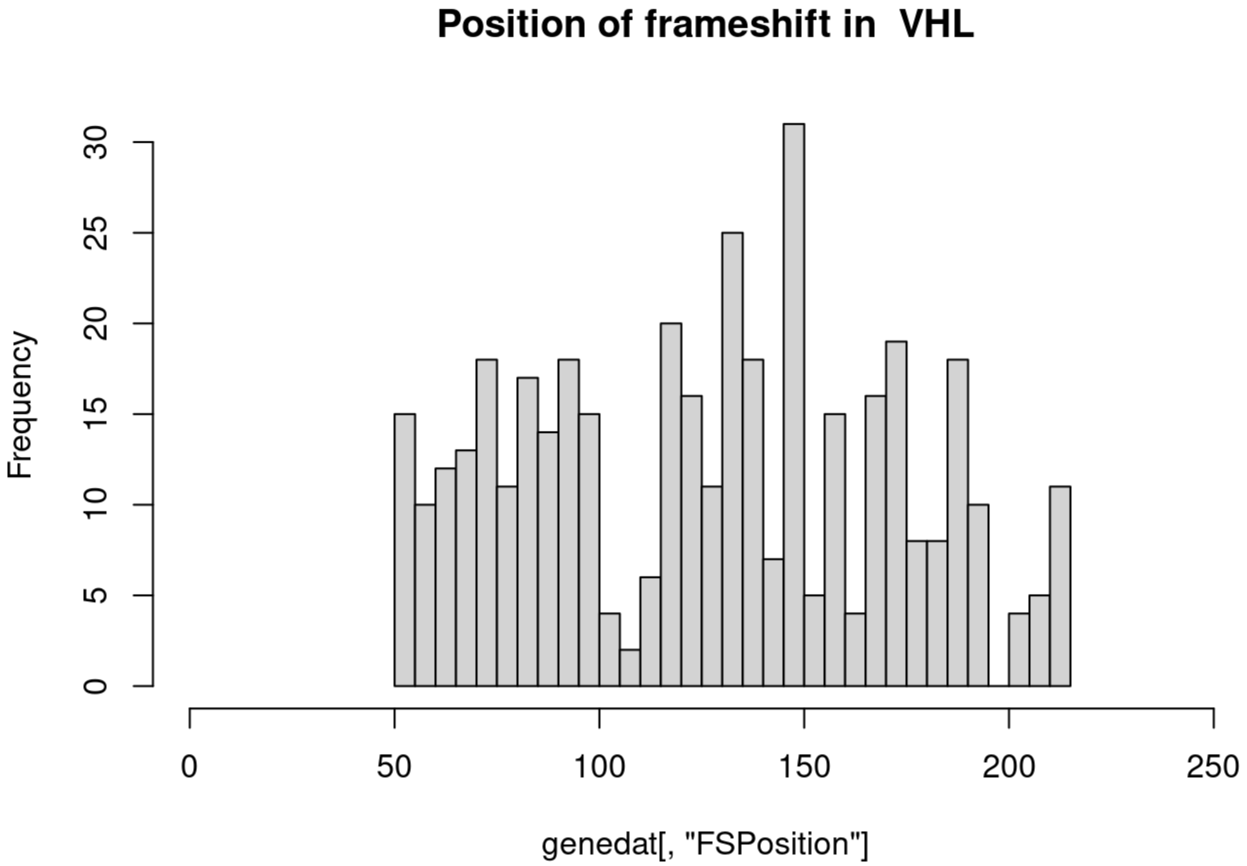
Position of frameshift in VHL
Frequency
genedat[, "FSPosition"]
0
5
10
15
20
25
30
0
50
100
150
200
250

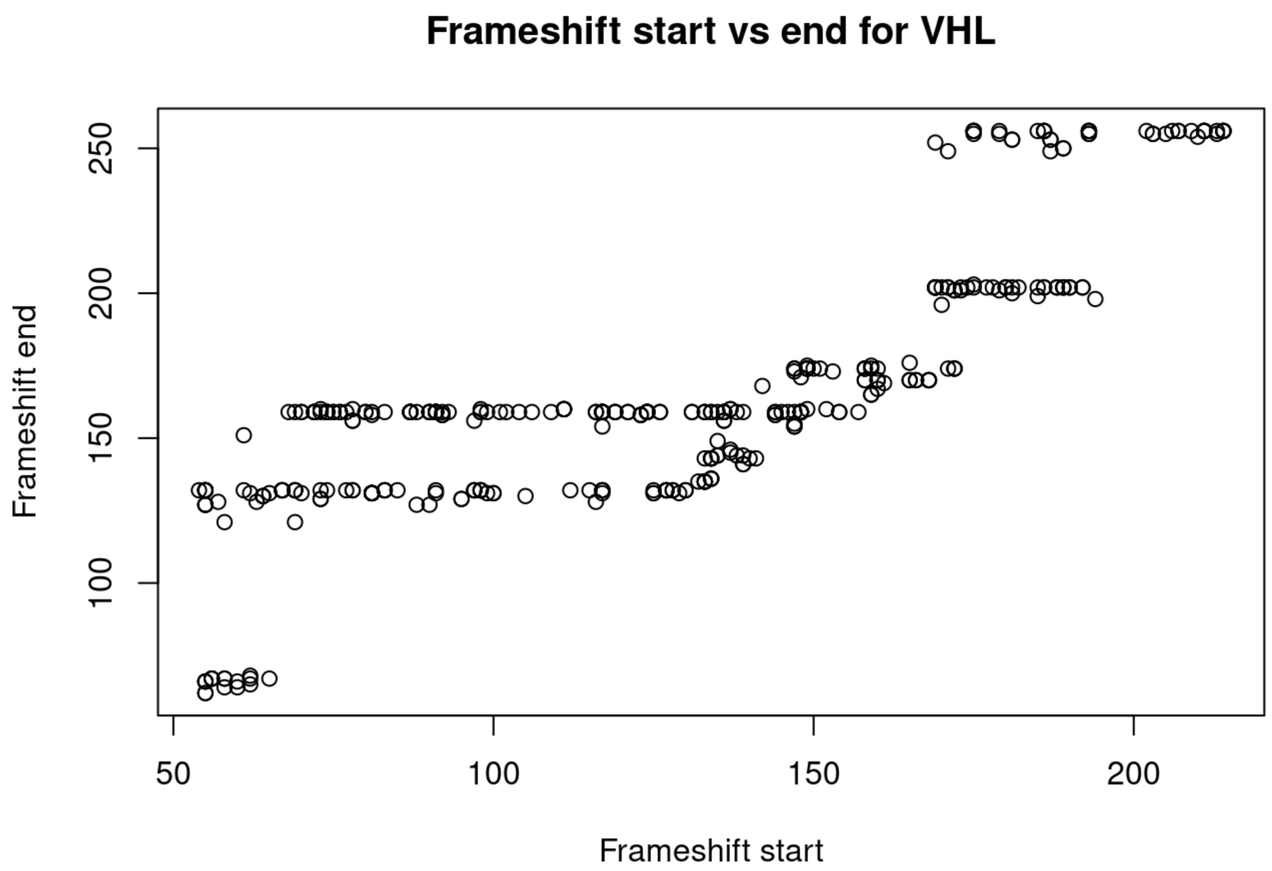
Frameshift start vs end for VHL
Frameshift end
Frameshift start
50
100
150
200
250

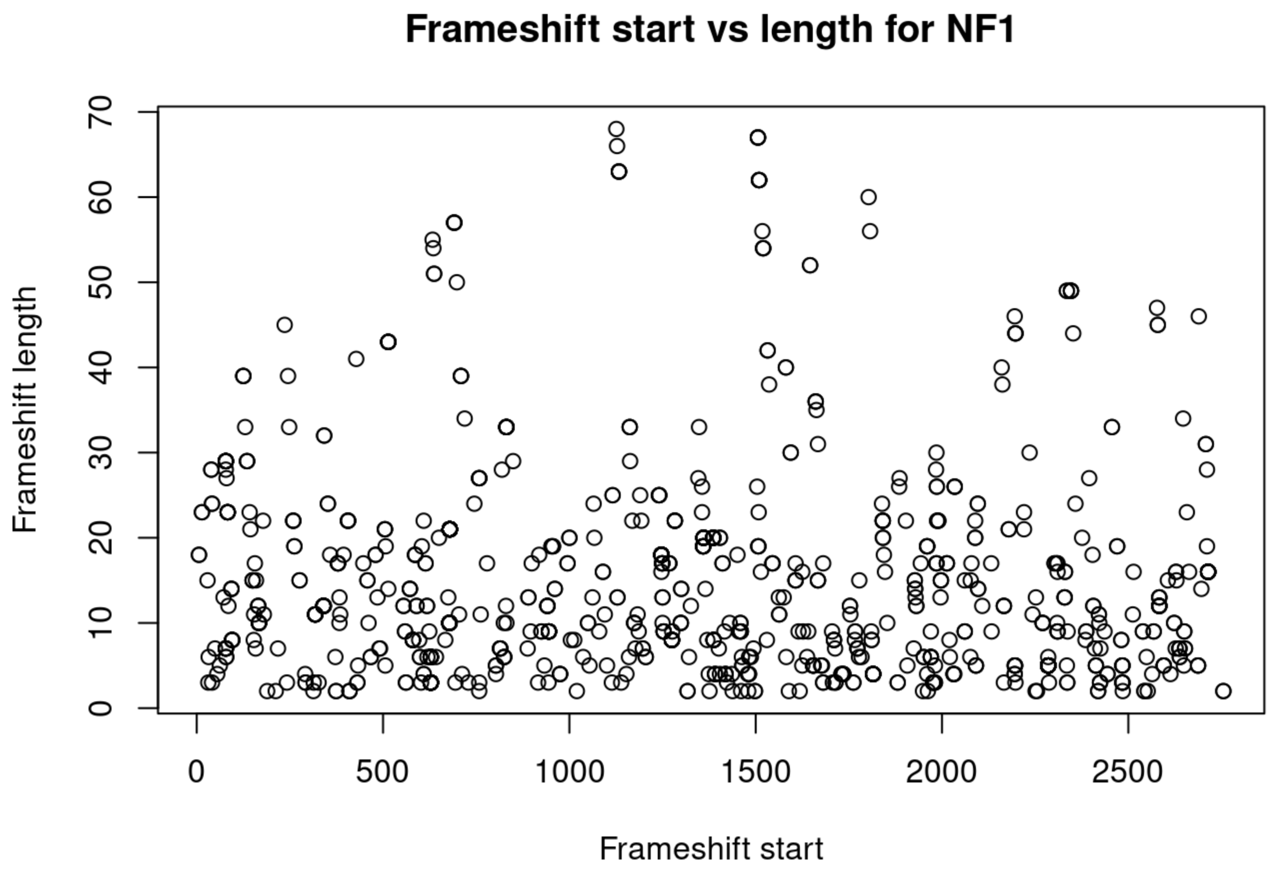
Frameshift start vs length for NF1
Frameshift length
Frameshift start
0
10
20
30
40
50
60
70
500
1000
1500
2000
2500

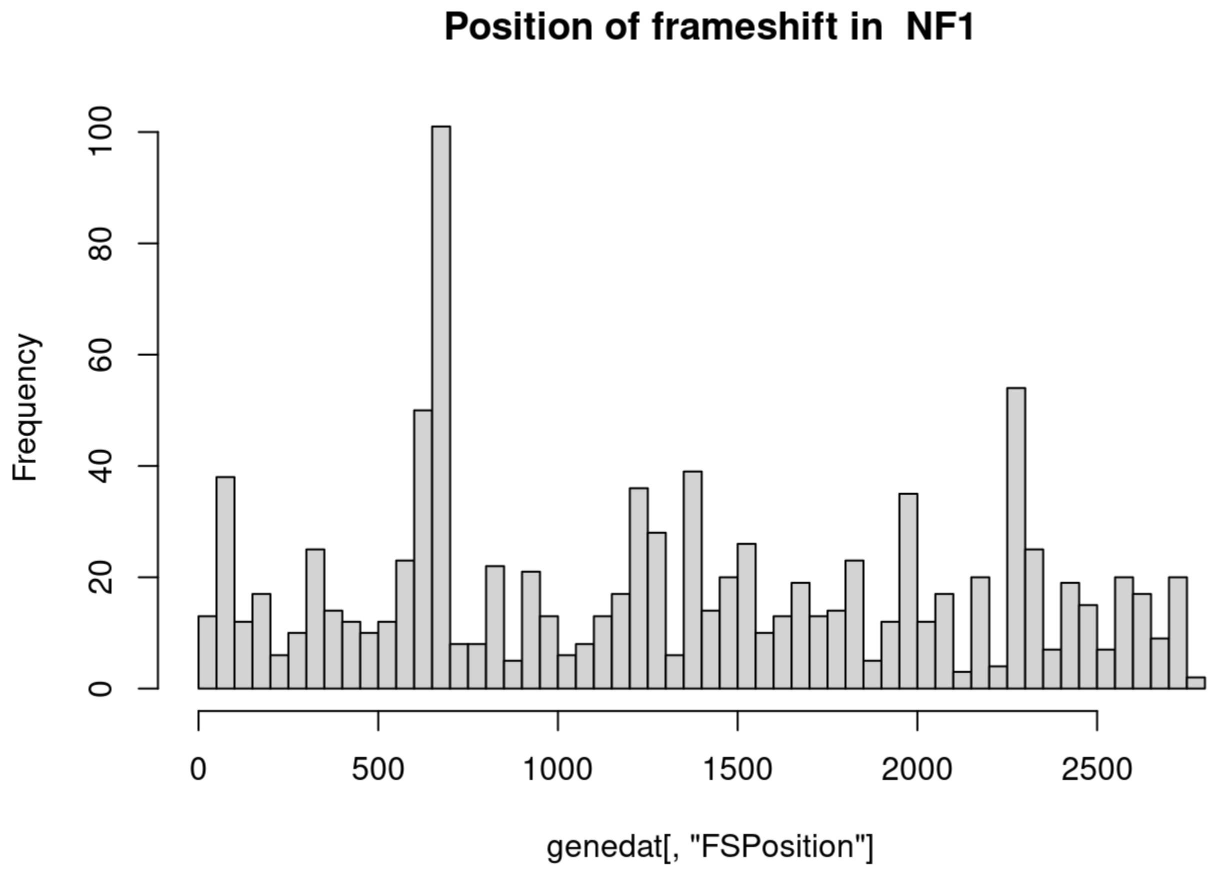
Position of frameshift in NF1
Frequency
0
20
40
60
80
100
0
500
1000
1500
2000
2500
genedat[, "FSPosition"]

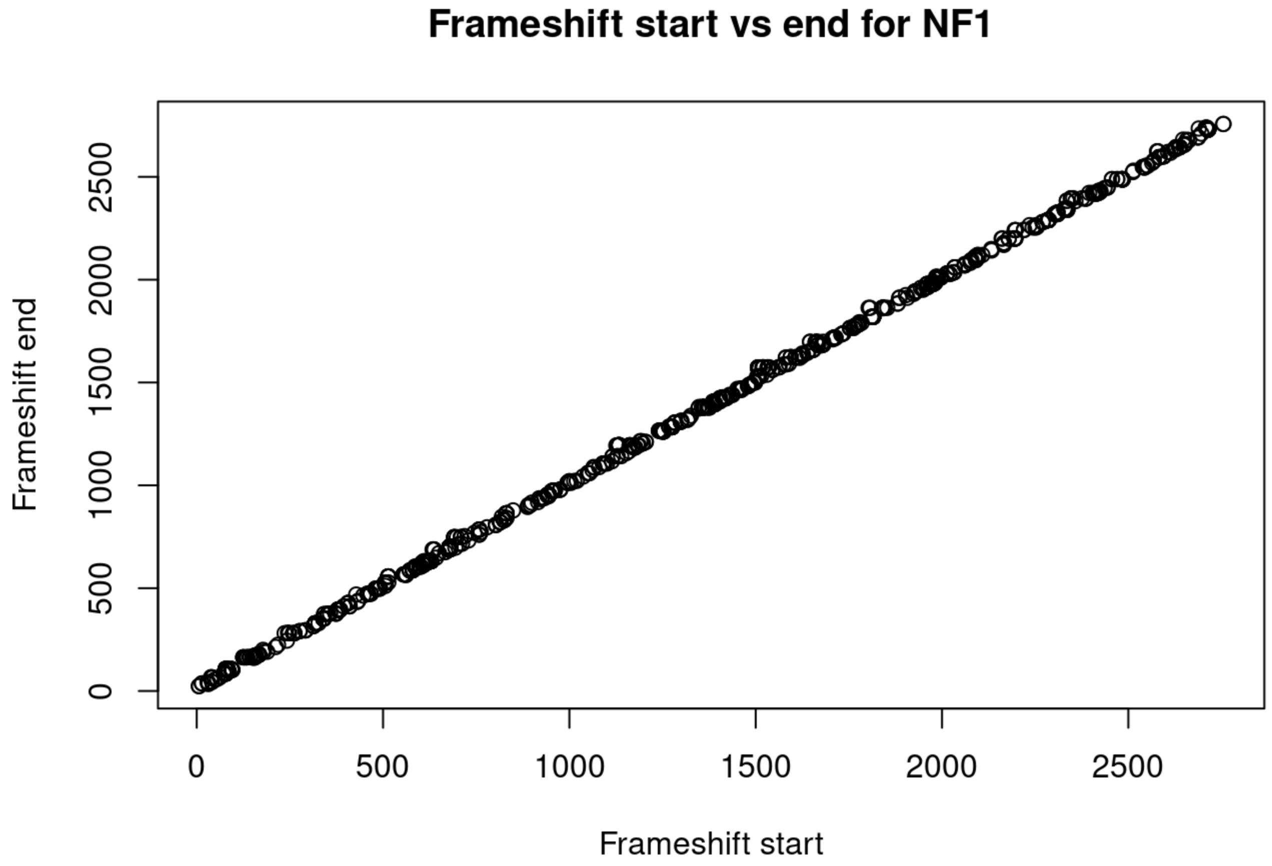
Frameshift start vs end for NF1
Frameshift end
0
500
1000
1500
2000
2500
Frameshift start

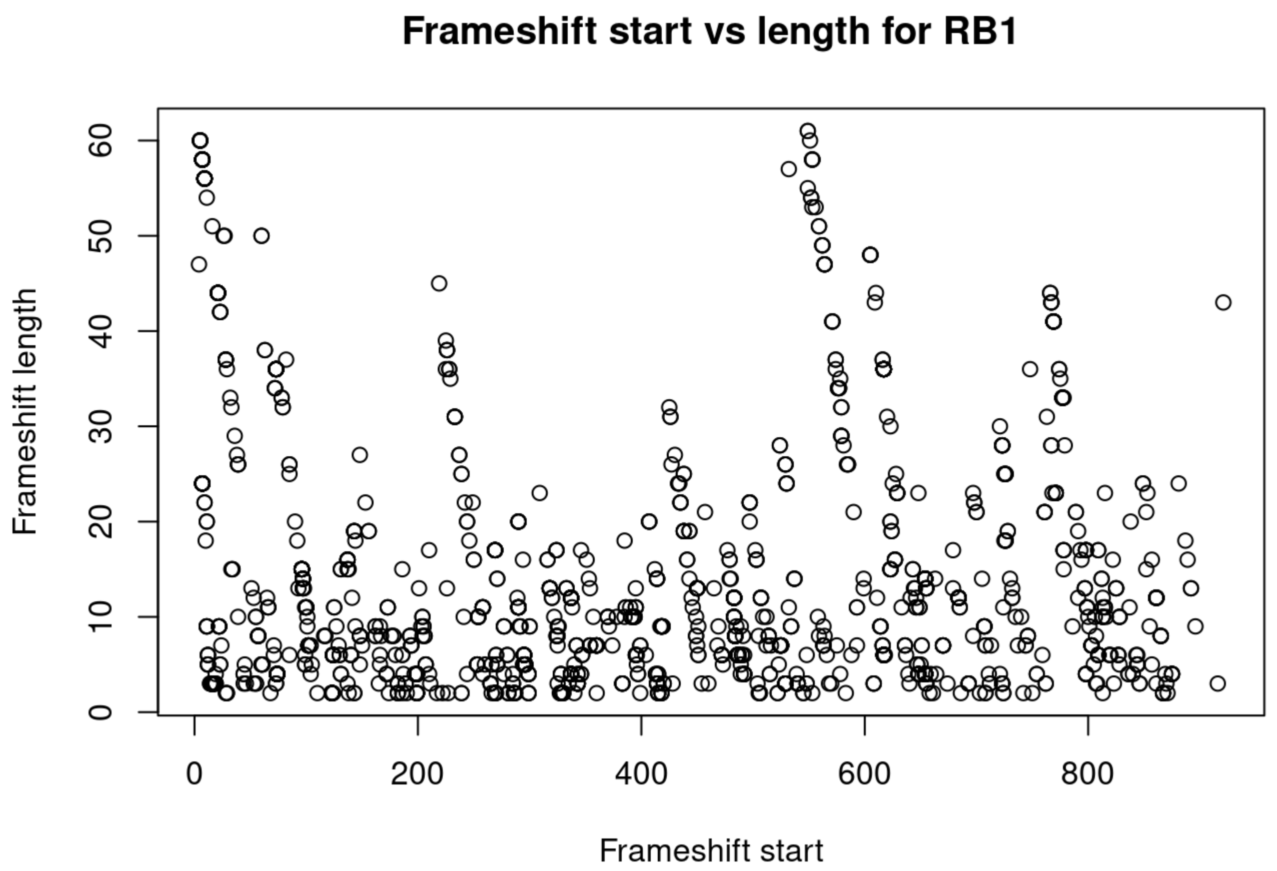
Frameshift start vs length for RB1
Frameshift length
Frameshift start
0
10
20
30
40
50
60
0
200
400
600
800

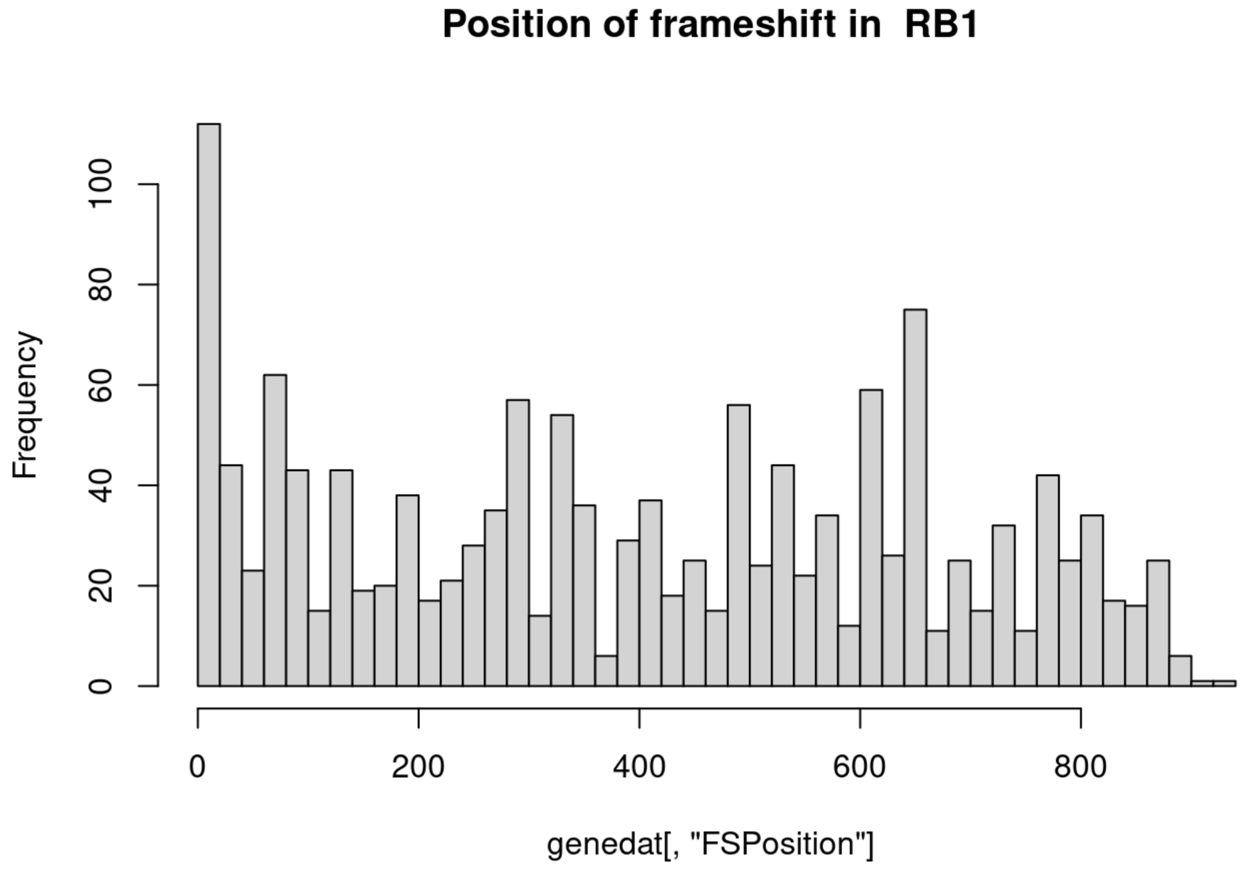
Position of frameshift in RB1
Frequency
genedat[, "FSPosition"]
0
20
40
60
80
100
0
200
400
600
800

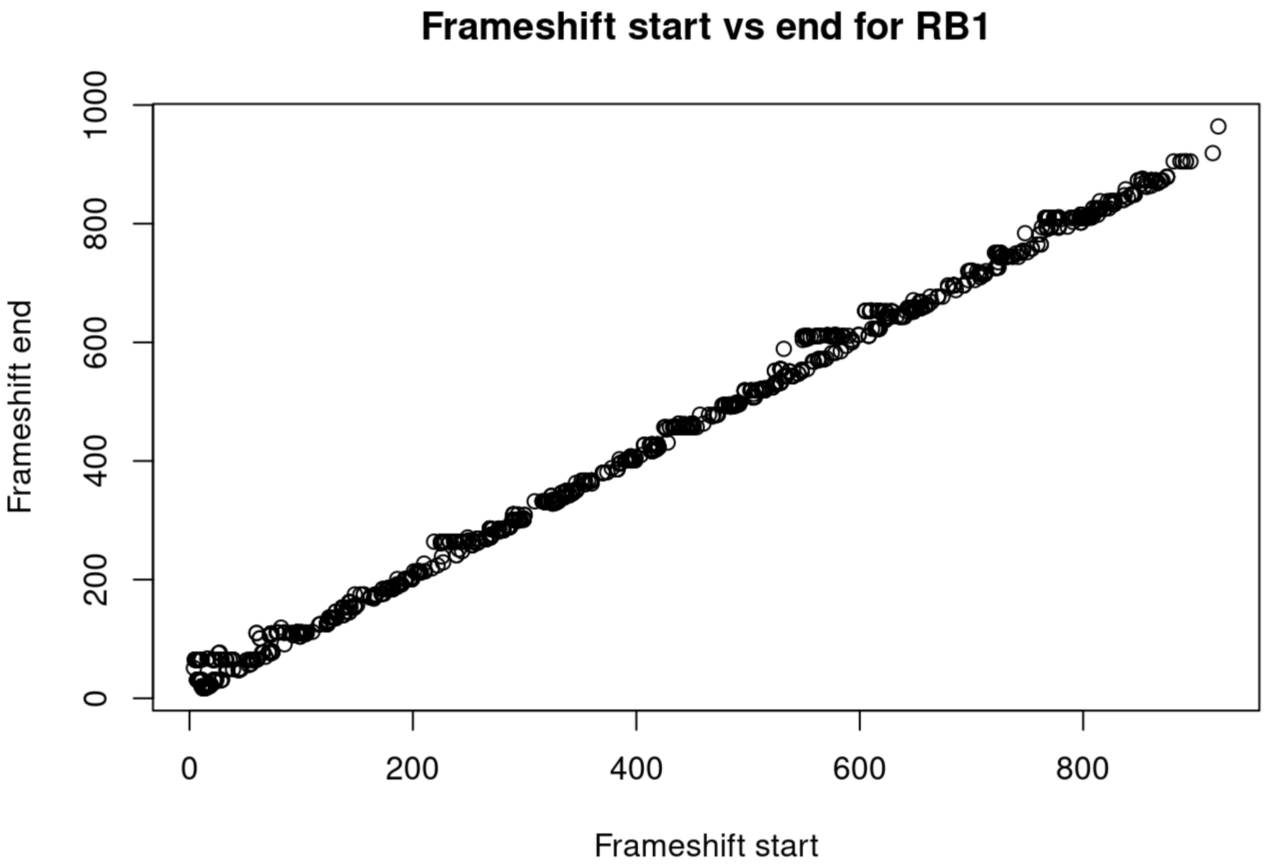
Frameshift start vs end for RB1
Frameshift end
Frameshift start
0
200
400
600
800
1000

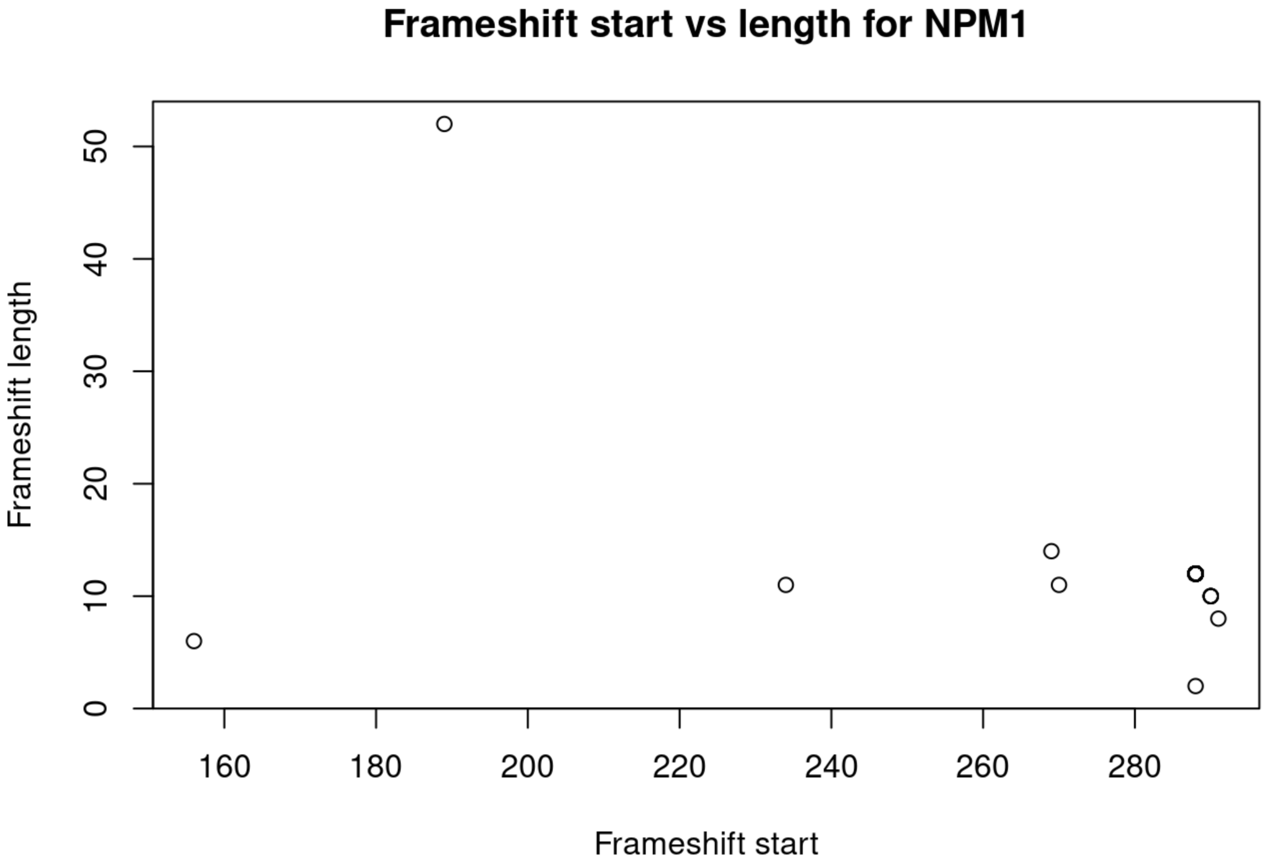
Frameshift start vs length for NPM1
Frameshift length
Frameshift start
0
10
20
30
40
50
160
180
200
220
240
260
280

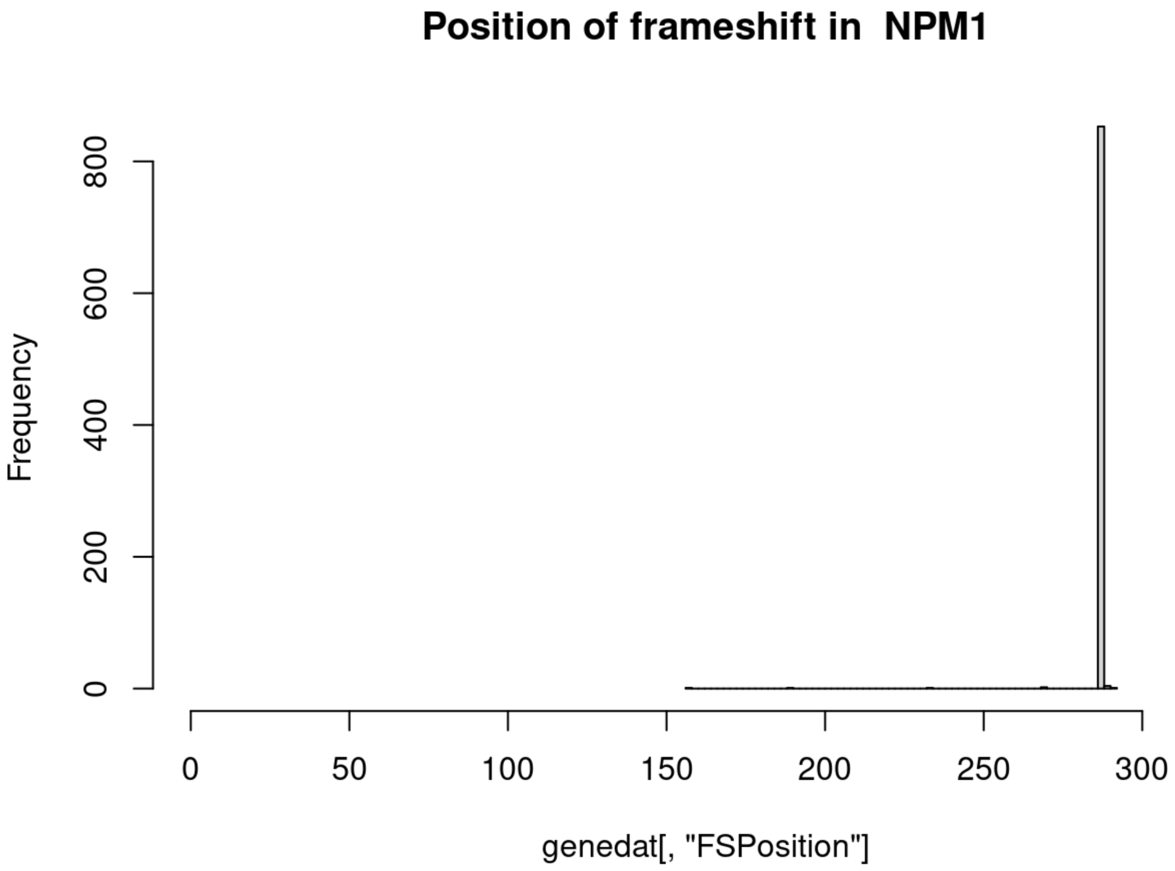
Position of frameshift in NPM1
Frequency
0
200
400
600
800
0
50
100
150
200
250
300
genedat[, "FSPosition"]

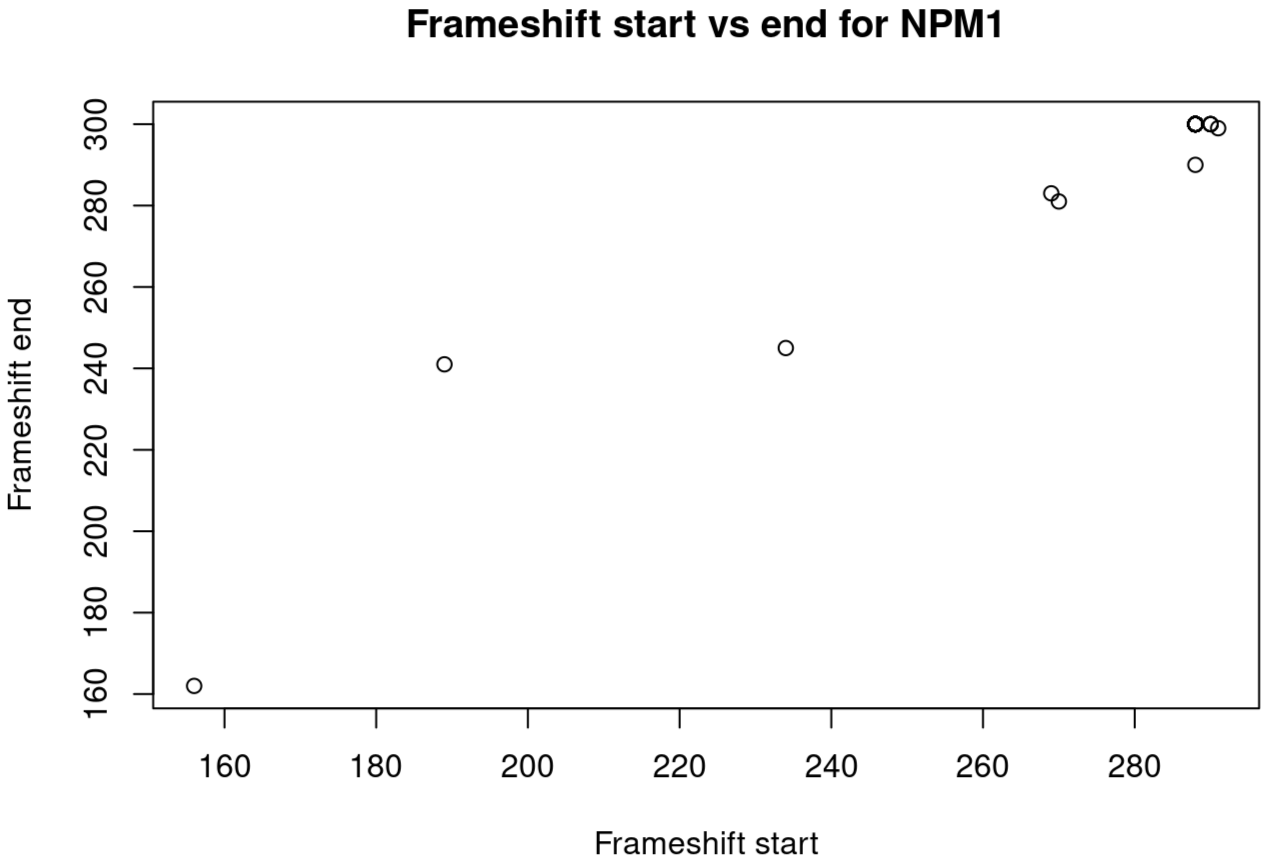
Frameshift start vs end for NPM1
Frameshift end
160
180
200
220
240
260
280
300
160
180
200
220
240
260
280
Frameshift start

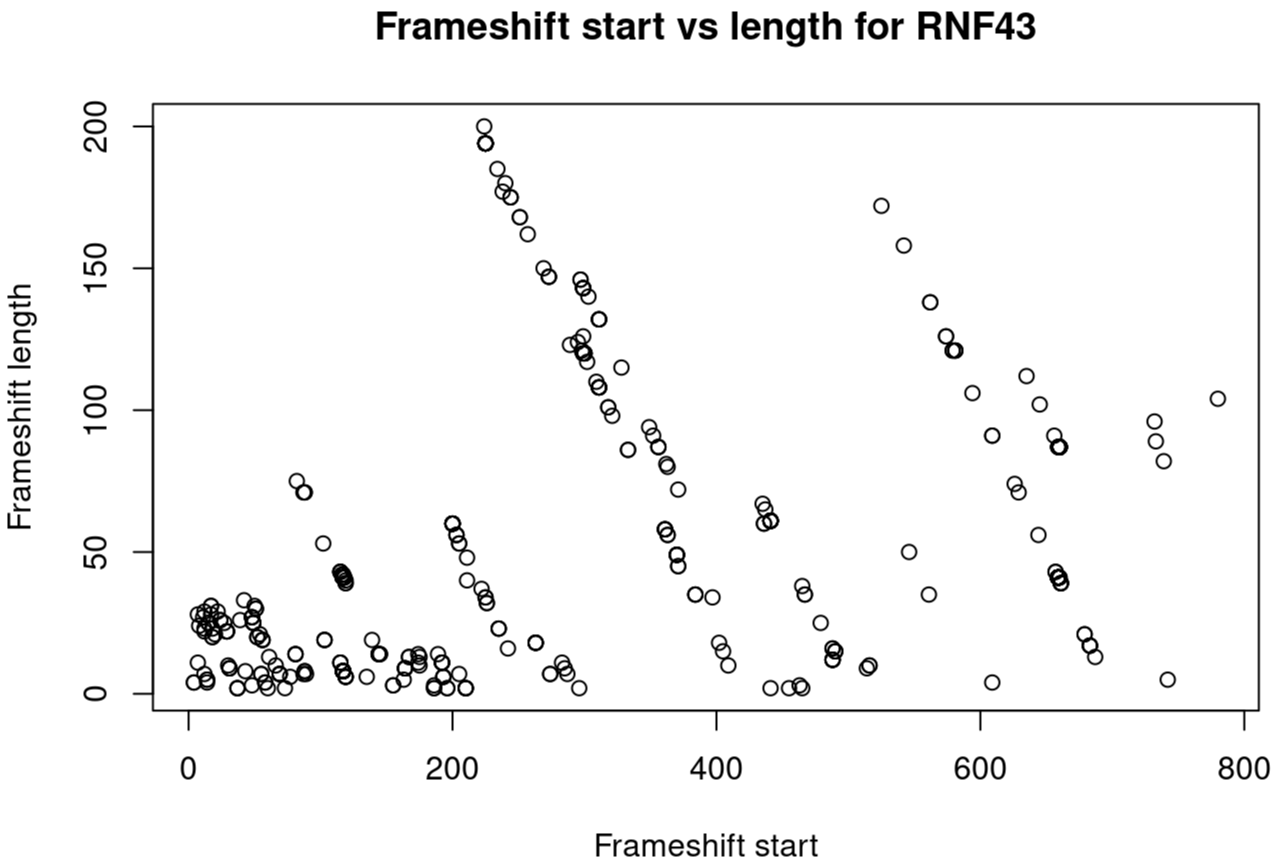
Frameshift start vs length for RNF43
Frameshift length
Frameshift start
0
50
100
150
200
0
200
400
600
800

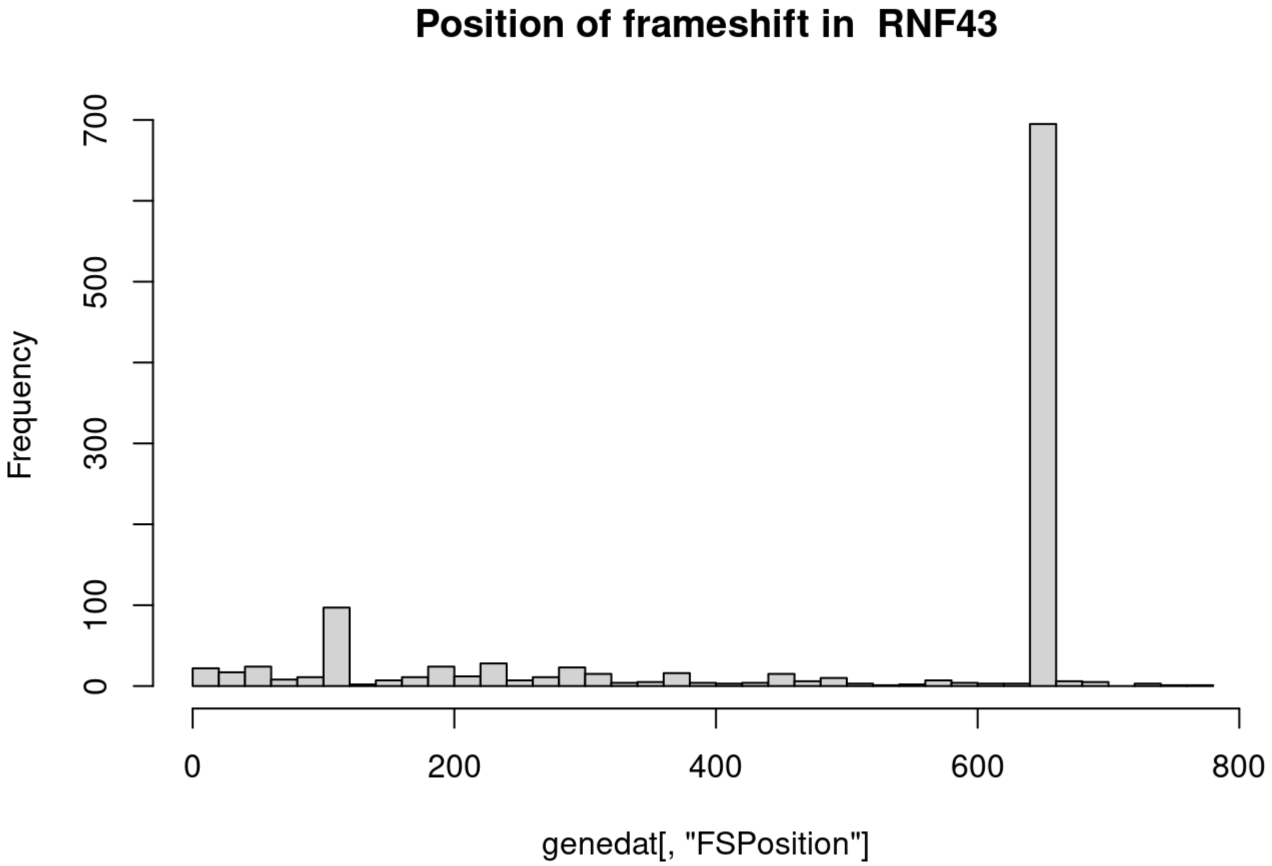
Position of frameshift in RNF43
Frequency
genedat[, "FSPosition"]
0
100
300
500
700
0
200
400
600
800

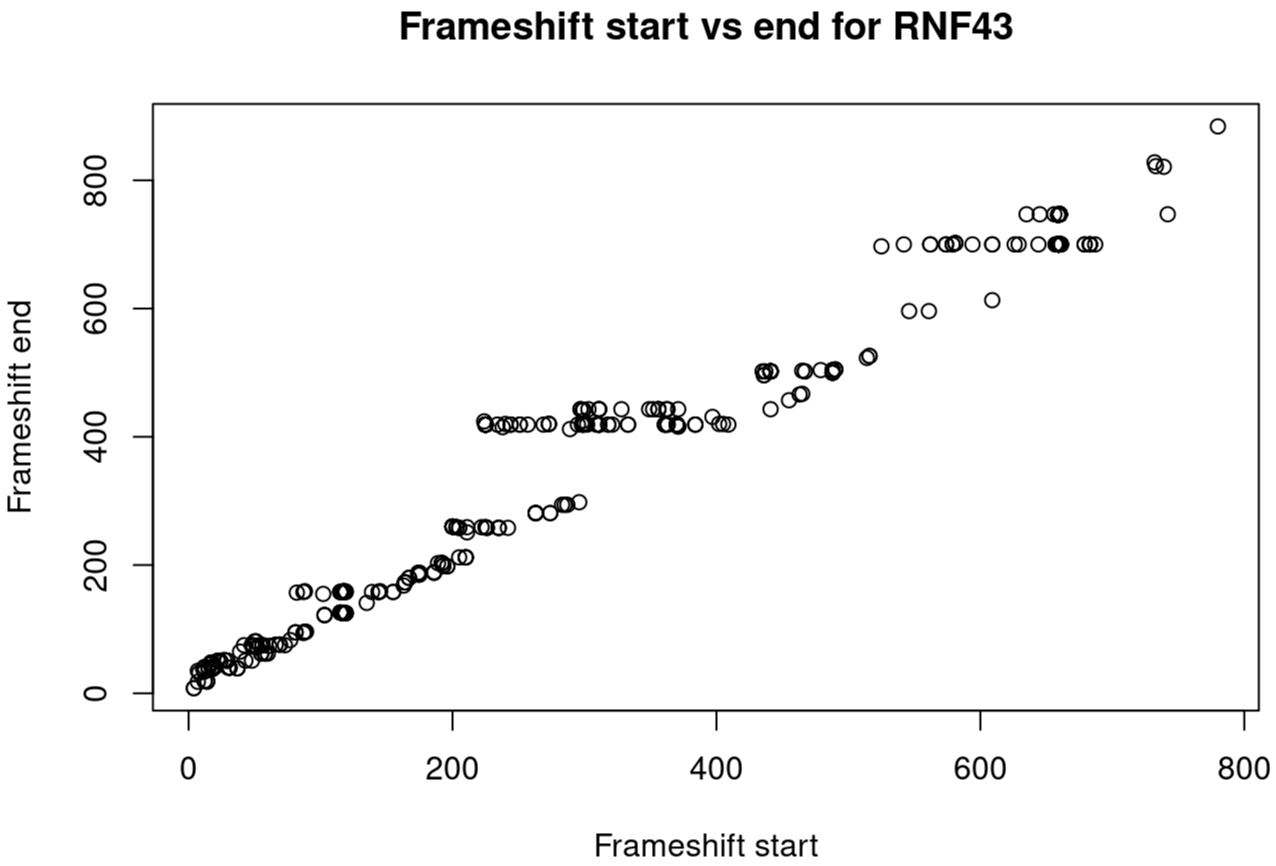
Frameshift start vs end for RNF43
Frameshift end
0
200
400
600
800
0
200
400
600
800
Frameshift start

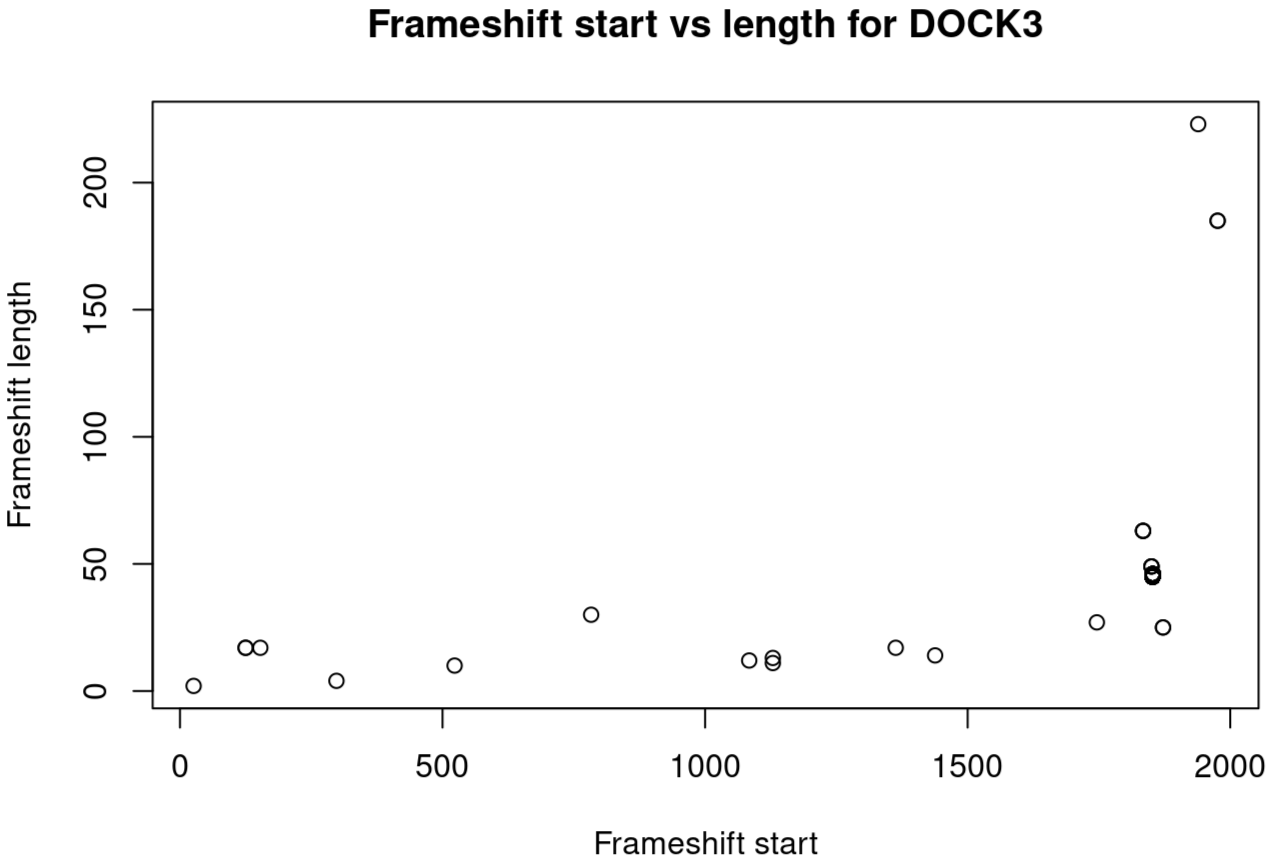
Frameshift start vs length for DOCK3
Frameshift length
0
50
100
150
200
0
500
1000
1500
2000
Frameshift start

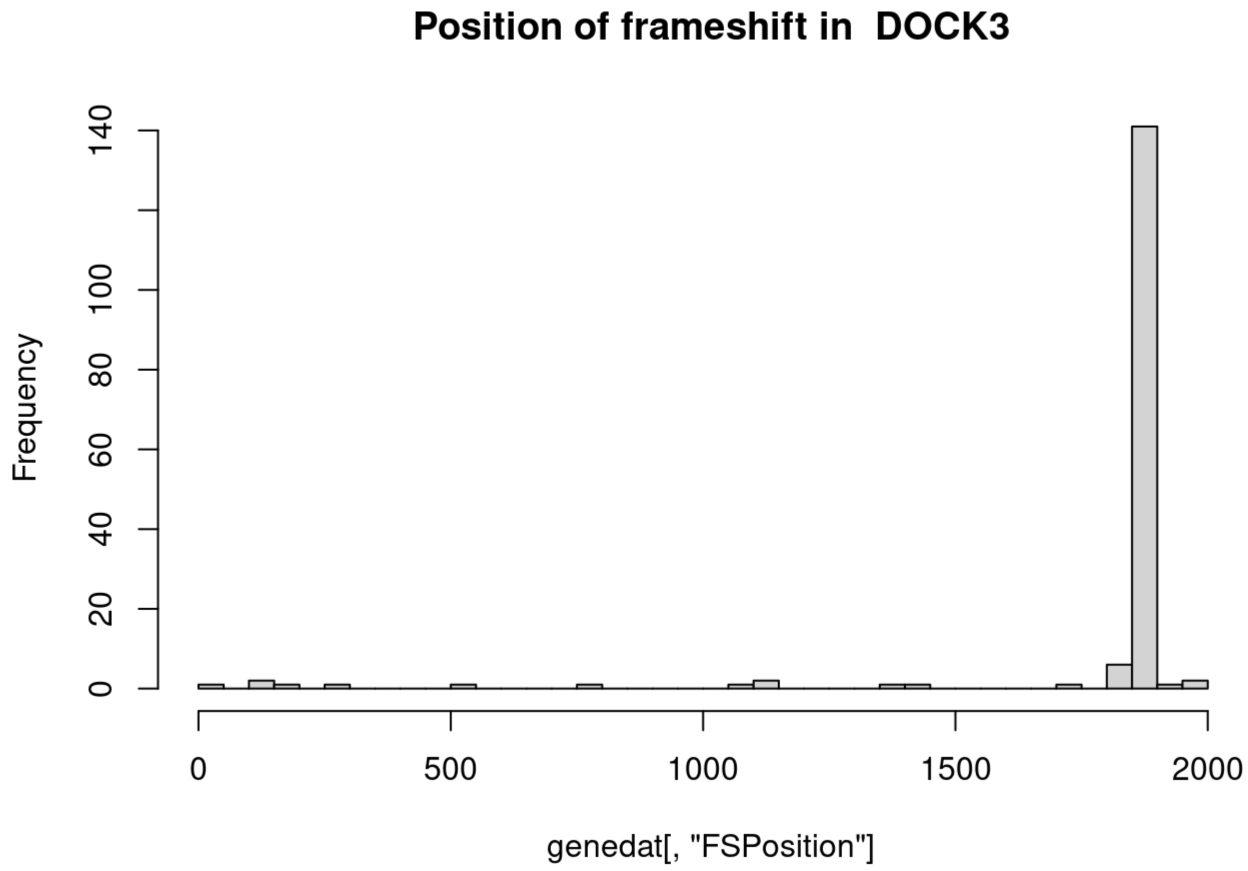
Position of frameshift in DOCK3
Frequency
0
20
40
60
80
100
140
0
500
1000
1500
2000
genedat[, "FSPosition"]

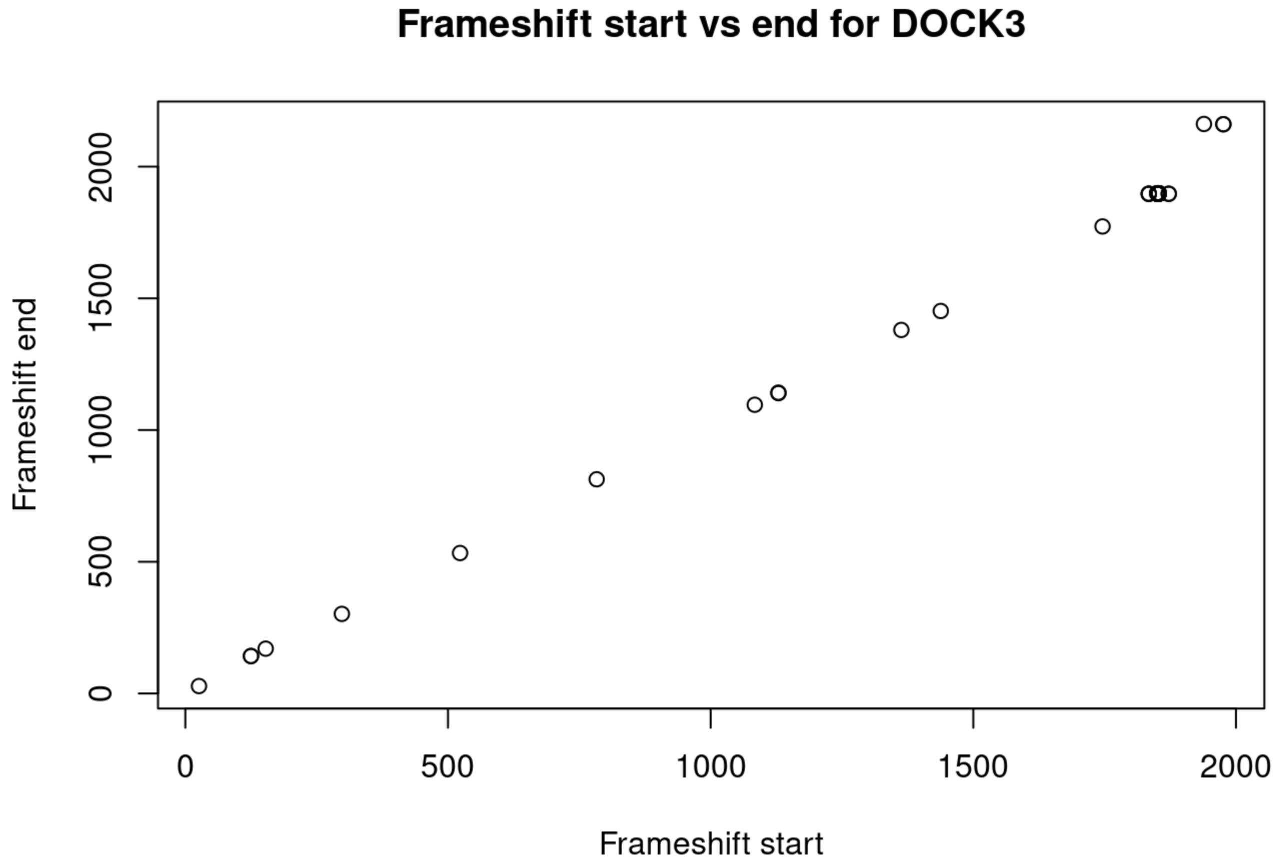
Frameshift start vs end for DOCK3
Frameshift end
0
500
1000
1500
2000
0
500
1000
1500
2000
Frameshift start

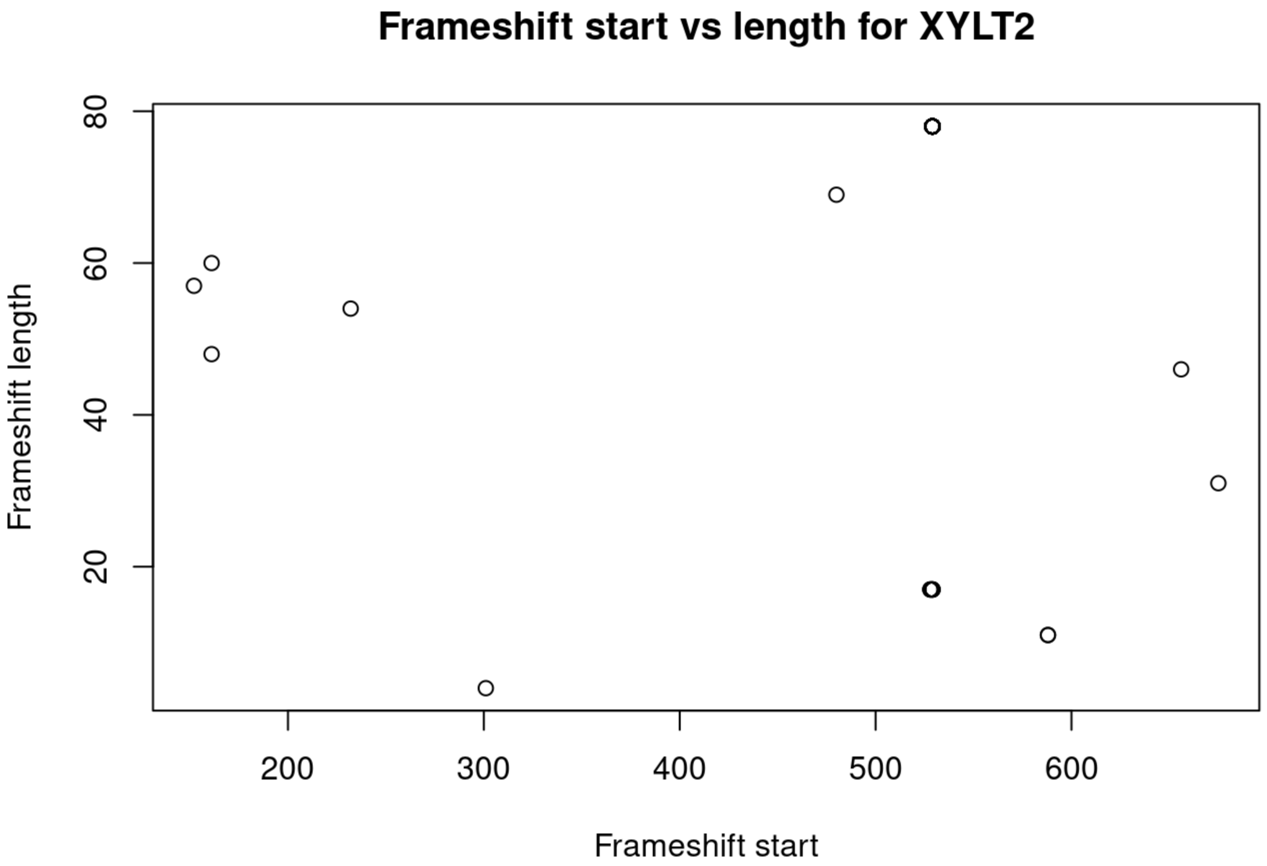
Frameshift start vs length for XYLT2
Frameshift length
Frameshift start
20
40
60
80
200
300
400
500
600

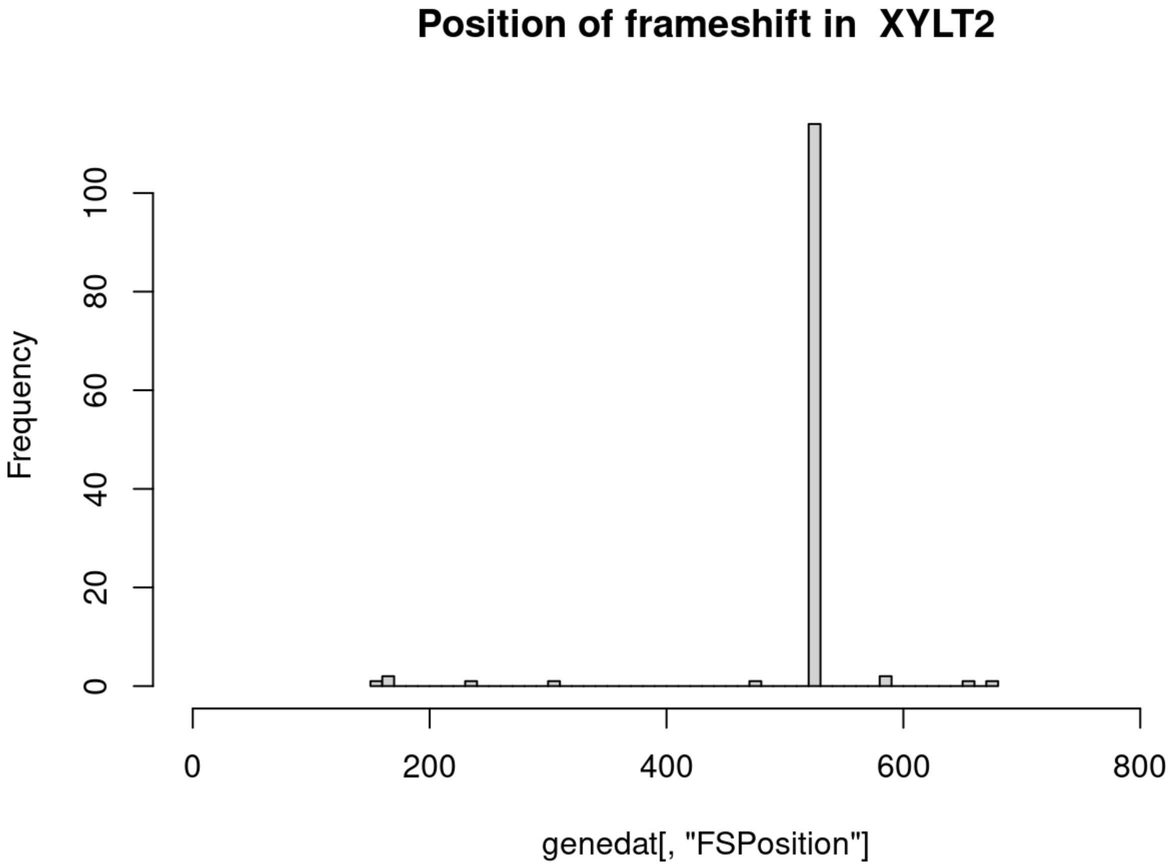
Position of frameshift in XYLT2
Frequency
genedat[, "FSPosition"]
0
20
40
60
80
100
0
200
400
600
800

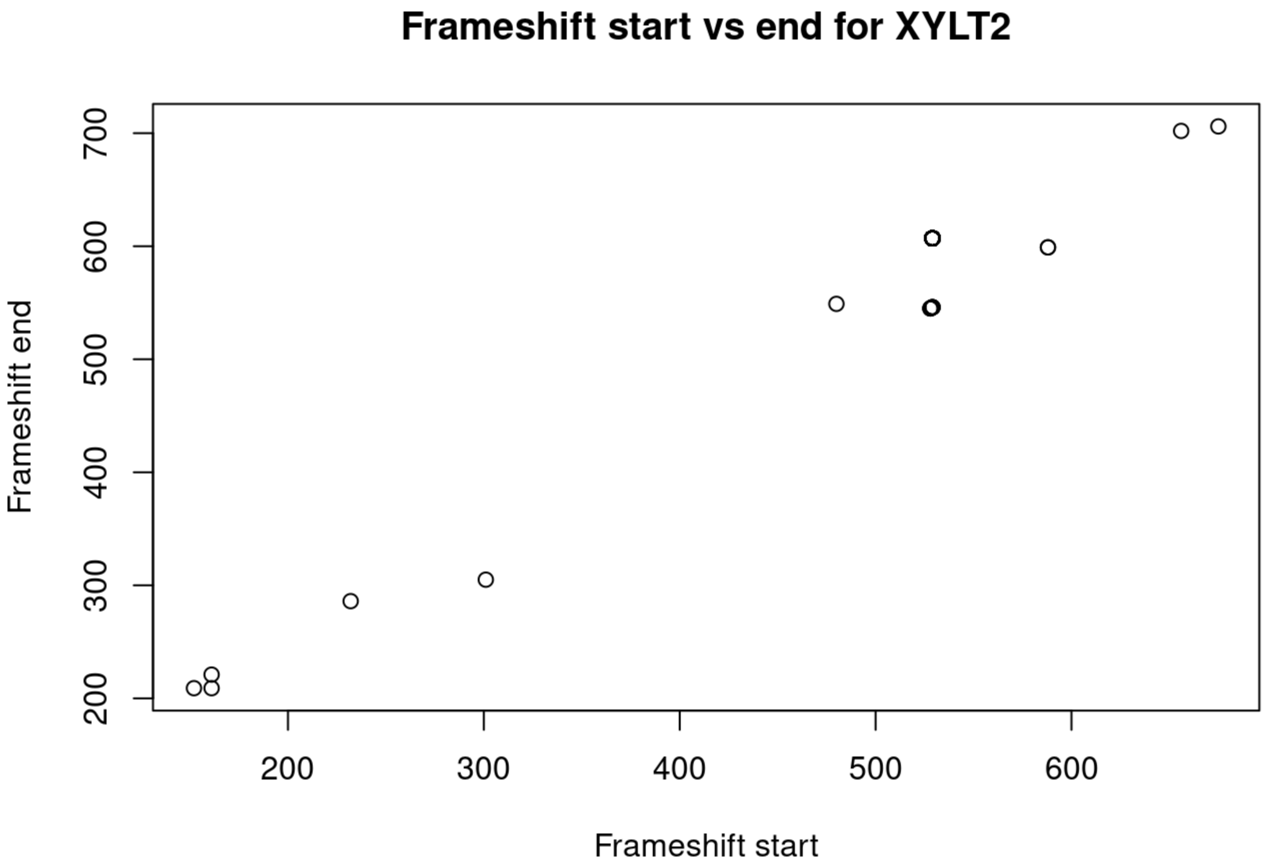
Frameshift start vs end for XYLT2
Frameshift end
200
300
400
500
600
700
200
300
400
500
600
Frameshift start

```
## [[1]]
## NULL
##
## [[2]]
## NULL
##
## [[3]]
## NULL
##
## [[4]]
## NULL
##
## [[5]]
## NULL
##
## [[6]]
## NULL
##
## [[7]]
## NULL
##
## [[8]]
## NULL
##
## [[9]]
## NULL
##
## [[10]]
## NULL
##
## [[11]]
## NULL
##
## [[12]]
## NULL
##
## [[13]]
## NULL
##
## [[14]]
## NULL
##
## [[15]]
## NULL
##
## [[16]]
## NULL
##
## [[17]]
## NULL
##
```

```
## [[18]]
## NULL
##
## [[19]]
## NULL
##
## [[20]]
## NULL
##
## [[21]]
## NULL
```

### Gene sites stats
